# The *Physcomitrium (Physcomitrella) patens* PpKAI2L receptors for strigolactones and related compounds highlight MAX2 dependent and independent pathways

**DOI:** 10.1101/2020.11.24.395954

**Authors:** Mauricio Lopez-Obando, Ambre Guillory, François-Didier Boyer, David Cornu, Beate Hoffmann, Philippe Le Bris, Jean-Bernard Pouvreau, Philippe Delavault, Catherine Rameau, Alexandre de Saint Germain, Sandrine Bonhomme

## Abstract

In flowering plants, the α/β hydrolase DWARF14 (D14) perceives strigolactone (SL) hormones and interacts with the F-box protein MORE AXILLARY GROWTH2 (MAX2) to regulate developmental processes. The key SL biosynthetic enzyme, CAROTENOID CLEAVAGE DEOXYGENASE8 (CCD8), is present in the moss *Physcomitrium (Physcomitrella) patens,* and PpCCD8-derived compounds regulate plant extension. Based on germination assays using seeds of the parasitic plant *Phelipanche ramosa,* we propose that these compounds are non-canonical SLs. Perception of PpCCD8-derived compounds does not require the PpMAX2 homolog. Candidate receptors are among the 13 *PpKAI2LIKE-A to -L* genes, homologous to the ancestral *D14* paralog *KARRIKIN INSENSITIVE2 (KAI2).* In Arabidopsis, AtKAI2 is the receptor for a still elusive endogenous KAI2-Ligand (KL). We show that in *P. patens,* among SL analogs, the (+)-GR24 enantiomer is a good mimic for PpCCD8-derived compounds, while the effects of the (-)-GR24 enantiomer, a KL mimic in flowering plants, are opposite. Interaction and binding assays of seven PpKAI2L proteins using pure enantiomers pinpoint at stereoselectivity towards (-)-GR24 for the (A-E) clade. Enzyme assays highlight strong hydrolytic activity of the PpKAI2L-H protein. Moss mutants for all *PpKAI2L* gene subclades were obtained and tested for their response to both enantiomers. We show that *PpKAI2L-A* to *-E* genes are not involved in PpCCD8-derived compound perception, but act in a PpMAX2-dependant pathway. In contrast, mutations in *PpKAI2L-G*, and *-J* genes abolish the response to (+)-GR24, suggesting that encoded proteins are receptors for PpCCD8-derived SLs.

## INTRODUCTION

Strigolactones (SLs) are butenolide compounds with dual roles in plants: exuded in soil, SLs signal the presence of a host to Arbuscular Mycorrhizal (AM) fungi (Akiyama et al. 2005; Besserer et al. 2006), and thus favor the establishment of symbiosis; as endogenous compouds, they (or derived compounds) play a hormonal role in developmental programs (Gomez-Roldan et al. 2008; Umehara et al. 2008) for reviews: (Lopez-Obando et al. 2015; Waters et al. 2017). SLs exuded from plant roots also act as signalling molecules in the rhizosphere inducing parasitic plant seed germination (Cook et al. 1966) for review: (Delavault 2017). SLs have been found in most land plants, including bryophytes, lycophytes, gymnosperms, and angiosperms (Yoneyama et al. 2018a). However, their synthesis and signaling pathways are mainly described in angiosperms where core enzyme pathways, among which two CAROTENOID CLEAVAGE DEOXYGENASEs (CCD7 and CCD8), convert carotenoids into carlactone (CL). CL is the reported precursor of all known SLs, and the substrate for further enzymes such as the CYTOCHROME-P450 MORE AXILLARY GROWTH1 (MAX1) (for review: (Alder et al. 2012; Al-Babili and Bouwmeester 2015). Depending on the plant species, CL is converted into canonical or non-canonical SLs. These differ in the structure attached to the conserved enol ether-D ring moiety, shared by all SLs, and essential for biological activity (Yoneyama et al. 2018a; Yoneyama 2020). In angiosperms, SLs are perceived by an α/β hydrolase DWARF14 (D14)/DECREASED APICAL DOMINANCE2 (DAD2)/ RAMOSUS 3 (RMS3) (Arite et al. 2009; Hamiaux et al. 2012; de Saint Germain et al. 2016) that interacts with an F-box protein MORE AXILLARY GROWTH2 (MAX2) to target SUPPRESSOR OF MAX2-LIKE (SMXL) repressor proteins for proteasome degradation (Soundappan et al. 2015; Waters et al. 2015). The originality of SL perception is that the D14 protein is both a receptor and an enzyme that cleaves its substrate (and covalently binds part of the SL), in a signaling mechanism that is still under debate (Yao et al. 2016; de Saint Germain et al. 2016; Shabek et al. 2018; Seto et al. 2019). In all cases, the pocket of the α/β hydrolase appears essential for substrate/ligand (SL) interaction (for review:(Bürger and Chory 2020)).

The evolutionary origins of SLs, and in particular whether their primary role is that of a hormone or rhizospheric signal are still unclear. SL identification and quantification is a challenge in many species, due to the very low amounts of the molecules present in plant tissues or exudates, and their high structural diversification (Xie 2016; Yoneyama et al. 2018b). Therefore, the occurrence of SLs in a species was often inferred from the presence of the core biosynthesis enzymes in its genome (Delaux et al. 2012; Walker et al. 2019). Recently SLs were proposed to only be produced in land plants (Walker et al. 2019). Evidence of an ancestral signaling pathway to SL pathway came from the identification of an ancient D14 paralog named KARRIKIN INSENSITIVE2/HYPOSENSITIVE TO LIGHT (KAI2/HTL) during screening of Arabidopsis mutants (Nelson et al. 2011). KAI2 is also an α/β hydrolase like D14, that also interacts with the MAX2 F-box protein, in a pathway regulating Arabidopsis seed germination and seedling development (Nelson et al. 2011; Waters et al. 2012). However, the endogenous signal perceived by KAI2 remains unknown and is reported thus far as the KAI2-Ligand (KL) (Conn and Nelson 2015). KAI2 is also involved in stress tolerance, drought tolerance and AM symbiosis (Gutjahr et al. 2015; Wang et al. 2018; Villaecija-Aguilar et al. 2019; Li et al. 2020).

To gain insight into SL signaling evolution, we focused our studies on a model for non-vascular plants, namely *Physcomitrium (Physcomitrella) patens* (*P. patens*). As a moss, *P. patens* belongs to the bryophytes, which also include two other clades, hornworts and liverworts (Bowman et al. 2019). Bryophytes are currently described as a monophyletic group of embryophytes sharing an ancestor with vascular plants (Puttick et al. 2018; Harris et al. 2020). Therefore, comparing signaling pathways between extant vascular plants and extant bryophytes can provide an insight about the evolutionary origin of these pathways. Furthermore, studying extant bryophytes may give clues for understanding how the first plants have been able to survive out of water, and conquer land, 450 million years ago (Bowman et al. 2019; Blázquez et al. 2020; Harris et al. 2020). In *P. patens,* both CCD enzymes required for SL synthesis are found (PpCCD7 and PpCCD8) (Proust et al. 2011), and carlactone has been detected as the product of PpCCD8 (Decker et al. 2017). The extended phenotype of *Ppccd8* mutant plants indicates that PpCCD8-derived molecules are required for moss filament growth regulation. Moreover, these molecules also act as a growth limiting signal between neighbouring moss plants, as they are exsuded in the medium (Proust et al. 2011). PpCCD8-derived molecules also appear to play a role in rhizoid elongation and gametophore shoot branching (Delaux et al. 2012; Coudert et al. 2015). Application of the artificial SL, (±)-GR24, complements the *Ppccd8* mutant phenotype, suggesting that PpCCD8-derived molecules are indeed SL-like compounds (Proust et al. 2011). However, the exact nature of PpCCD8-derived molecules is still elusive (Yoneyama et al. 2018b) and the absence of *MAX1* homologs in *P. patens* suggests that the synthesis pathway in this moss may differ from that of vascular plants. Nevertheless, phylogenetic analysis of *MAX1* homologs highlights the presence of this gene in other mosses and suggests that the biosynthesis pathway is otherwise conserved in land plants (Walker et al. 2019). SL signaling also seems to differ between angiosperms and *P. patens.* Indeed, contrary to its flowering plant homolog, PpMAX2 is likely not involved in the response to PpCCD8-derived molecules, as the corresponding mutant does respond to (±)-GR24 (Lopez-Obando et al. 2018). The PpMAX2 F-box protein appears to be involved in a light dependent pathway required for moss early development and regulation of gametophores number and size (Lopez-Obando et al. 2018). No true homolog for the D14 SL receptor is found in the *P. patens* genome, whereas 11-13 *PpKAI2-LIKE (PpKAI2L)* candidate genes, named *PpKAI2L-A* to *PpKAI2L-M,* were described (Lopez-Obando et al. 2016a). These genes are grouped into four subclades, hereafter renamed as PpKAI2L-(A-E), (FK), (HIL), and (JGM) (see also below). A comprehensive phylogenetic assessment placed the *P. patens* clades (FK), (HIL), and (JGM) into a super clade called DDK (D14/DLK2/KAI2) containing spermatophyte (angiosperm and gymnosperm) D14 clades, while clade (A-E) belongs to the eu-KAI2 clade, highly conserved and common to all land plants (Bythell-Douglas et al. 2017). Nevertheless, moss proteins from the DDK clade were found to be as different from D14 as from KAI2 (Bythell-Douglas et al. 2017). Prediction of PpKAI2L protein structures found various pocket sizes, as observed for D14 and KAI2 from vascular plants (Lopez-Obando et al. 2016a). Larger pocket size was predicted for PpKAI2L-F and -K, and maybe also for PpKAI2L-J and -G, while smaller pocket was predicted for clade (A-E) proteins. Consequently, these proteins could be receptors with diverse substrate preferences, and might bind either PpCCD8-derived compounds or the elusive KL (Conn and Nelson 2015). Accordingly, in our previous study of PpMAX2, we hypothesized that this F-box protein might be involved in a putative *P. patens* KL signaling pathway (Lopez-Obando et al. 2018). But the question as to the involvement of PpKAI2L proteins in the PpMAX2 pathway remains open. Recently, the crystal structures of PpKAI2L-C, -E and -H were published (Bürger et al. 2019). It was shown that *in vitro* purified PpKAI2L proteins -C, -D and -E were destabilized by (-)-5-deoxystrigol, a canonical SL with non-natural stereochemistry, but binding affinity was not determined for the pure enantiomer. In contrast, PpKAI2L proteins -H, -K and -L could bind the karrikin KAR_1_ (Bürger et al. 2019). Proteins from the (JGM) clade were not studied, and no evidence for a role of one (or several) PpKAI2L as receptors for CCD8-derived molecules was provided, nor were experiments carried out in *P. patens* to validate the results. Moreover, the involvement of PpKAI2L proteins in the putative PpMAX2-dependent KL signaling pathway was not explored.

The aim of the present study was to investigate the nature of the PpCCD8-derived molecules in moss and identify the receptors for these compounds. These goals would be facilitated by using a mimic of these molecules in assays. So far, the racemic (±)-GR24 has been used as a SL analog, but recent reports in angiosperms found that the different enantiomers present in this synthetic mixture do not have the same effect (Scaffidi et al. 2014). Indeed, (+)-GR24, (also called GR24^5DS^) with a configuration close to the natural strigol is mostly perceived by D14 and mimics CCD8-derived SLs (e.g. carlactone). On the other hand, as for (-)-5-deoxystrigol, the configuration of (-)-GR24 (also called GR24^*ent*-5DS^) has so far not been found in natural SLs. However, KAI2 perceives the (-)-GR24 analog better than D14 proteins and (-)-GR24 has therefore been described as a KL mimic (Scaffidi et al. 2014; Zheng et al. 2020).

Here we tested the activity of *P. patens* as a stimulant for *P. ramosa* germination, in an attempt to shed light on the type of SLs derived from PpCCD8. Refining and supplementing the work of (Bürger et al. 2019), we fully characterized seven PpKAI2L proteins *in vitro* by testing their cleavage activity and binding to pure GR24 enantiomers. We showed that stereoselectivity of most of the PpKAI2L proteins for GR24 enantiomers is weak, except for the (A-E) clade showing preferrential affinity for (-)-GR24. We highlighted the strong and non selective enzyme activity of PpKAI2L-H. We expressed these proteins in the Arabidopsis *d14-1 kai2-2* double mutant to examine conservation of the SL and/or KL perception function. Finally, we used CRISPR-Cas9 technology to generate several *P. patens* multiple mutants, affected in all four *PpKAI2L* clades. By coupling analysis of these mutants' phenotype and response to pure GR24 enantiomers with our biochemistry results, we provide strong evidence that clade (A-E) PpKAI2L proteins could be moss KL receptors in a PpMAX2 dependent pathway, while clade (JGM) PpKAI2L would function as moss SL receptors.

## RESULTS

### PpCCD8-derived compounds induce the germination of a hemp-specific population of *Phelipanche ramosa*

A recent report (Yoneyama et al. 2018b) indicated that canonical SLs previously identified in *P. patens* tissues (Proust et al. 2011) could be contaminants. However other evidence suggests that *P. patens* does synthesize SL-like products derived from carlactone. Indeed, PpCCD8 shows carlactone synthase activity (Decker et al. 2017), and both the synthetic SL analog (±)-GR24 and carlactone do complement the *Ppccd8* phenotype (Proust et al. 2011; Decker et al. 2017). Still, quantification of SL and related compounds is a challenge in many species (Boutet-Mercey et al. 2018; Yoneyama et al. 2018b; Floková et al. 2020; Rial et al. 2019), and despite many attempts with recent technologies and standards, so far no known SL has been identified from *P. patens* (Yoneyama et al. 2018b). An alternative to evaluate the SL-like activity of PpCCD8-derived molecules is to test the ability of *P. patens* exudates to induce the germination of parasitic seeds. Parasitic plants such as *Phelipanche ramosa* can parasitize various host plants, in response to specific exuded germination stimulants (GS). Different genetic groups of *P. ramosa* seeds have been identified, depending on the crop grown in the field where the seeds were collected (Huet et al. 2020). Seeds from two populations of *P. ramosa* harvested in hemp (*P. ramosa* group 2a) and oilseed rape (*P. ramosa* group 1) fields (Stojanova et al. 2019; Huet et al. 2020) were assayed with WT moss exudates (Figure 1). As a control, both groups of seeds were germinated in the presence of (±)-GR24 (Figure 1A). WT moss exudates induced germination of *P. ramosa* group 2a seeds but not *P. ramosa* group 1 seeds (Figure 1A). In another assay, *P. ramosa* seeds were added to culture plates close to WT or *Ppccd8* plants, with and without (±)-GR24 (Figure 1B-C). *P. ramosa* group 2a but not group1 seeds germinated on WT moss plates, while no germination was observed in the vicinity of *Ppccd8* plants. In all cases (WT and *Ppccd8*), the addition of (±)-GR24 to the medium restored seed germination. Thus, PpCCD8-derived compounds induce the germination of a specific population of *P. ramosa* seeds, responding to not yet identified GS exuded by hemp.

**Figure 1:**
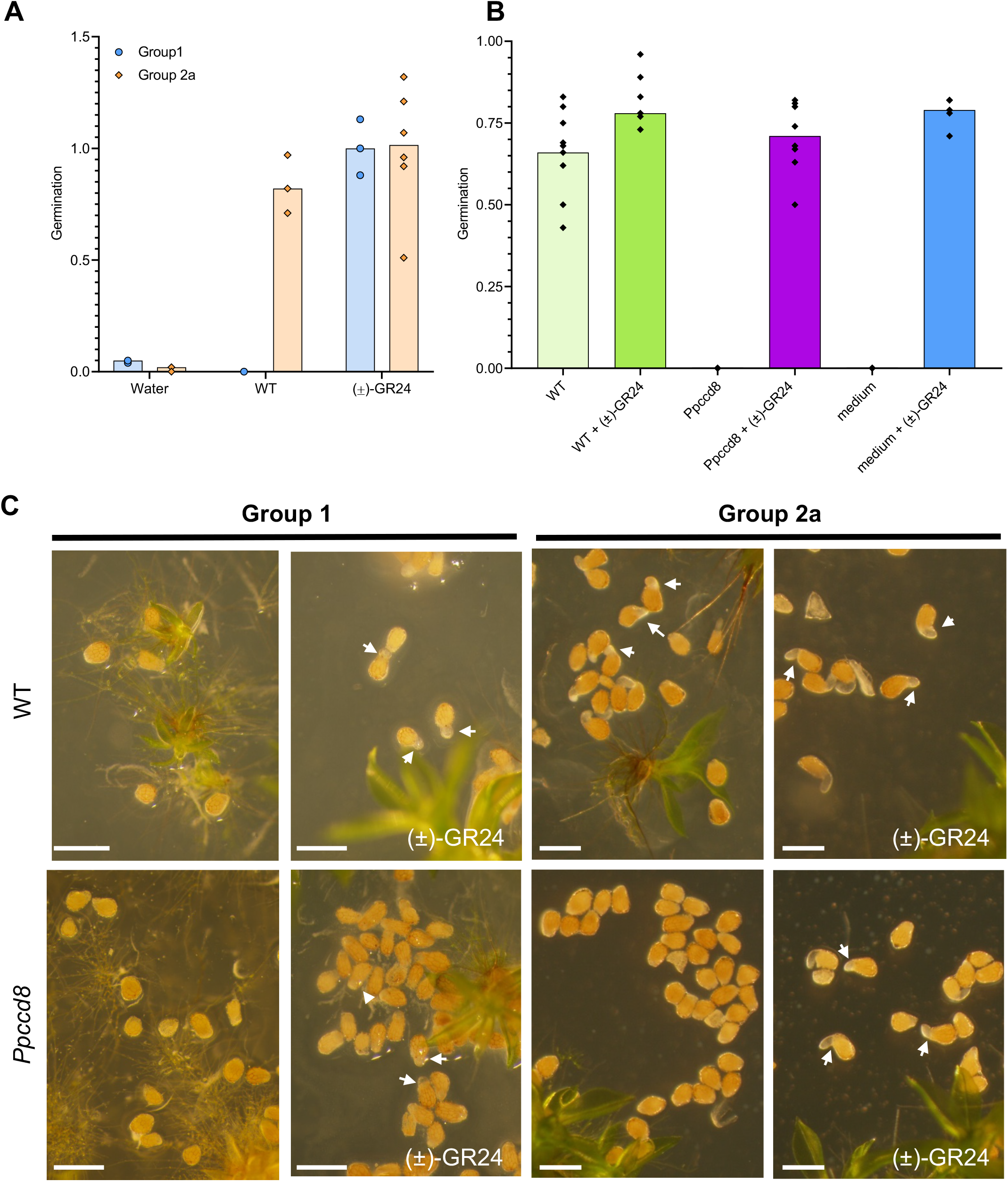
PpCCD8-derived compounds are germination stimulants (GS) of a specific group of *Phelipanche ramosa*. **(A)** GS activities of *P. patens* exudates on *P. ramosa* group 1 and 2a seeds relative to 0.1 μM (±)-GR24. **(B)** Germination rate of seeds of *P. ramosa* group 2a on plates with *P. patens* WT, *Ppccd8* mutant plants, or with culture medium only, with or without 0.1 μM (±)-GR24. **(C)** Seeds from *P. ramosa* group 1 (left) and group 2a (right) on plates with WT (top) or *Ppccd8* mutant plants (bottom), with or without 0.1 μM(±)-GR24. Arrows indicate germinating seeds. Scale bar = 0.5 mm.

### *P. patens* responds strongly to (+)-GR24 and carlactone application, but poorly to (-)-GR24 and KAR2 in the dark

With the aim of determining which molecules could be used to mimic the yet unknown PpCCD8-derived compounds and moss KL, we tested the two most active enantiomers of GR24 ((+)-GR24 and (-)-GR24), the SL precursor carlactone (CL), and the Karrikin KAR2. The length of *caulonema* filaments grown in the dark has been used as a proxy for quantifying the *P. patens* phenotypic response to SL using (±)-GR24 (Hoffmann et al. 2014; Lopez-Obando et al. 2018). Here we relied on a new protocol (Guillory and Bonhomme 2021), and found that the number of caulonemal filaments per plant ended up a more robust proxy of the response to compounds than *caulonema* length (Figure 2 and Supplemental Figure 1). Since the (±)-GR24 can activate non-specific responses (Scaffidi et al. 2014), we tested the (+) and (-)-GR24 enantiomers separately on both WT and the *Ppccd8* mutant. As in previous studies (Hoffmann et al. 2014; Lopez-Obando et al. 2018), we predicted a clearer response in the synthesis mutant than in WT, due to the absence of endogenous PpCCD8-derived compounds potentially mimicked by (+)-GR24. Both the number and the length of *caulonema* filaments significantly decreased following application of (+)-GR24 in WT and the *Ppccd8* mutant, in a dose-response manner (Figure 2 and Supplemental Figure 1). A dose of 0.1 μM was sufficient to see a clear and significant response in terms of number in both genotypes (Figure 2). Application of (-)-GR24 led to a significant decrease in *Ppccd8* filament length, as did (+)-GR24, but there was no effect on WT filament length (Supplemental Figure 1). Strikingly, no significant changes in *caulonema* filament number were observed with (-)-GR24, except in WT for which the 0.1 μM and 10 μM doses led to a significant increase (more pronounced at 0.1 μM, Figure 2A). However, in further assays (see below, Figure 11), a slight but significant decrease of *Ppccd8 caulonema* filament number was observed following (-)-GR24 application. In summary, the phenotypic response of *P. patens* to (+)-GR24 was pronounced and significant with a decrease in both filament number and length in WT and *Ppccd8*. The response to (-)-GR24 was less clear, with sometimes opposite effects between the two genotypes.

**Figure 2:**
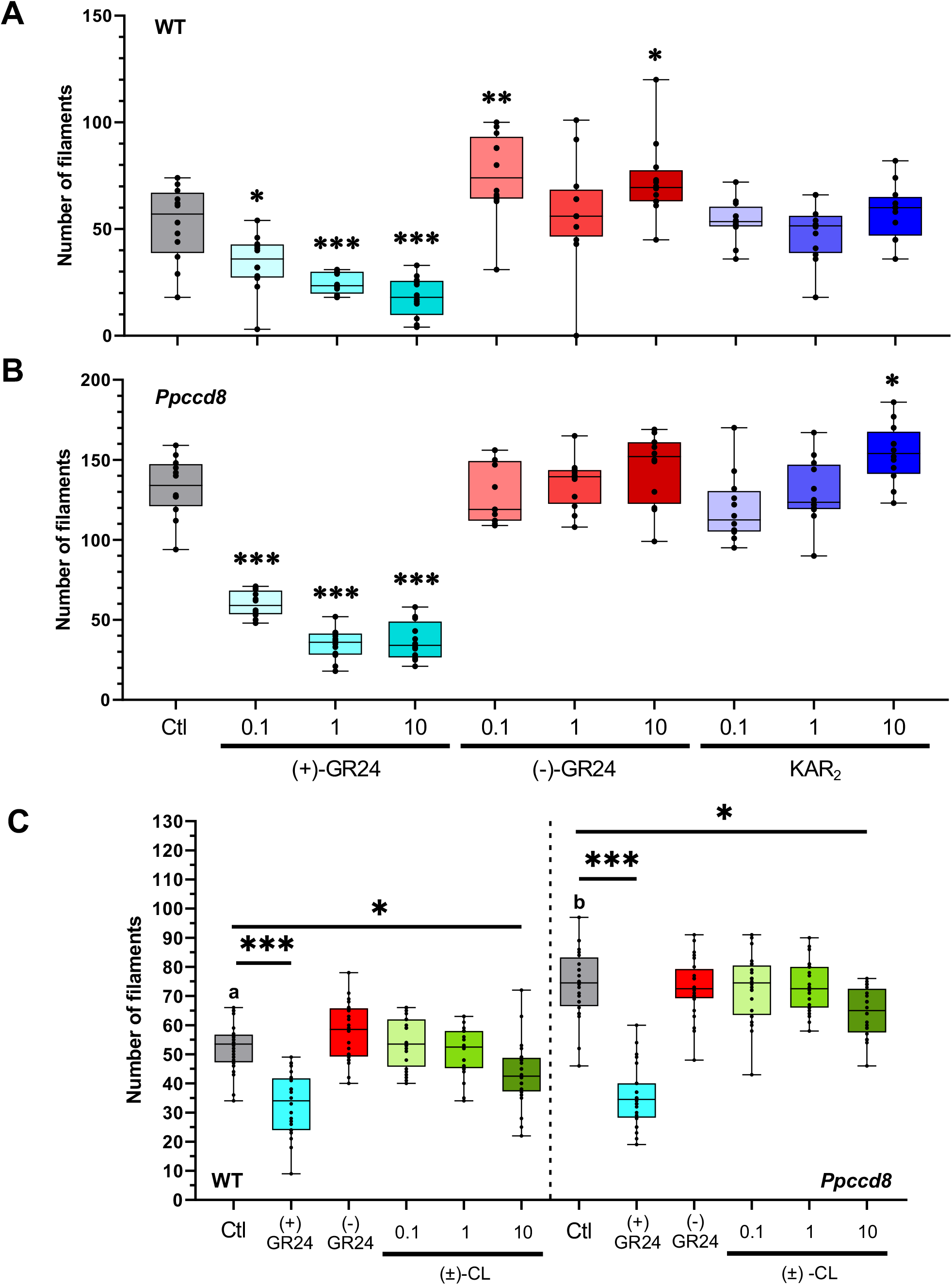
Phenotypic response to (+)-and (-)-GR24 enantiomers and natural compounds: number of *caulonema* filaments. *Caulonema* filaments were counted for WT **(A)** and the *Ppccd8* SL synthesis mutant **(B)** grown for 10 days vertically in the dark, following application of increasing concentrations (0.1, 1 and 10 μM) of (+)-GR24 (blue boxes), (-)-GR24 (red boxes) and KAR_2_, (dark blue boxes). Control: 0.01% DMSO. **(C)** *Caulonema* filament numbers of WT and *Ppccd8* mutant grown for 10 days vertically in the dark, following application of increasing concentrations (0.1, 1 and 10 μM) of (±)-CL (green boxes). Control (Ctl): 0.01% DMSO. (+)-GR24 (blue boxes) and (-)-GR24 (red boxes) were applied at 1 μM. Significant differences between control and treated plants within a genotype based on a Kruskal-Wallis test, followed by a Dunn *post-hoc* test for multiple comparisons: ***, P<0.001; **, P<0.01; *, P<0.5; For each genotype and treatment, n = 24 plants grown in three different 24 well-plates.

Recent biochemistry experiments (Bürger et al. 2019) showed that some PpKAI2L proteins could bind KAR_1_ (PpKAI2L-H, K and L), while a previous study concluded that *P. patens* was insensitive to KAR_1_ (Hoffmann et al. 2014). Indeed, no transcriptional *(PpCCD7* expression) or phenotypic response (*caulonema* length in the dark) to KAR_1_ was observed in either WT or the *Ppccd8* mutant (Hoffmann et al. 2014). In the present work, we tested the KAR2 molecule, described as being more active than KAR_1_ in Arabidopsis (Waters et al. 2015; Yao et al. 2021) (Figure 2A-B). KAR2 has an unmethylated butenolide group, unlike KAR_1_. In WT, no significant effect on *caulonema* number was observed following application of increasing doses of KAR2 (Figure 2A). In *Ppccd8*, we observed an increase in filament number, as with (-)-GR24 in WT, and this increase was significant at 10 μM (Figure 2B). Surprisingly, the length of WT filaments decreased significantly with the 1 μM dose only, while higher KAR2 concentrations (10 μM) had no significant effect (Supplemental Figure 1). KAR2 application also reduced *caulonema* length in the *Ppccd8* mutant but at low concentrations (0.1 μM and 1 μM, Supplemental Figure 1). To conclude, KAR2 phenotypic effects on *P. patens* were slight and not clearly dose responsive, albeit in a different fashion than those of (-)-GR24.

We also tested racemic carlactone (CL), described as the natural product of PpCCD8 in *P. patens* (Decker et al. 2017). As previously reported (Decker et al. 2017), CL application had a negative effect on *caulonema* filament number for both WT and the *Ppccd8* mutant, however in our assays the effect was only significant at 10 μM (Figure 2C).

Overall, findings from these phenotypic assays suggest that GR24 enantiomers have distinct effects in *P. patens,* as observed in Arabidopsis (Scaffidi et al. 2014). Indeed, the (+)-GR24 analog mimics CL effects, although it is far more potent, and can thus be used to mimic the effects of PpCCD8-derived compounds. On the other hand, the (-)-GR24 analog and KAR2 have slight phenotypic effects which tend to be opposite to those of PpCCD8-derived compounds.

### All *PpKAI2L* genes are expressed at relatively low levels and putatively encode proteins with a conserved catalytic triad

As for D14 and KAI2 receptors, a catalytic triad (Ser, His, Asp) is encoded by all 13 *PpKAI2L* genes (Figure 3 and Supplemental Figure 2), including *PpKAI2L-A* and -*M* for which recent sequencing data (P. patens v3.3, (Lang et al. 2008)) contradicts previous pseudogene predictions. In the following, the four subclades previously described in (Lopez-Obando et al. 2016a), will be renamed for convenience: clade (A-E) for clade (i) including PpKAI2L-A, -B, -C, -D, and -E, clade (FK) for clade (ii) including PpKAI2L-F and -K, clade (HIL) for clade (i.i-i.ii) including PpKAI2L-H, -I, -L, and clade (JGM) for clade (iii) including PpKAI2L-J, - G, and -M.

**Figure 3:**
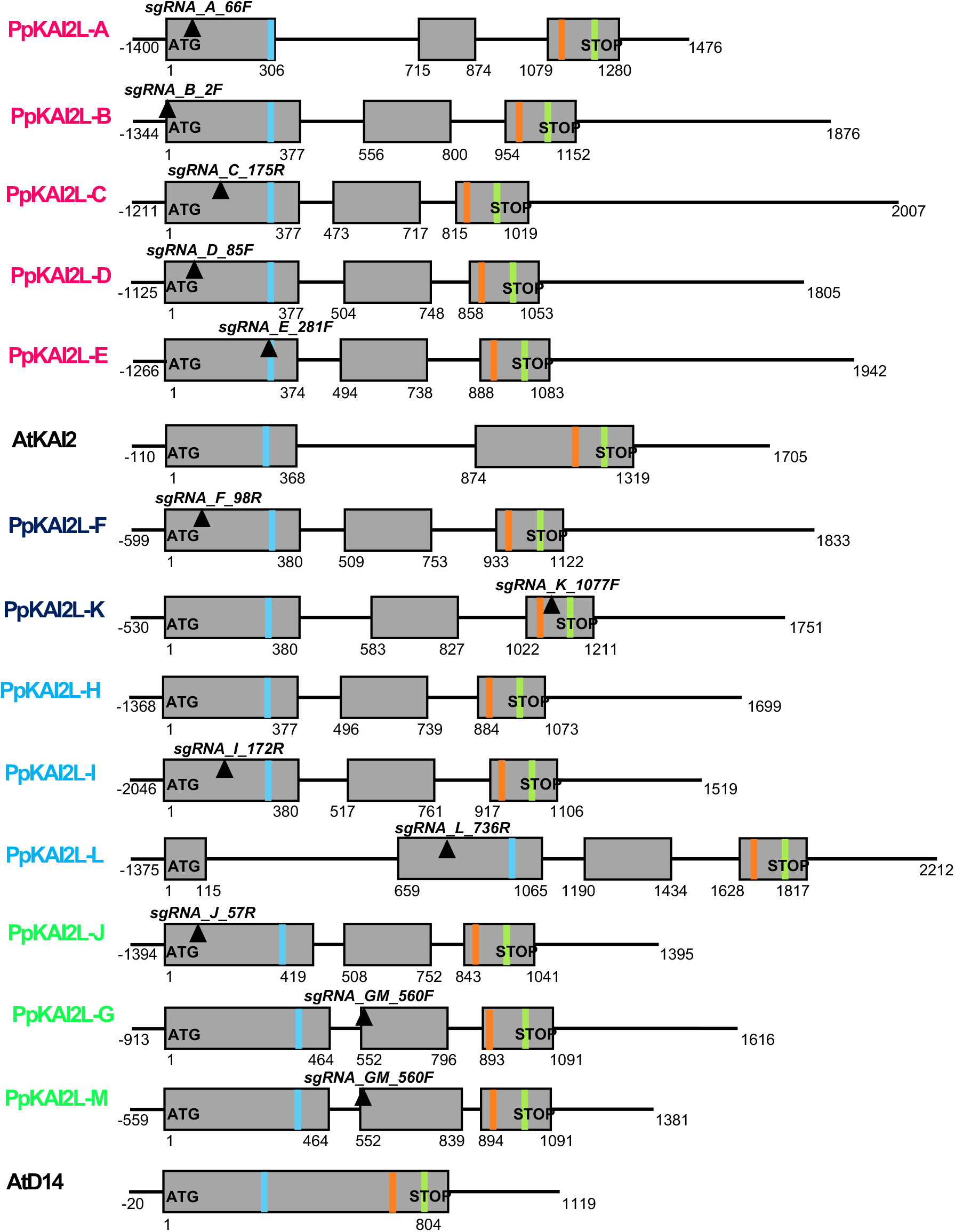
Gene models of the *PpKAI2L* gene family showing the catalytic triad and position of crRNAs. Genes are presented as organized in subclades. Exons are displayed as grey boxes, introns and UTRs are depicted as thin black lines. Start and Stop codons are written in bold, while plain text indicates the start/end position for each feature, relative to the start codon. Only 5’-UTR regions are not represented true to scale. Transcript versions that were used are V3.1 (downloaded from the Phytozome website in September 2019) for all *PpKAI2L* genes except for *PpKAI2L-B, PpKAI2L-H* and *PpKAI2L-M* (V3.2). Regions targeted by crRNAs are indicated as black inverted triangles, with their names written in bold italic. Light blue, orange and light green bands represent respectively the codons for the S, D and H residues of the catalytic triad (see Supplemental Table 3 for reference sequences).

To test if the high number of PpKAI2L genes hint at different spatial expression profiles, we obtained the expression pattern of all *PpKAI2L* genes in *P. patens,* using a cDNA library from various organs/tissues (Supplemental Figure 3 A), including spores, *protonema* of increasing age and different tissue composition (6-day-old: primarily *chloronema*; 11-day-old: mix of *chloronema* and *caulonema;* 15-day-old: mix of *chloronema, caulonema* and gametophores buds), and gametophores from 5-week-old plants. *PpKAI2L* genes transcripts were detected in all tested tissues, at relatively low levels compared to the control genes. Notably, we also found that *PpKAI2L-A* is expressed, thus confirming it is not a pseudogene. Quantitative -PCR could not be used to assess expression of *PpKAI2L-M*, as its predicted transcript is almost identical to that of *PpKAI2L-G,* and the observed transcript levels are attributable to both *PpKAI2L-G* and *-M*. In spores, *PpKAI2L-F* and *-J* had higher transcript levels than other *PpKAI2L* genes. In *protonema* and gametophores however, *PpKAI2L-D* from clade (A-E) showed the highest transcript levels compared to any other *PpKAI2L* genes. *PpKAI2L-I* transcript levels are the lowest, in all tested tissues (Supplemental Figure 3A). When considering the clades separately (Supplemental Figure 3B-E), *PpKAI2L-D* had the highest transcript levels among clade (A-E) genes, while *PpKAI2L-H* transcript levels were higher compared to that of *PpKAI2L-I* and -*L* (clade (HIL)). *PpKAI2L-F* and -*K* (clade (FK)) showed comparable transcript levels, and *PpKAI2L-J* had slightly higher transcript levels than *PpKAI2L-G/M* (clade (JGM)). The data are consistent with those previously reported (Ortiz-Ramirez et al. 2016), shown in Supplemental Figure 4.

### The PpKAI2L-C, -D, -E proteins are destabilized by (-)-GR24 as observed for AtKAI2; PpKAI2L-F, -K, -L and -H interact weakly with GR24 enantiomers

To investigate whether the PpKAI2L proteins behave similarly to AtD14 or AtKAI2 *in vitro*, all *PpKAI2L* CDS were cloned and over-expressed in *E. coli*. After successful purification and solubilization of seven PpKAI2L proteins (-C,-D,-E,-F,-H,-K and -L), their interaction with SL analogs and their potential enzymatic activities were investigated. Unfortunately, because of low solubility, the six other PpKAI2L proteins could not be purified in sufficient amounts to ensure their quality. Notably, none from the (JGM) clade could be characterized biochemically.

Interactions of purified PpKAI2L proteins were tested with SL analogs, in both nano Differential Scanning Fluorimetry (nanoDSF) and classical DSF (Figure 4 and Supplemental Figure 5). All four GR24 isomers (Figure 4A and Supplemental Figure 5H) destabilized AtD14 (Figure 4B and Supplemental Figure 5A and 5I), while AtKAI2 was destabilized by (-)-GR24 addition only (Figure 4C and Supplemental Figure 5B and 5J), as previously reported (Waters et al. 2015). All tested clade (A-E) proteins (PpKAI2L-C -D and -E) were destabilized when (-)-GR24 was added, as was AtKAI2 (Figure 4D to 4F and Supplemental Figure 5C to 5E and 5K to 5M), in accordance with the reported stereoselectivity for un-natural (-)-5DS (Bürger et al. 2019). Puzzingly, the PpKAI2L-C, -D and -E proteins showed a tendency to be stabilized by (+)-GR24 at high concentrations (Supplemental Figure 5C to 5E), not reported when using (+)-5DS. None of the GR24 isomers affected the stability of the PpKAI2L-F (clade (FK)) and PpKAI2L-H (clade (HIL)) proteins (Figure 4G and 4I and Supplemental Figure 5F, 5G, 5N and 5P). PpKAI2L-K (clade (FK)) was destabilized by both (-)-GR24 and (+)-GR24 (Figure 4H), and stabilized by (+)-2’-*epi*-GR24 (T_m_ + 1.5 °C) (Supplemental Figure 5O). PpKAI2L-L (clade (HIL)) showed a slight increase in Tm following addition of all isomers (≤ 1°C), suggesting a slight stabilization (Figure 4J and Supplemental Figure 5Q).

**Figure 4:**
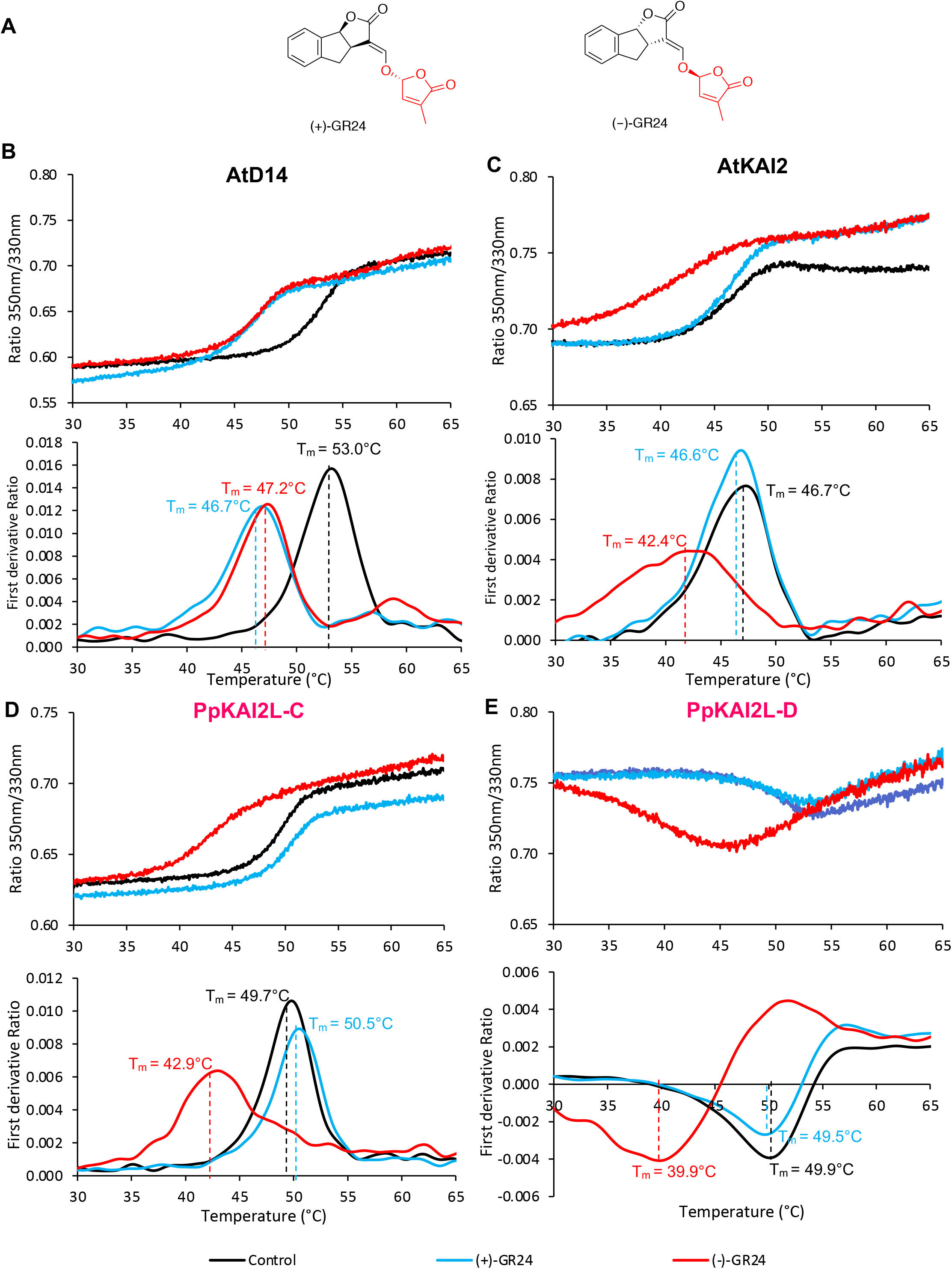
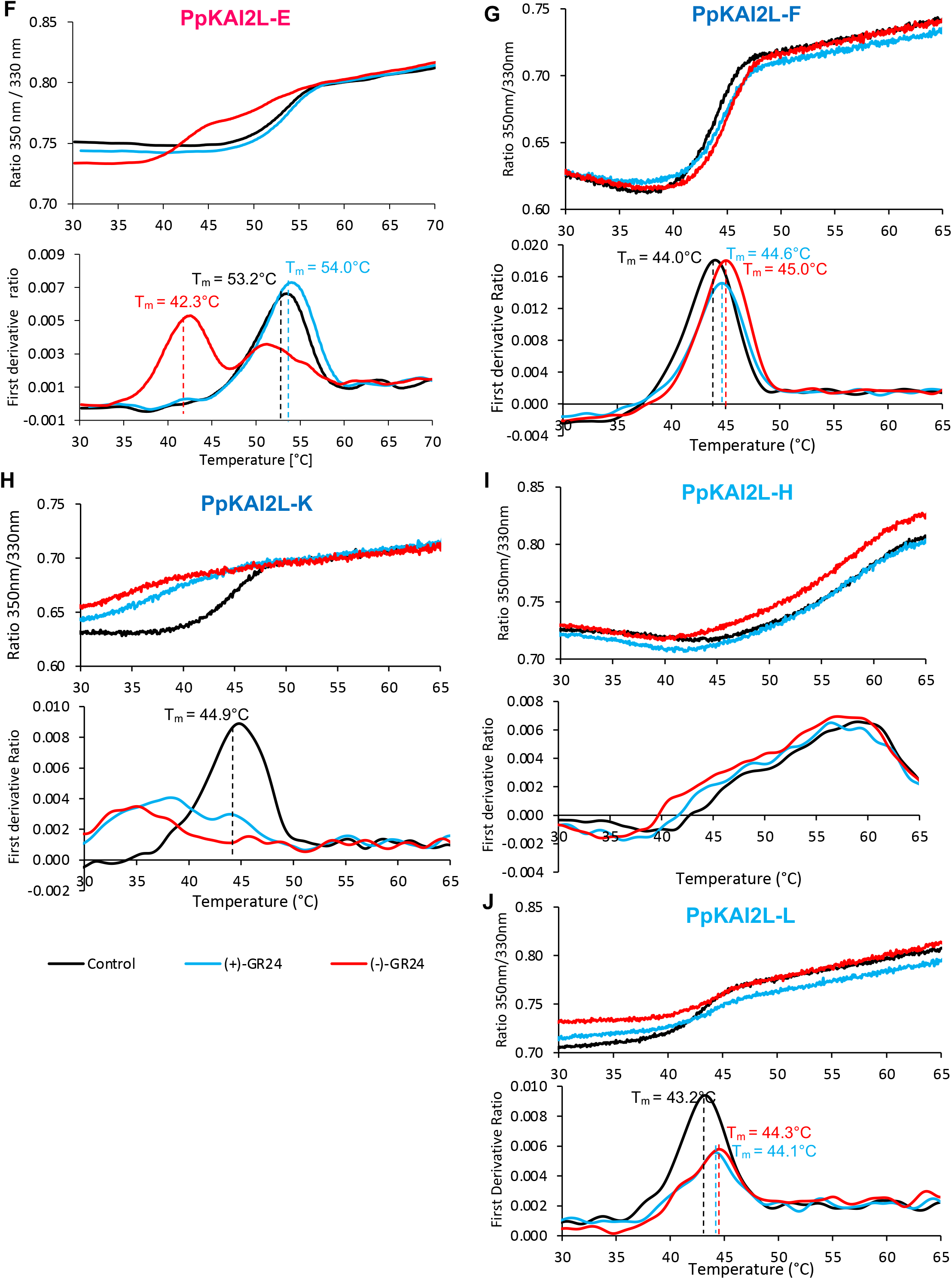
Nano Differential Scanning Fluorimetry (nanoDSF) analysis of the PpKAI2L protein response to GR24 isomers. **(A)** Chemical structures of the (+)-GR24 and (-)-GR24 enantiomers. **(B-J)** Thermostability of AtD14, AtKAI2 and PpKAI2L proteins at 10 μM in the absence of a ligand (black line) or in the presence of (+)-GR24 (blue line) or (-)-GR24 (red line) at 100 μM, analyzed by nanoDSF. For each protein, the top panel shows changes in fluorescence (ratio F_350nm_/F_330nm)_ with temperature. The bottom panel shows the first derivative for the F_350nm_/F_330nm_ curve, against the temperature gradient from which the apparent melting temperatures (T_m_) for each sample was determined. The experiment was carried out twice.

Since we observed sometimes opposite effects on the PpKAI2L proteins stability with the different isomers, we reasoned that their binding affinity had to be assessed for the GR24 isomers, rather than for (±)-GR24 as previously reported (Bürger et al. 2019). Binding affinities were quantified with *K_d_* affinity calculations following intrinsic tryptophan fluorescence measurements (Figure 5 and Supplemental Figure 6). Both PpKAI2L-D and PpKAI2L-E showed a comparable affinity for (-)-GR24, (respectively 92 μM and 39 μM), similar to that recorded for AtKAI2 (45 μM) and for AtD14 (94 μM) (Figure 5A to 5D). In contrast to the AtD14 control (Figure 5A), but as for AtKAI2, the *K_d_* value for (+)-GR24 could not be determined for clade (A-E) proteins, indicating low affinity. While no change in PpKAI2L-H protein stability was observed in nanoDSF, interactions between this protein and three GR24 isomers ((-)-GR24, (+)-GR24 and (-)-2’-*epi*-GR24) were detected (Figure 5G and supplemental Figure 6G). *K_d_* values between 100 μM and 200 μM were estimated, indicating a weak affinity for these compounds. Finally, intrinsic flurorescence assays with PpKAI2L-F and PpKAI2L-K confirmed that these two proteins behave differently: PpKAI2L-F appeared not to interact with any GR24 isomers, whereas PpKAI2L-K showed an affinity of 41 μM for (-)-GR24 and 107 μM for (+)-GR24 (Figure 5E-F).

**Figure 5:**
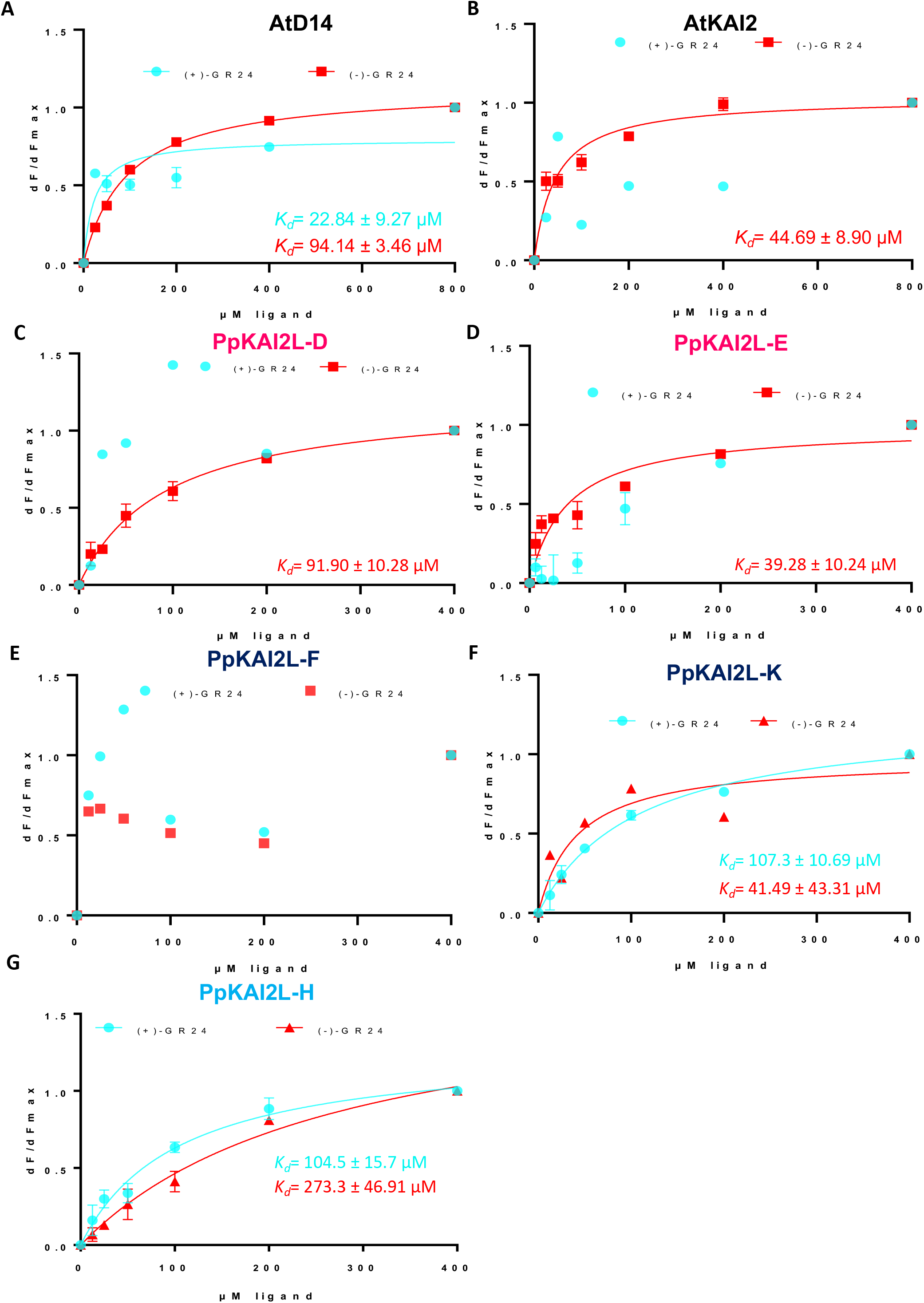
Intrinsic tryptophan fluorescence shows that GR24 isomers bind to PpKAI2L proteins with different affinities. Plots of fluorescence intensity *versus* concentration of (+)-GR24 (turquoise) or (-)-GR24 (red). The change in intrinsic fluorescence of AtD14 **(A)**; AtKAI2 **(B)**; PpKAI2L-D **(C)**; PpKAI2L-E; **(D)** PpKAI2L-F **(E)**; PpKAI2L-K **(F)**; and PpKAI2L-H **(G)** was monitored (see Supplementary Figure 4) and used to determine the apparent *K_d_* values. The plots represent the mean of two replicates and the experiments were repeated at least three times. Graph generation and analysis were carried out using the GraphPad Prism 7.0 Software.

### PpKAI2L-C, -D, -E, and -K preferentially cleave (-)-GR24, while PpKAI2L-H and -L cleave both (-)-GR24 and (+)-GR24

As all PpKAI2L proteins contain the conserved catalytic triad and since we found that most of them were able to bind at least one of the GR24 isomers, we tested their enzyme activity against SL analogs. First, when incubated with the generic substrate for esterases, 4-nitrophenyl acetate (*p*-NPA), AtKAI2 and all tested PpKAI2L proteins showed enzyme activity (Supplemental Figure 7A-B), consistent with previous report (Bürger et al. 2019). Kinetic constants were in the same range for all proteins but one, and similar to that of AtKAI2. The exception was PpKAI2L-H, which had a higher *V_max_* and *K_M_*, highlighting faster catalysis and a better affinity for *p*-NPA than all the others (Supplemental Figure 7C). Moreover, since different binding affinities were observed (see above), we further characterized the enzyme activity of PpKAI2L proteins by testing their hydrolysis activity on the GR24 isomers (Figure 6). We compared their substrate bias to that of the pea SL receptor, RMS3/PsD14, and to that of AtKAI2 (Figure 6A). All three purified clade (A-E) proteins (PpKAI2L-C -D and -E) showed comparable enzymatic stereoselectivity towards the (-)-GR24 isomer (between 15 and 20%), close, though significantly lower, to that of AtKAI2 that reached 30%. In contrast to AtKAI2, PpKAI2L-C could also cleave the (+)-GR24 isomer, nevertheless to a lesser extent (5-10%) suggesting a less stringent selectivity. As for PpKAI2L proteins from clade (FK), PpKAI2L-F showed very low activity towards both isomers (less than 5%), while PpKAI2L-K activity was comparable to that of clade (A-E) and to that of PpKAI2L-L (clade (HIL)). Finally, although none of the PpKAI2L proteins showed as high enzyme activity as RMS3 (100% cleavage of (+)-GR24 and (-)-GR24)), the PpKAI2L-H protein showed a relatively high catalytic activity towards the GR24 isomers, especially towards (-)-GR24 (almost 70%, Figure 6A).

**Figure 6:**
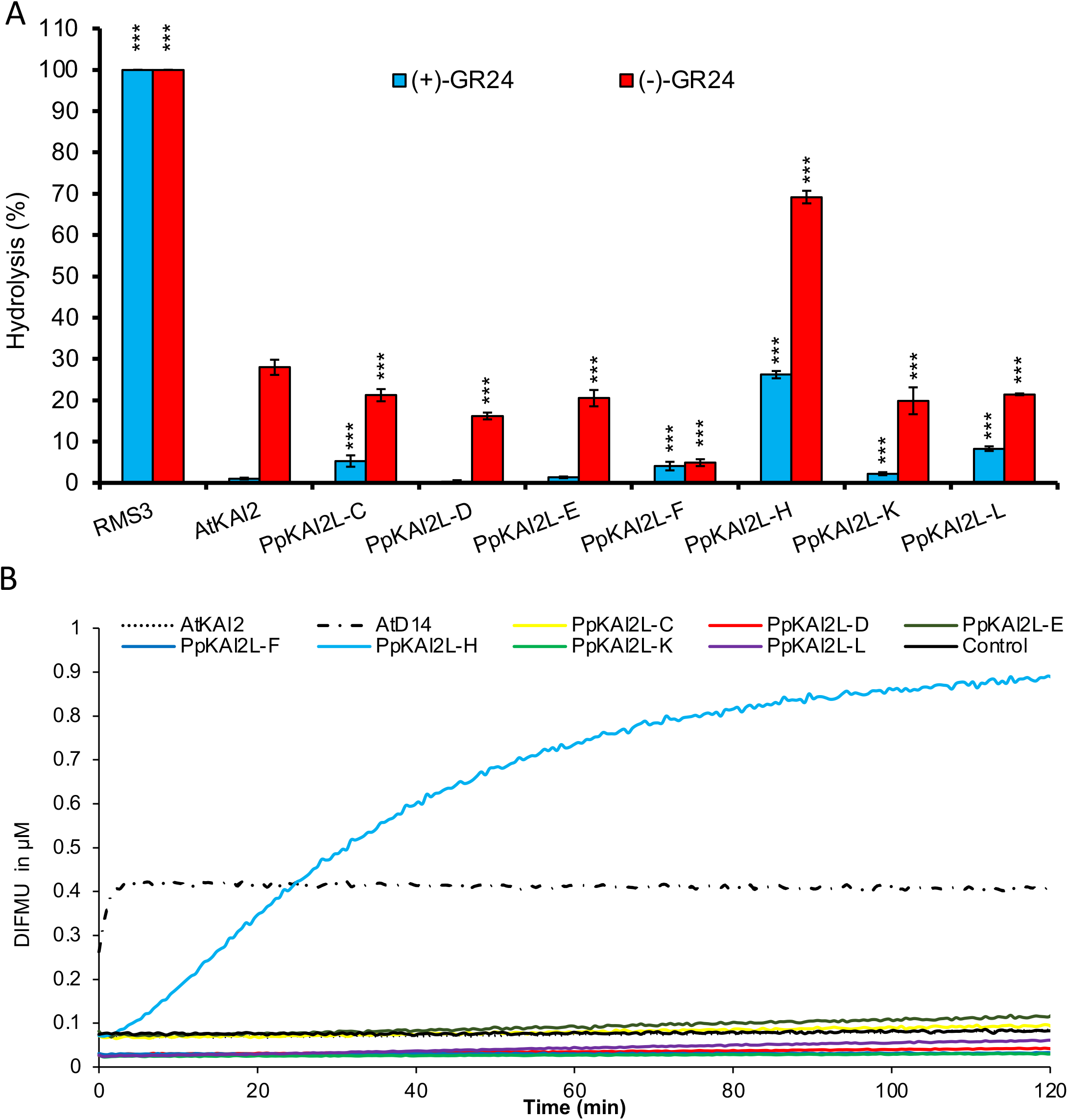
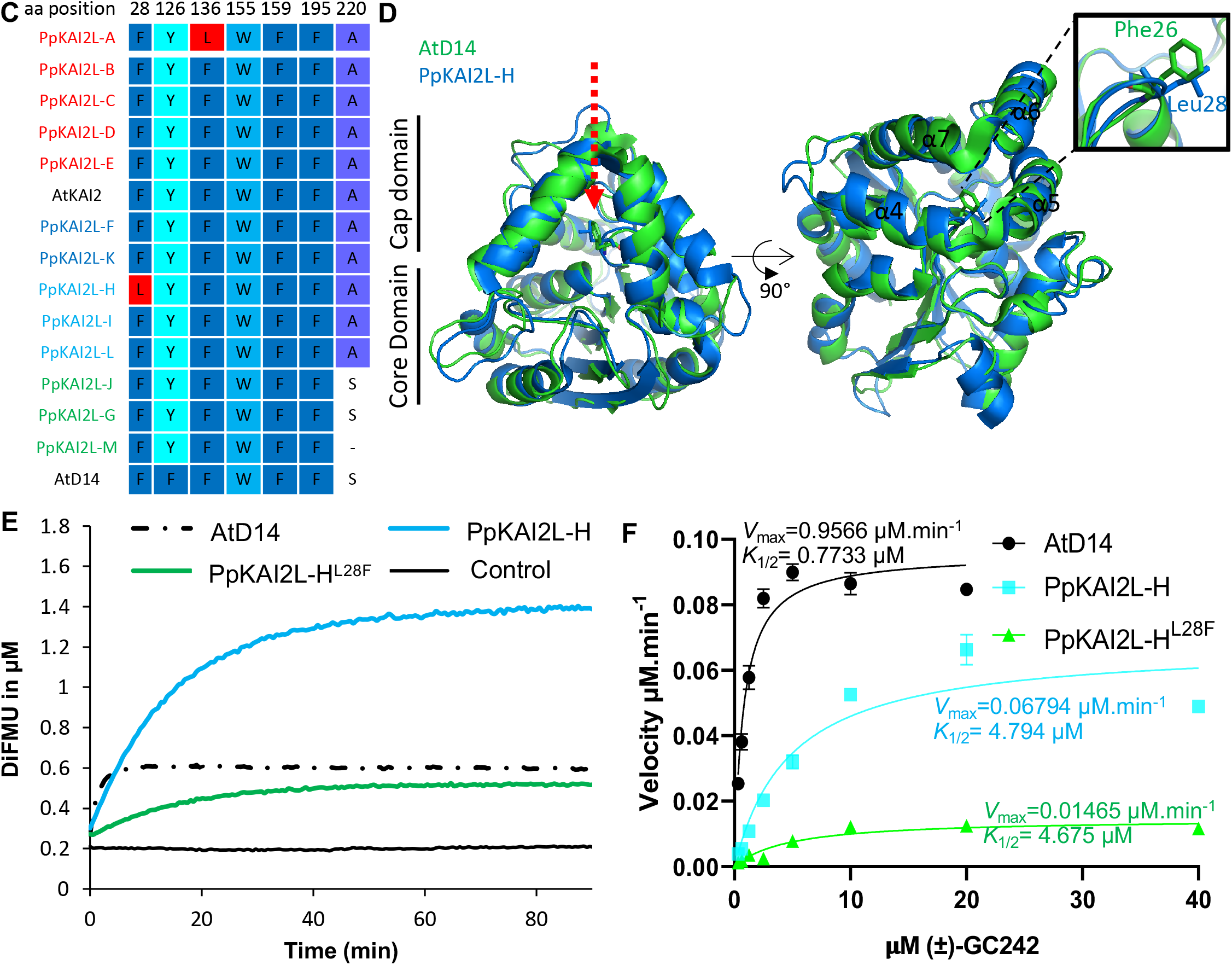
PpKAI2L enzymatic activities confirm stereoselectivity and reveal PpKAI2L-H special feature. **(A)** PpKAI2L enzymatic activity towards GR24 isomers; (+)-GR24 and (-)-GR24 at 10 μM were incubated with 5 μM RMS3, AtKAI2 or the seven PpKAI2L proteins for 150 min at 25 °C. UPLC-UV (260 nm) analysis was used to detect the remaining amount of GR24 isomers. Columns represent the mean value of the hydrolysis rate calculated from the remaining GR24 isomers, quantified in comparison with (±)-1-indanol as internal standard. Error bars represent the SD of three replicates (means ± SD, n = 3). Asterisks indicate statistical significance from the AtKAI2 values, for each isomer, as assessed for each compound by a Dunnett nonparametric relative contrast effects test, with AtKAI2 considered as the control group (*** p ≤ 0.001). **(B)** Enzyme kinetics for PpKAI2L, AtD14 and AtKAI2 incubated with (±)-GC242 (structure shown in Supplemental Figure 8A). Progress curves during probe hydrolysis, monitored at 25 °C (λ_em_ 460 nm). Protein catalyzed hydrolysis with 330 nM of protein and 4 μM of probe. These traces represent one of the three replicates and the experiments were repeated twice. **(C)** Sequence alignment of active site amino acid residues in PpKAI2L proteins. Amino acids that differ from AtKAI2 are colored in red. A fully expanded alignment can be found in Supplemental Figure 2. **(D)** Superimposition of the AtD14 and PpKAI2L-H structure showing the position of F^28^ and L^28^ residues. Zoom onto helices α4 and α5. **(E)** Enzyme kinetics for PpKAI2L-H, PpKAI2L-H^L28F^ and AtD14 incubated with (±)-GC242. Progress curves during probe hydrolysis, monitored at 25 °C (λ_em_ 460 nm). Protein catalyzed hydrolysis with 330 nM of protein and 20 μM of probe. These traces represent one of the three replicates and the experiments were repeated twice. **(F)** Hyperbolic plot of pre-steady state kinetics reaction velocity with (±)-GC242. Initial velocity was determined with pro-fluorescent probe concentrations from 0.310 μM to 40 μM and protein at 400 nM. Points are the mean of three replicates and error bars represent SE. Experiments were repeated at least three times.

### PpKAI2L-H high cleavage activity on synthetic SL analogs is explained by a specific leucine residue

The high enzyme activity of PpKAI2L-H (Figure 6A) and its lack of thermal shift when incubated with GR24 isomers (Figure 4I and Supplemental Figure 5N and 5P) set this protein apart from other PpKAI2L. To better characterize PpKAI2L-H enzyme activity, we used a pro-fluorescent probe as substrate ((±)-GC242), in which the ABC rings of GR24 are replaced by a coumarine-derived moiety (DiFMU) (de Saint Germain et al. 2016). (±)-GC242 is bioactive on moss as it reduced the number of *caulonema* filaments in the dark in a dose responsive manner (evaluated on the *Ppccd8* mutant, Supplemental Figure 8A). The use of (±)-GC242 as substrate confirmed PpKAI2L-H relatively high enzyme activity, compared to all the other PpKAI2L proteins (Figure 6B). Indeed, after 2 hours, PpKAI2L-H catalyzed the formation of 1μM DiFMU, while other PpKAI2L activities were indistinguishable from background noise. However, the PpKAI2L-H enzymatic profile did not show the biphasic curve (a short burst phase, quickly followed by a plateau phase), which characterizes the AtD14 single turnover activity (Figure 6B, (de Saint Germain et al. 2016)). The lack of a plateau for PpKAI2L-H suggested that this protein acted as a Michaelian enzyme on the SL analog. To try to understand this singularity, we compared the solvent exposed residues in the binding pocket of the PpKAI2L proteins and noticed that PpKAI2L-H harbors a leucine^28^ (L^28^) residue instead of the phenylalanine found in AtD14 (F^26^), AtKAI2 and all other PpKAI2L proteins (Figure 6C). The F residue is located at the junction between helix α4 and α5, near the catalytic site, and can precisely interact with the D-ring of the SL (Figure 6D). Furthermore, a mutant PpKAI2L-H protein where L^28^ is changed to F showed a biphasic cleavage profile similar to AtD14, both reaching a plateau at 0.4μM DiFMU, correlating with the protein concentration (Figure 6E and 6F). PpKAI2L-H and PpKAI2L-H^L28F^ proteins had comparable affinity towards (±)-GC242 *(K_1/2_*= 4,794 μM *vs* 4,675 μM) but showed different *V_max_* values (*V_max_*=0,06794 μM.min^-1^ *vs* 0,01465 μM.min^-1^), suggesting that the L^28^ residue affects the velocity of catalytic activity (Figure 6E and 6F).

### Moss PpKAI2L proteins, like vascular plant receptors, covalently link GR24 enantiomers

To further investigate whether PpKAI2L proteins play a role as receptors, we examined covalent attachment of the GR24 isomers to the PpKAI2L proteins (Supplemental Figure 9). Mass spectrometry analyses highlighted 96 Da increments (corresponding to the D ring mass), when AtKAI2 was incubated with (-)-GR24, and all PpKAI2L-C, -D, -E, -F or -L were incubated with (-)-GR24. Strikingly, 96 Da increments were also observed when PpKAI2L-E -F -L and -K were incubated with the other isomer (+)-GR24, in contrast to other reports (Bürger et al. 2019). However, for PpKAI2L-E, the intensity peak was much lower with (+)- GR24 than with (-)-GR24, confirming the better affinity for the latter (Figure 5). PpKAI2L-H did not covalently bind the D ring, following incubation with either of both enantiomers, further suggesting that it displays Michaelian enzyme activity.

Thus, poor interactions were observed with (+)-GR24, which was reported to mimick SLs, and had the most potent effect in our phenotypic assays on *P. patens* (Figure 2 and Supplemental Figure 1). Strikingly, all clade (A-E) PpKAI2L proteins showed the strongest affinity for the (-)-GR24 enantiomer, reported in vascular plants as a good mimic for the putative KL (Scaffidi et al. 2014; Zheng et al. 2020). *In planta* studies were then carried out to investigate if (and which of) these PpKAI2L homologs are necessary for SL or KL perception.

### None of the *PpKAI2L* genes complement the Arabidopsis *d14-1 kai2-2* double mutant

We used cross species complementation assays to test whether any of the PpKAI2L proteins could carry out similar functions to that of AtD14 and/or AtKAI2 in Arabidopsis SL and/or KL signaling (Figure 7). CDS of the *PpKAI2L-C, -D, -F, -G,* and-*J* genes were cloned downstream of the AtD14 or AtKAI2 promoter, and resulting constructs were expressed in the Arabidopsis double mutant At*d14-1 kai2-2*, which shows both a hyperbranched phenotype and elongated hypocotyls (Supplemental Figure 10A to 10C). As controls, the double mutant was transformed with the AtKAI2 or AtD14 CDS under the control of endogenous promoters. Only lines expressing AtD14 under the control of the AtD14 promoter fully restored rosette branching to WT (Ler) values (Figure 7A). Under the control of the AtD14 promoter, neither AtKAI2 or any of the PpKAI2L could significantly restore the branching phenotype of the At*d14-1 kai2-2* mutant, with the exception of one line expressing PpKAI2L-J (#4.6), which showed a significantly lower number of rosette branches than the mutant (still higher than WT or the AtD14 expressing line, Figure 7A). We conclude that none of the tested *PpKAI2L* genes can fully complement the function of AtD14 in shoot branching.

**Figure 7:**
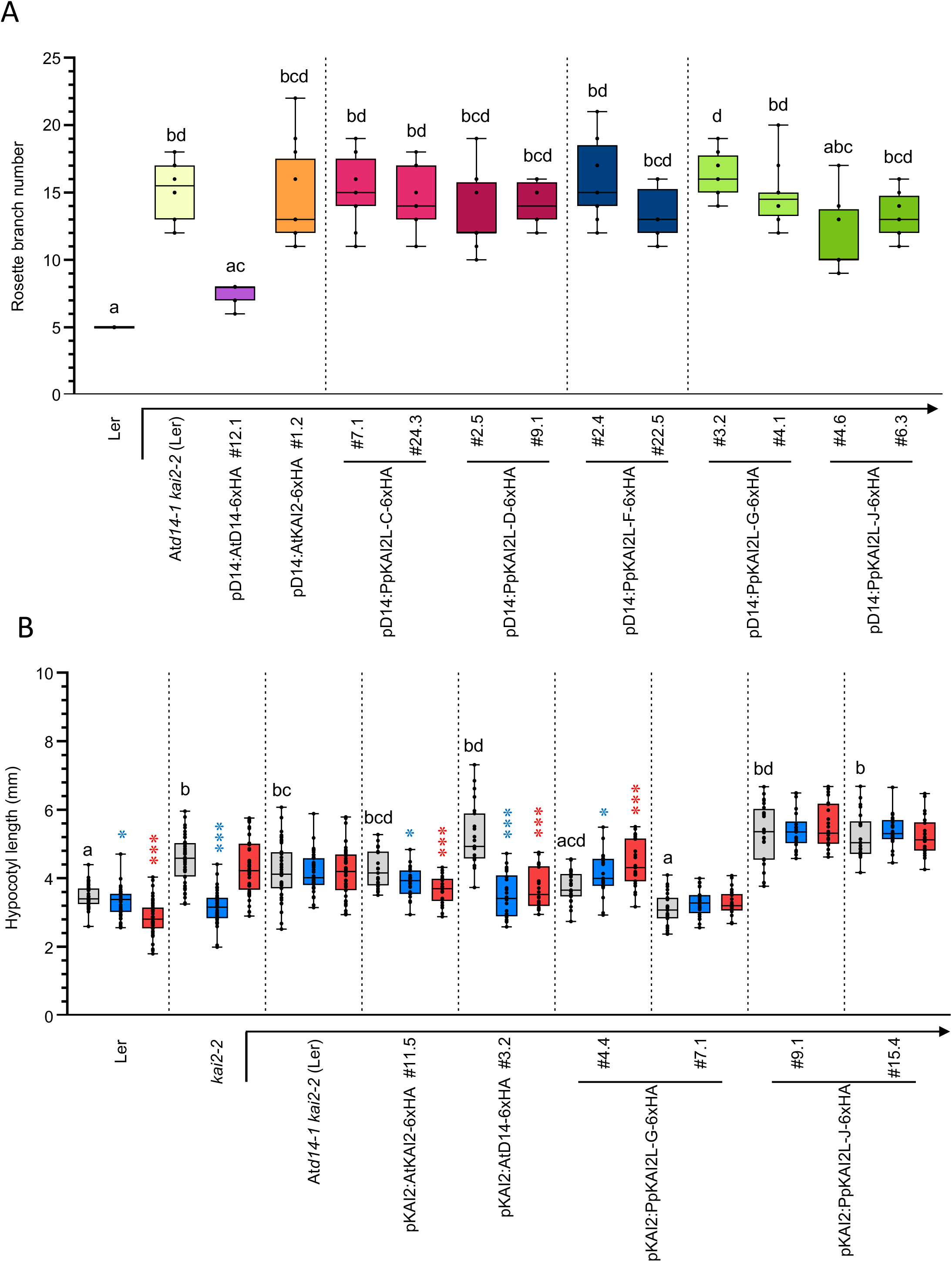
Complementation assays of the Arabidopsis *Atd14-1 kai2-2* double mutant with *PpKAI2L* genes. Complementation assays of the *Atd14-1 kai2-2* mutant (in Ler background), transformed using the *AtD14* promoter **(A)** or *AtKAI2* promoter **(B)** controlling the expression of *AtD14, AtKAI2* (controls) or *PpKAI2L* CDS (as noted below the graph). Ler (WT), *kai2-2* and At*d14-1 kai2-2* mutants are shown as controls. **(A)** Number of rosette axillary branches per plant. Results are mean of n =12 plants per genotype, except for Ler and lines pD14:AtD14 #12.1 and pD14:PpKAI2L-C#24.3 (n = 11). Different letters indicate significantly different results between genotypes based on a Kruskal-Wallis test (P < 0.05, Dunn *post hoc* test with P values corrected following the Benjamini-Hochberg method). **(B)** Hypocotyl length under low light, on ½ MS medium with DMSO (control, grey bars) 1 μM (+)-GR24 (blue bars) or 1 μM (-)-GR24 (red bars). Different letters indicate significantly different results between genotypes in control conditions based on a Kruskal-Wallis test (P < 0.05, Dunn *post hoc* test with P values corrected following the Benjamini-Hochberg method). Symbols in blue and red give the statistical significance of response to (+)-GR24 and (-)-GR24 respectively (Mann-Whitney tests, * 0.01 ≤ p < 0.05, *** p ≤ 0.001).

We also examined possible complementation of the AtKAI2 function in the At*d14-1 kai2-2* mutant, by monitoring hypocotyl length under low light conditions, with or without 1 μM (+)-GR24 or (-)-GR24 in the culture medium (Figure 7 B). Compared to WT, the double mutant showed longer hypocotyls under control conditions, as did the single *kai2-2* mutant. Moreover, neither addition of (+)-GR24 nor (-)-GR24 had any effect on this phenotype in the double mutant. In contrast, (+)-GR24 treatment led to shorter *kai2-2* hypocotyls, likely due to perception and transduction by the AtD14 protein which is still active in the single mutant. As expected from the loss of AtKAI2 function, (-)-GR24 addition had no effect on *kai2-2* hypocotyls. Expressing AtKAI2 under the control of the AtKAI2 promoter surprisingly did not fully restore the hypocotyl length of the double mutant under our control conditions, but it did restore the response to (-)-GR24 as anticipated. Even more surprisingly, the pKAI2:AtD14 expressing line showed longer hypocotyls than the double mutant under control conditions (DMSO). In addition, similar longer hypocotyl phenotypes were found under control conditions for lines expressing PpKAI2L-C (Supplemental Figure 10) and PpKAI2L-J (Figure 7B). This unexpected effect of the introduced α/β hydrolases will be discussed below. In contrast, only lines expressing PpKAI2L-G had short hypocotyls under control conditions, suggesting that the expressed protein had indeed complemented the AtKAI2 function. When either (+)-GR24 or (-)-GR24 was added, short hypocotyls (similar to WT) were observed in the pKAI2:AtD14 expressing line, indicating AtD14-mediated signal transduction of both enantiomers when AtD14 is present in tissues where AtKAI2 is normally active. However, adding GR24 enantiomers in the medium had no such effect on lines expressing PpKAI2L proteins (Figure 7B and Supplemental Figure 10). Overall these assays showed that PpKAI2L-G is able to mediate Arabidopsis KL signaling in hypocotyls, even though it could not fully ensure the AtKAI2 response function.

### Multiplex editing of all *PpKAI2L* genes

Multiplex Gene Editing using CRISPR-Cas9 allowed us to isolate *P. patens* mutants affected in one or several *PpKAI2L* genes (Lopez-Obando et al. 2016b) (Figure 3, Figure 8 and Supplemental Figure 11). For clade (A-E), two triple (*Ppkai2L-a-b-c*, *Ppkai2L-c-d-e)* and two quintuple mutants *(Ppkai2L-a-b-c-d-e)* were chosen for further analysis (Supplemental Table 2). The three other clades (HIL), (FK), and (JGM) were targeted in separate experiments using combinations of specific crRNAs for each gene. As biochemistry experiments suggested a pure enzymatic role for PpKAI2L-H, a deletion mutant was obtained through homologous recombination, where the full CDS was removed from the moss genome *(Δh* mutant, Figure 8 and Supplemental Figure 12A). This *Δh* mutant was employed in further transformation experiments with crRNAs from the same (HIL) clade and/or from clade (FK) and (JGM), generating *△h-i-l and △h-f-k-j* mutants (Supplemental Table 2). Eventually, a 7X mutant was obtained *(△h-i-f-k-j-g-m)* where all the mutations except those in *PpKAI2L-J* and *-M* were null (Supplemental Table 2 and see below).

**Figure 8:**
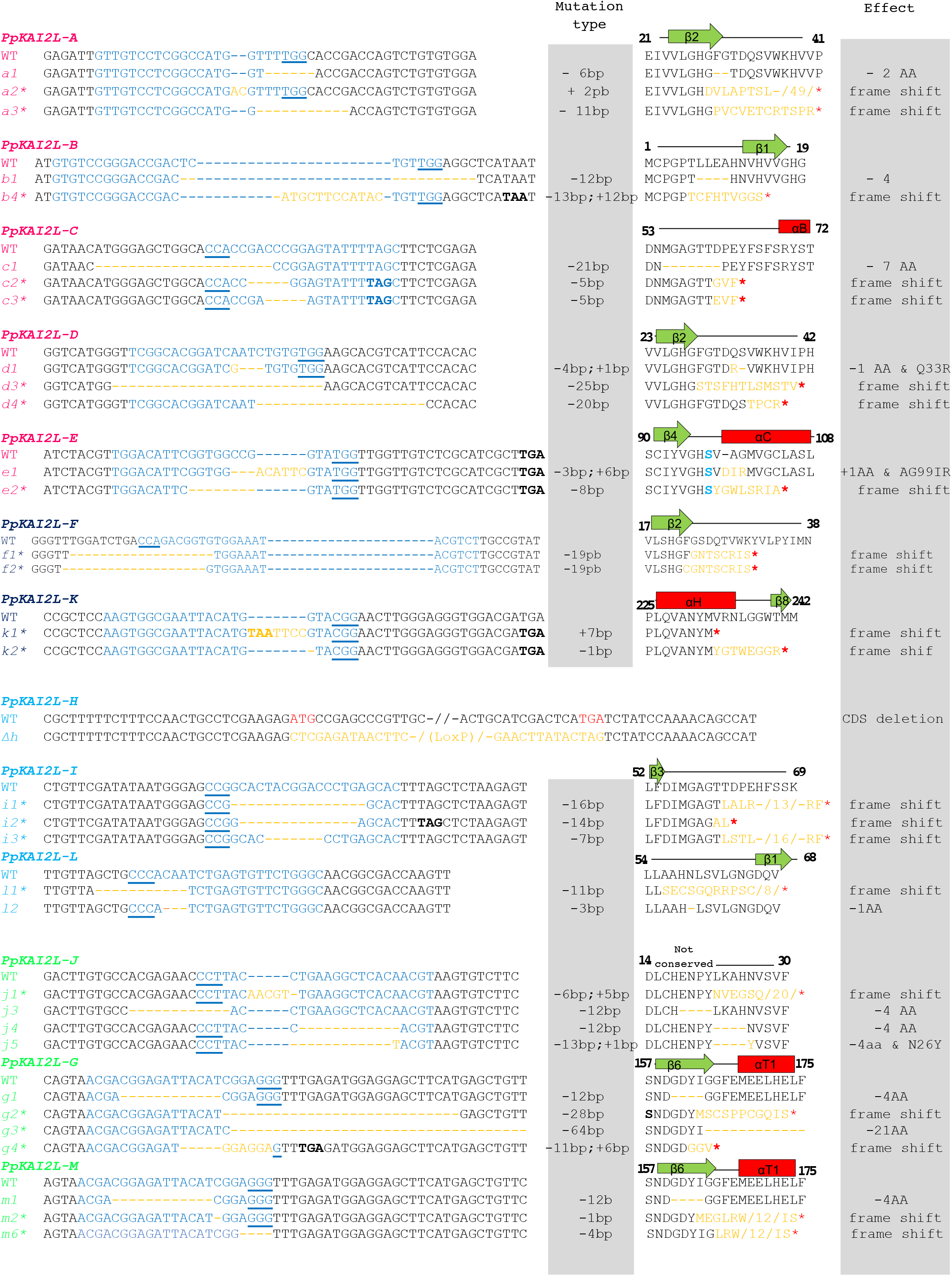
Mutations obtained in all 13 *PpKAI2L* genes. For all 13 *PpKAI2L* genes, WT nucleotide and protein sequences are shown above the altered sequences found in the generated CRISPR-Cas9 mutant lines (in italics, numbered). The number of the first shown amino acid (aa) and the predicted secondary structure are indicated above the WT protein sequence. The crRNA sequence is shown in blue, with the PAM site underlined. Deletions are shown as dashes, insertions are noted with orange letters. Mutation type is detailed on the right. Premature STOP codons are noted in bold, with a red star on the aa sequence. On protein sequences, the number of masked aa is noted between slashes. For *PpKAI2L-E,* the serine (S) of the catalytic triad is noted in bold blue. For *PpKAI2L-H*, a deletion of the full coding sequence between the ATG and STOP was obtained through homologous recombination; the use of CRE recombination led to 46 residual nucleotides, (not shown) corresponding to the LoxP site (see Methods). See Supplemental Table 2 for the list of mutants carrying one or several of the shown mutations. Predicted knock out mutations due to premature STOP codons or a large deletion induced in the mutant sequences are noted with an *.

### The *Ppkai2L* clade (A-E) quintuple mutants phenocopy *Ppmax2-1* in white light, while mutants in other clades are more similar to WT or *Ppccd8*

Our rationale was that a mutant affected in the response to PpCCD8-derived compounds should show a phenotype similar to that of the *Ppccd8* synthesis mutant. We first performed a phenotypic analysis of the mutants in light conditions. After four weeks of culture, *Ppccd8* mutant plants were slightly bigger than WT ((Proust et al. 2011), Figure 9), while the *Ppmax2-1* mutant was smaller, with fewer but bigger gametophores ((Lopez-Obando et al. 2018), Figure 9). The diameter of mutants in clade (A-E) genes was significantly smaller than that of *Ppccd8* and WT, and slightly larger than that of *Ppmax2-1* (Figure 9A-B and Supplemental Figure 12B). The phenotype of the clade (A-E) mutants, with early and large gametophores resembled that of the *Ppmax2-1* mutant although not as pronounced (Figure 9A). To the naked eye, three week-old plants from mutants of all three clades (FK) (JGM) and (HIL) genes were indistinguishable from WT (Figure 9A). These observations were also true for very young plants (10 day-old, Supplemental Figure 12B). After a month of growth however, all mutants affecting genes from clade (FK) and (JGM) showed a slightly larger diameter, intermediate between that of WT and *Ppccd8.* The triple *Ppkai2L j3-g3*-m1* mutant was even larger than *Ppccd8* (Figure 9D). Mutants in clade (HIL) such as *Ppkai2L △h, Ppkai2L △h-i2\** and *Ppkai2L △h-i3*-l1* looked the same as WT (Figure 9D and Supplemental Figure 12B).

**Figure 9:**
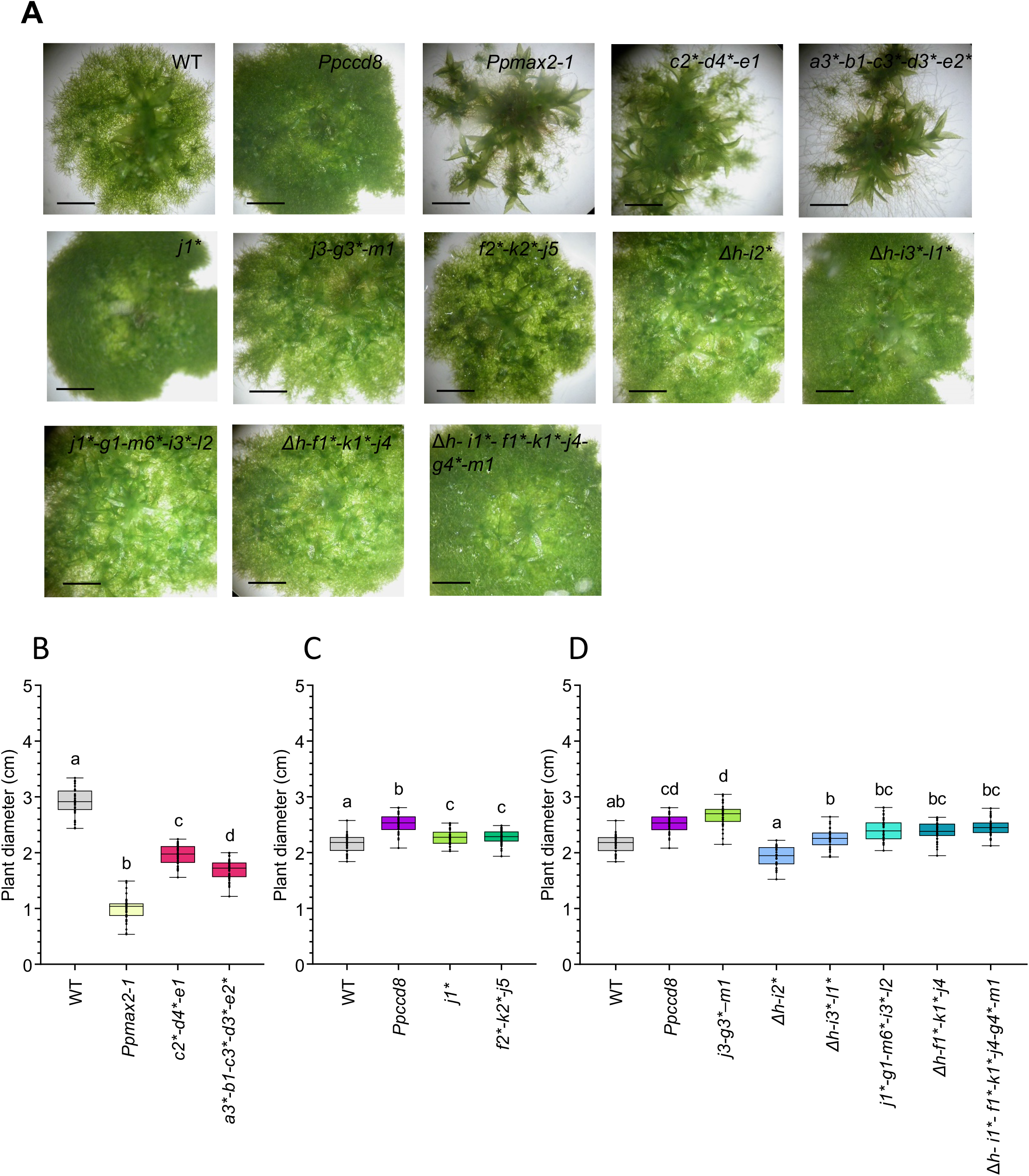
Phenotype of the *Ppkai2L* mutants in light. Plant diameters were measured after a ~ 4-week growth in the light. **(A)** 3-week-old plants. Scale bar = 2 mm **(B)** and **(C)**: 30-day-old plants, n > 40; (D): 28-day-old plants, n = 30. **(B)**, **(C)** and **(D):** All plants were grown on cellophane disks. Letters indicate statistical significance of comparisons between all genotypes (Kruskal-Wallis test followed by a Dunn *post hoc* test (p < 0.05)). Mutations are explained in Figure 8 and Supplemental Table 2. Asterisks indicate null mutations.

### Like *Ppmax2,* clade (A-E) quintuple mutants are affected in photomorphogenesis

Mutants in clade (A-E) genes showed the typical phenotype of the *Ppmax2-1* mutant in white light. We previously showed that the *Ppmax2-1* mutant is affected in photomorphogenesis under red light (Lopez-Obando et al. 2018). After two months growing under red light, *Ppmax2-1* gametophores were much more elongated than WT gametophores, whereas *Ppccd8* mutant gametophores were shorter (Figure 10). Among clade (A-E) mutants, gametophores of both triple mutants *Ppkai2L a2*-b4*-c2\** and *Ppkai2L c2*-d4*-e1* were a similar height to WT. Interestingly, the quintuple mutant *(Ppkai2L a1-b1-c1-d1-e2*)* showed significantly elongated gametophores, similar to *Ppmax2-1* (Figure 10 A and C). The other tested quintuple mutant *(Ppkai2L a3*-b1-c3*-d3*-e2*)* also had elongated gametophores under red light, intermediate between WT and *Ppmax2-1* (Supplemental Figure 13). The weak phenotype of both triple mutants suggests a functional redundancy among clade (A-E) genes, as KO mutations for PpKAI2L-A, B, C and/or D did not result in as elongated gametophores as in the *Ppmax2-1* mutant.

**Figure 10:**
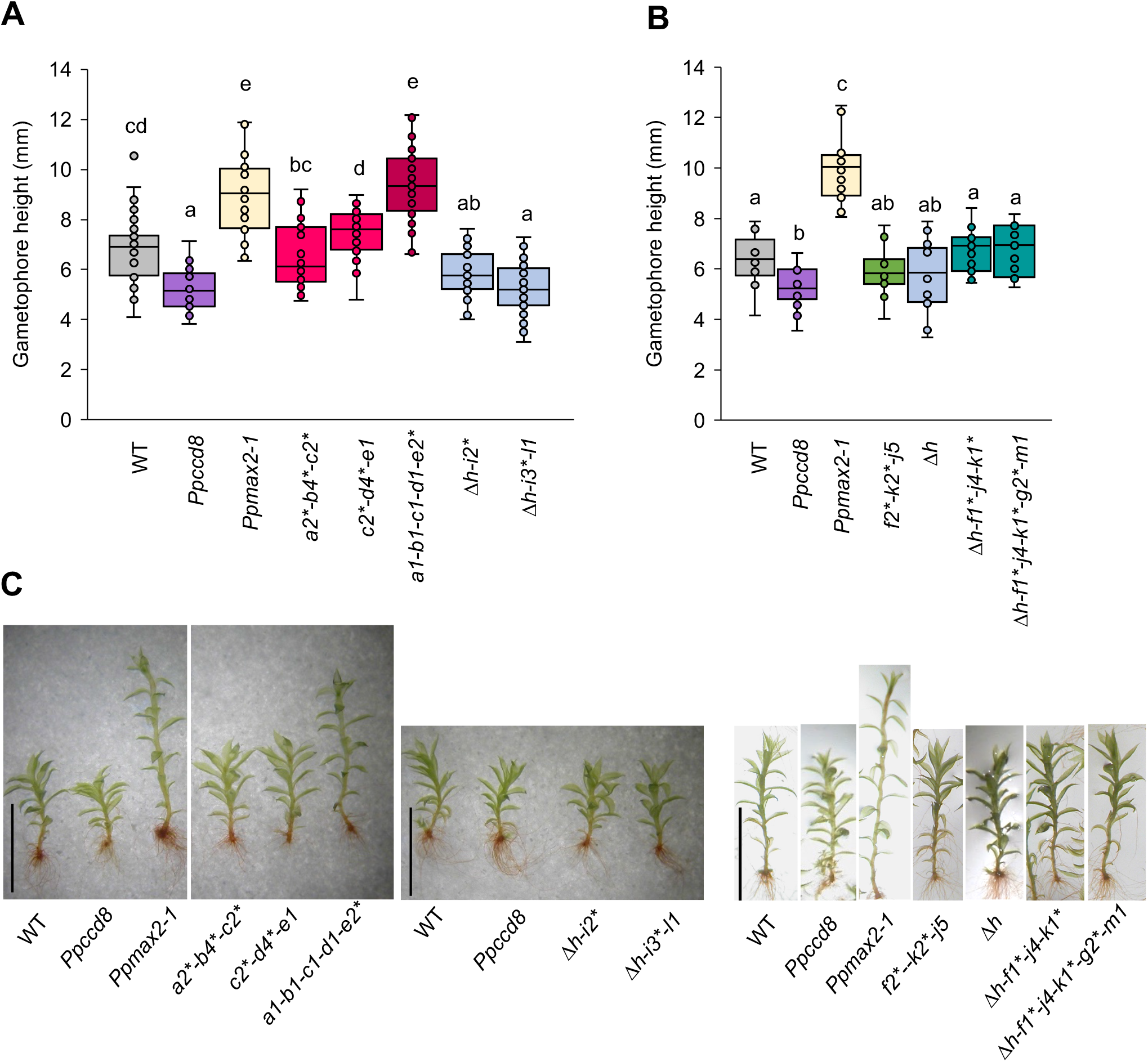
*Ppkai2L* mutant gametophores in red light. **(A)** Gametophore height of *Ppkai2L* mutants affecting clade (A-E) genes *(a2*-b4*-c2*; c2*-d4*-e1; al-bl-cl-dl-e2*)* and clade (HIL) genes *(Δh-i2\** and *△h-i3*-l?),* compared to that of WT, *Ppccd8* and *Ppmax2-1* mutants, following two months growth under red light. Mutations are explained in Figure 8 and Supplemental Table 2. Asterisks indicate null mutations. Box plots of n = 32-36 gametophores, grown in 3 Magenta pots, harboring between 15 and 25 leaves. Statistical groups (all genotypes comparison) are indicated by letters and were determined with a Kruskal-Wallis test followed by a Dunn *post hoc* test (p <0.05). **(B)** Gametophore height of *Ppkai2L* mutants affecting clade (HIL) gene (*Δh*), both clades (FK) and (JGM) *(f2*-j5*-k2*)* and all three clades (HIL) (FK) and (JGM) *(Δh-f1*-j4-k1\** and *△h-f1*-j4-k1*-g2*-m1).* Mutant genotypes carry mutations as indicated in Figure 8 and Supplemental Table 2, with asterisks for null mutations. Box plots ofn = 11-15 gametophores, grown in 3 Magenta pots, harboring between 15 and 25 leaves. Statistical groups (all genotypes comparison) are indicated by letters and were determined with a Kruskal-Wallis test followed by a Dunn *post hoc* test (p < 0.05). **(C)** Examples of gametophores following a 2-month growth under red light, from WT, *Ppccd8, Ppmax2-1,* and *Ppkai2L* mutants as shown in (A) and (B). Scale bar = 5 mm.

Gametophores from mutants where genes from clade (FK) and/or (JGM) were mutated *(Ppkai2L f2*-j5-k2*, Ppkai2L Δh-f1*-j4-k1\** and *Δh-f1*-j4-k1*-g2*-m1,* Figure 10B) were similar in height to WT, suggesting that genes from clade (FK) and clade (JGM) have no role in photomorphogenesis in red light. The clade (HIL) *Ppkai2L △h-i2*and Ppkai2L △h-i3*-l1* mutants showed shorter gametophores under red light, similar to *Ppccd8* (Figure 10A and 10C). In contrast, the gametophores of the single *Ppkai2L △h* mutant were intermediate in height between WT and *Ppccd8* (Figure 10B). This may suggest a specific role for the (HIL) clade genes, with opposite impact on gametophore development compared to that of the PpMAX2 pathway.

In conclusion, the phenotype of the *Ppkai2L* mutants in red light allowed the functions of the clade (A-E) genes to be differentiated. These are likely to be involved in a PpMAX2 dependent pathway, related to photomorphogenesis, whereas genes from the three other clades (DDK clade), are more likely to play a role independent from PpMAX2.

### Mutants in the (JGM) clade no longer respond to (+)-GR24 application

In the assays reported above, we measured the response of WT and *Ppccd8* mutant to SLs analogs (Figure 2). To determine which of the *Ppkai2L* mutants carry mutations in potential receptors for PpCCD8-compounds (SL-related) or other (KL-related) compounds, we used similar assays to test their phenotypic response to GR24 enantiomers at 0.1 μM (Figure 11).

**Figure 11:**
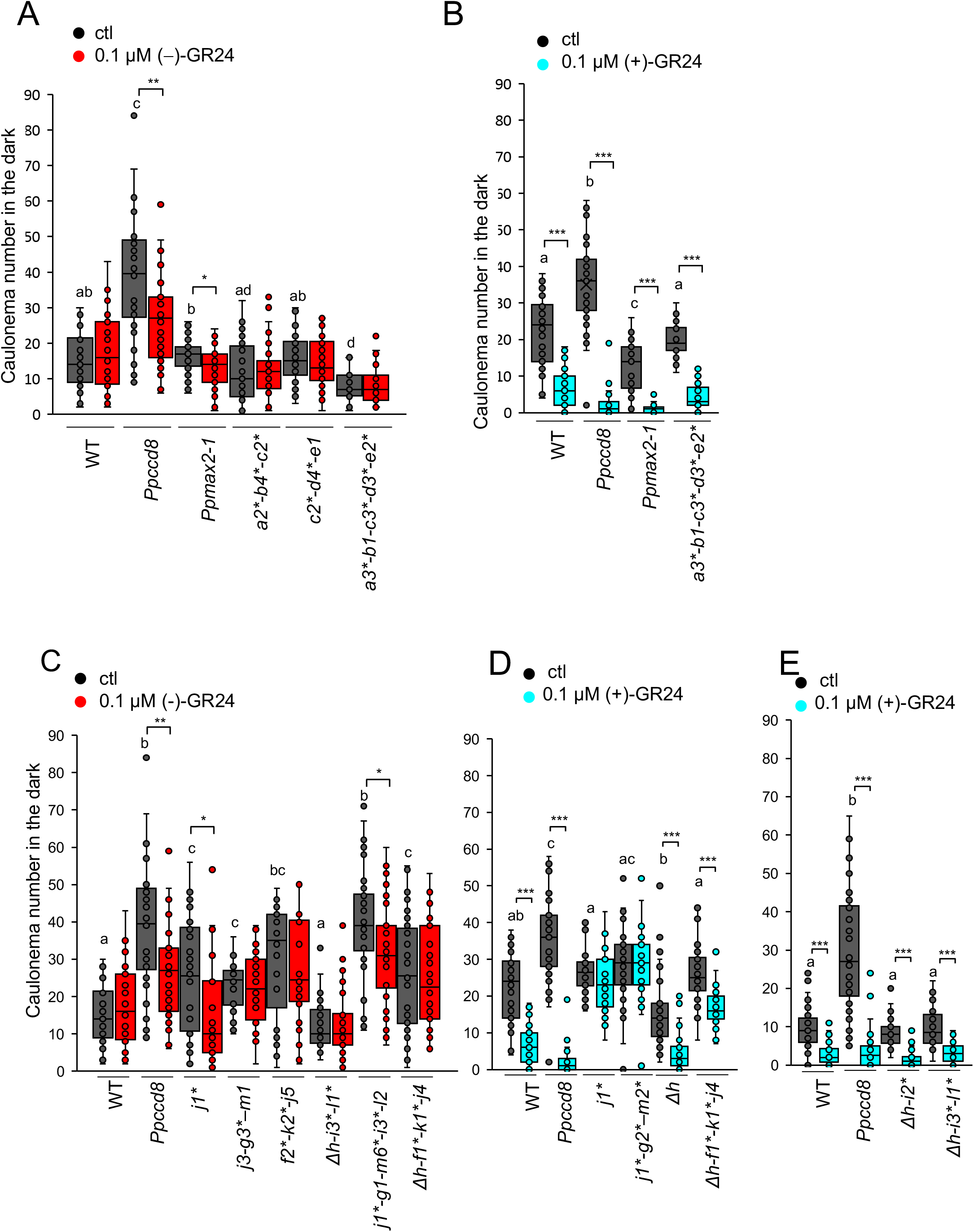
Phenotypic response of *Ppkai2L* mutants to (-)-GR24 and (+)-GR24 application: *caulonema* number in the dark. (**A-B**) *Caulonema* numbers from mutants affecting clade (A-E) genes following application of 0.1 μM (-)-GR24 (in red, **A**) or 0.1μM (+)-GR24 (in turquoise, **B)**. DMSO was applied as a control treatment (ctl, dark grey). WT and both the *Ppccd8* and *Ppmax2-1* mutants were used as control genotypes. (**C-E**) *Caulonema* numbers from mutants affecting clade (JGM) genes: *j1*; j3-g3*-m1; j1*-g2*-m2*;* clade (FK) and (JGM) genes: *f2*-k2*-j5, clade* (HIL) genes: *Δh, Δh-i2*, Δh-i3*-l1*,* clade (HIL) and (JGM) genes: *j1*-g1-i3*-l2,* or all 3 clades genes: *Δh-f1*-j4-k1*,* following application of 0.1μM (-)-GR24 (in red, **C**) or 0.1μM (+)-GR24 (in turquoise, **D, E**). 0.01% DMSO was applied as a control treatment (ctl, dark grey). WT and the *Ppccd8* mutant were used as control genotypes. Mutations are explained in Figure 8 and Supplemental Table 2. Asterisks indicate null mutations. For each genotype, *caulonema* were counted after two weeks in the dark, from 24 individuals, grown in three different 24-well plates. Statistical groups (comparing genotypes in control conditions) are indicated by letters and were determined by a one-way ANOVA with Welch test (95% Cl). Significant differences between control and treated plants within a genotype based on one-way ANOVA with Welch test (*** p < 0.001; ** p < 0.01; * p < 0.05).

For the (A-E) clade, under control conditions, all *Ppkai2L* mutants showed an equivalent number of filaments to WT, except for the quintuple mutant. The latter tended to have less filaments in control conditions, a similar phenotype to *Ppmax2-1* (Figure 11A-B and Supplemental Figure 14A). Addition of 0.1 μM (-)-GR24 did not have a significant effect on clade (A-E) mutants or WT (Figure 11A). However, in this assay, both the *Ppccd8* and *Ppmax2-1* mutants showed a significant decrease in *caulonema* filament number. In a separate experiment, a dose of 1 μM of (-)-GR24 had opposite effects on WT and *Ppccd8* filament number with an increase and decrease, respectively (Supplemental Figure 14A, and Figure 2 above), but had no significant effect on the *Ppmax2-1* mutant, although the same tendency towards a decrease was observed. At this higher dose, the quintuple *A-E* mutant showed a significant decrease in *caulonema* filament number, like *Ppccd8,* but in contrast to WT. Thus, similar to *PpMAX2* loss of function, mutating clade (A-E) *PpKAI2L* genes does not abolish a response to the (-)-GR24 enantiomer. Application of (+)-GR24 had a significant negative effect on the number of filaments for the quintuple *Ppkai2L a3*-b1-c3*-d3*-e2\** mutant, like for WT and the *Ppccd8* and *Ppmax2-1* mutants (Figure 11B). Thus, mutating any of the clade (A-E) *PpKAI2L* genes does not hamper the response to (+)-GR24, and therefore likely neither the response to CCD8-derived compounds.

We then tested the effect of GR24 enantiomers on *Ppkai2L* mutants from the three other clades. Strikingly, under control conditions, like *Ppccd8,* all the mutants had more filaments than WT, except for clade (HIL) mutants which tended to have less filaments (Figure 11C to 11E and Supplemental Figure 14). Both the single mutant *Ppkai2L-j1\** and the quintuple mutant *Ppkai2L j1*-g1-m6*-i3*-l2* showed a significant response to (-)-GR24 (less *caulonema),* as did *Ppccd8* (Figure 11C and Supplemental Figure 14A and 14B). No clear response to (-)-GR24 was seen in mutants with KO mutations in *PpKAI2L-F, -K, -H, -G, -M, -I* or -*L*, like for WT. Finally we examined the response to (+)-GR24 for mutants of the (FK) (JGM) and (HIL) clades (Figure 11D and 11E and Supplemental Figure 14C and 14D). The number of *caulonema* was clearly reduced in WT and *Ppccd8* (as shown above, Figure 2), and in mutants carrying the *Ppkai2L Δh* mutation, alone or in combination with *f*, k* i\** or *l\** null mutations. Thus clade (FK) and clade (HIL) genes do not play a role in the response to (+)-GR24. However, the response to the (+)-GR24 enantiomer was abolished in all mutants where the *PpKAI2L-J* gene is KO (Figure 11 D: *j1\** and *j1*-g2*-m2*,* and Supplemental Figure 14C and 14D). Interestingly, in the two lines where the *j* mutation is not null but the *PpKAI2L-G* gene is KO (7x and *j6-g5*-m1* mutants), the response to (+)-GR24 was also abolished (Supplemental Figure14C and 14D). Thus, from phenotypic assays on mutant *caulonema*, both the *PpKAI2L-J* and *-G* genes appear necessary for the response to (+)-GR24, and are therefore the best candidates for receptors to PpCCD8-derived SLs.

To confirm that PpKAI2L-J and -G are likely receptors for PpCCD8-derived SLs, we measured the transcript levels of SL responsive genes in the corresponding mutants (Figure 12). We previously reported that in WT and the *Ppccd8* mutant, *PpKUF1LA* gene transcript abundance increases 6 h after plant transfer onto medium containing 3 μM (±)-GR24, and that this response is enhanced in dark conditions (Lopez-Obando et al. 2018). We used this marker along with the *Pp3c6 15020* gene encoding a putative histidine kinase, and previously found to be upregulated by (±)-GR24 (our unpublished data). Using GR24 enantiomers, we confirmed that transcript levels for both genes increased following 1 μM (+)-GR24 addition in WT and *Ppccd8,* but not in *Ppmax2-1.* Strikingly, an increase in transcript levels following (-)-GR24 application was observed for both markers in the *Ppccd8* mutant, and for *PpKUF1LA* only in WT (Figure 12). In the quintuple mutant of clade (A-E), addition of (+)-GR24 but not (-)-GR24 increased *PpKUF1LA* and *Pp3c6 15020* transcript levels. In contrast, in the *Ppkai2L-j1\** mutant, transcript levels of both genes increased following (-)-GR24 addition, and was slightly increased (*PpKUF1 LA*) or unchanged (*Pp3c6_15020*) by (+)-GR24 application. In the *Ppkai2L j3-g3*-m1* mutant, the response marker transcript levels increased slightly *(PpKUF1LA}* or remained unchanged with (+)-GR24 addition, but were unchanged by (-)-GR24. Thus, the transcriptional response of the tested mutants confirms that clade (A-E) genes are not involved in the response to (+)-GR24, while this response is impaired in clade (JGM) mutants. Only the *Ppccd8* and *Ppkai2L-j1\** mutants showed a clear and significant transcriptional response, with both markers, to (-)-GR24 addition.

**Figure 12:**
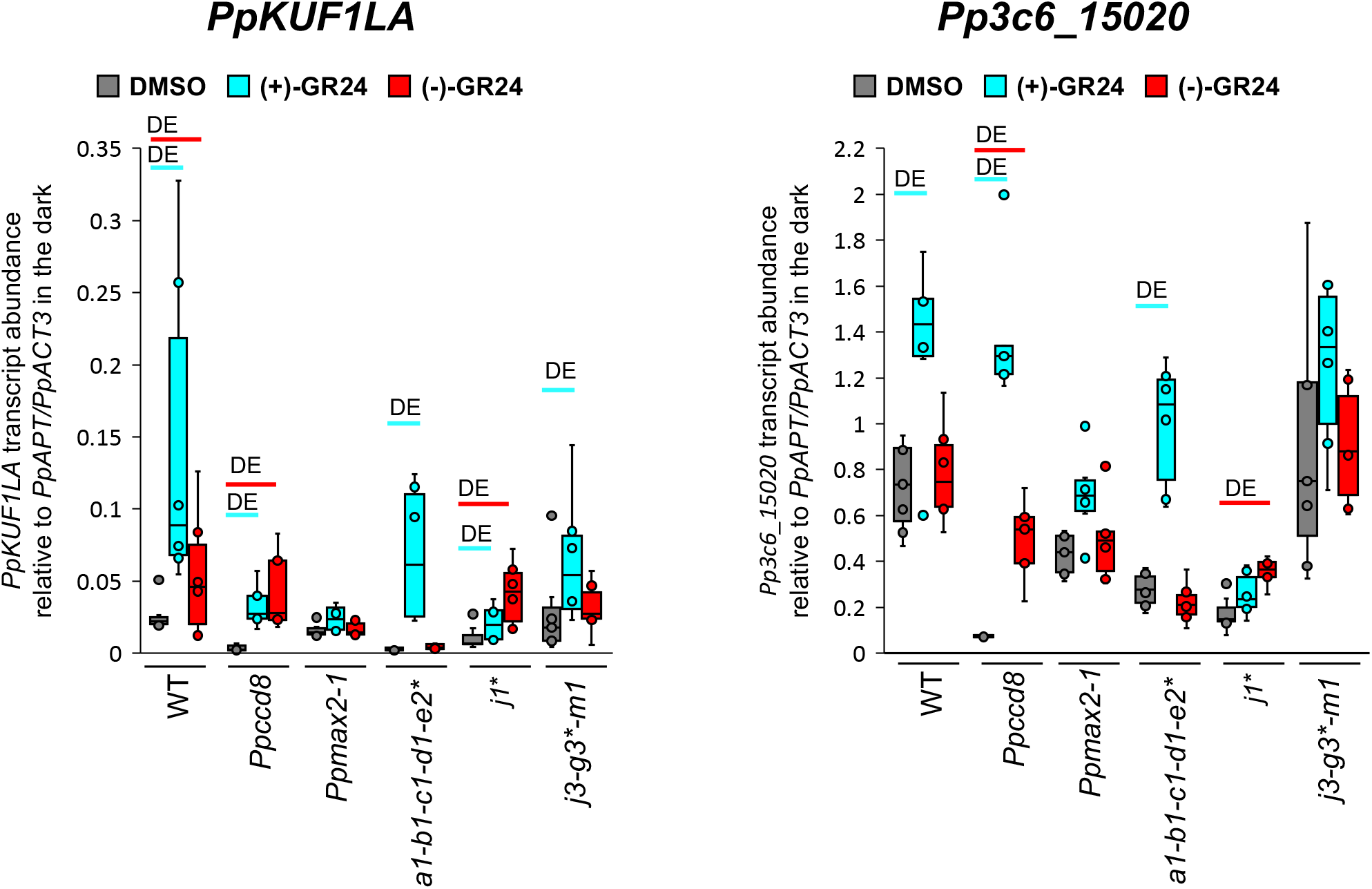
*Ppkai2L* mutant transcriptional response to (+)-and (-)-GR24. Transcript abundance analysis of the SL-responsive genes *PpKUF1LA* and *Pp3c6 15020,* in WT*, Ppccd8, Ppmax2-1* and *Ppkai2L* mutants *a1-b1-c1-d1-e2\** (clade (A-E)), *j1\** and *j3-g3*-m1* (clade (JGM)), after a 6-hour treatment in the dark with DMSO (control, grey), 1 μM (+)-GR24 (turquoise) or 1 μM (-)-GR24 (red). Box plots of at least four biological repeats are shown, relative to mean *(PpAPT-PpACT3}* transcript abundance. Two-fold differences in median values of transcript levels between control and treated plants are estimated as significant (DE) and noted in corresponding colors (turquoise for (+)-GR24 and red for (-)-GR24).

## DISCUSSION

### Are PpCCD8-derived products non-canonical SLs?

*P. patens* exudates were previously reported to induce the germination of *Orobanche ramosa* (old denomination of *Phelipanche ramosa*) seeds (Decker et al. 2017), but the origin of the tested population was not specified. Here we further demonstrated that PpCCD8-derived products are germination stimulants of *P. ramosa* group 2a seeds, harvested in a hemp field, but do not induce the germination of *P. ramosa* group 1 seeds, collected from an oilseed rape field. Differences in the suceptibility of root parasitic weeds can be attributed to the chemical nature of host plant exudates (Yoneyama et al. 2018b). Our results suggest that the PpCCD8-derived products share similarities with hemp secondary metabolites. So far, no known canonical SLs has been isolated from hemp (Huet et al. 2020). Since *P. patens* likely produces carlactone (Decker et al. 2017), but lacks a true *MAX1* homolog (Proust et al. 2011), we can hypothesize that PpCCD8-derived compounds correspond to non-canonical SLs, derived from carlactone or hydroxyl carlactones (Yoneyama 2020). Further supporting this hypothesis, we previously showed that GR5, a non-canonical SL analog, was as bioactive as (±)-GR24 on *P. patens caulonema* length (Hoffmann et al. 2014). As mimics of SLs in the present study, we used the (+)- and (-)-GR24 artificial analogs, available at the time of our study (see below). Of note, both isomers are active on *P. ramosa* group 1 and group 2a seeds. However, the (+)-GR24 isomer, which has a canonical SL structure, has similar germination stimulating activity to (±)- GR24, while the (-)-GR24 isomer is far less active (de Saint Germain et al. 2019). For future identification of PpCCD8-derived SLs, non-canonical SL analogs such as the recently described methyl phenlactonoates (Jamil et al. 2020) may be more appropriate.

### Looking for the best mimic of SLs or KL

The (-)-GR24 analog has a non-natural configuration that has never been found from plant exudates, contrary to the (+)-GR24 enantiomer, which bears similarity to 5-deoxystrigol ((+)- 5DS) and strigol-type canonical SLs (Scaffidi et al. 2014). In our bioassays of moss phenotypes, CL application decreased the number of *caulonema* of both WT and the *P. patens* mutant, in a dose-responsive manner. A similar (but much stronger) effect was observed with (+)-GR24, that we thus consider as so far the best mimic of *P. patens* CCD8-derived compounds. It is not surprising that (+)-GR24 is more potent than CL, as assays were carried out in a wet medium and natural SLs were described as being far less stable than synthetic analogs in aqueous medium (Akiyama et al. 2010; Boyer et al. 2012). Moreover, we cannot exclude that CL needs to be transformed in a more bioactive non-canonical SL *in planta* to trigger the effects observed here. In contrast, the effects of (-)-GR24 are weak, not dose responsive, and sometimes contradictory in WT versus the *Ppccd8* mutant. Indeed, in several assays, we observed a significant increase in *caulonema* number in WT (Figure 2, Supplemental Figure 14A) while this number consistently decreased in *Ppccd8,* mimicking the result of SL application (Figure 11, Supplemental Figure 14A and 14B). Interestingly, we also observed an increase in *caulonema* number when testing KAR2, although this increase was only significant at 10 μM for *Ppccd8.* So far we had not observed any effect of karrikins (KAR_1_) on *P. patens* phenotypes (Hoffmann et al. 2014), and this is thus the first hint of a possible effect of some karrikins on moss, which needs to be confirmed. We hypothesize that the increase of *caulonema* filament number is the effect triggered through the as yet unidentified moss KL (see also below and Figure 13). It is puzzling however that the effect of KAR2 is better seen in *Ppccd8* (thus in the absence of SLs), than in WT, while the same effect of (-)-GR24 (increasing of filament number) is only seen in WT. We propose (Figure 13 and see below) that clade (JGM) PpKAI2L proteins possibly perceive (-)-GR24, which would explain the apparent contrary effects of this enantiomer. In the absence of endogenous SL (in the *Ppccd8* mutant), the (-)-GR24 would thus trigger the SL pathway. Thus, (-)-GR24 is not a very robust mimic of the as yet unidentified moss KL, and KAR2 is likely not a potent KL mimic either. This conclusion also suggests that *P. patens* KL may be quite different from that of angiosperms.

**Figure 13:**
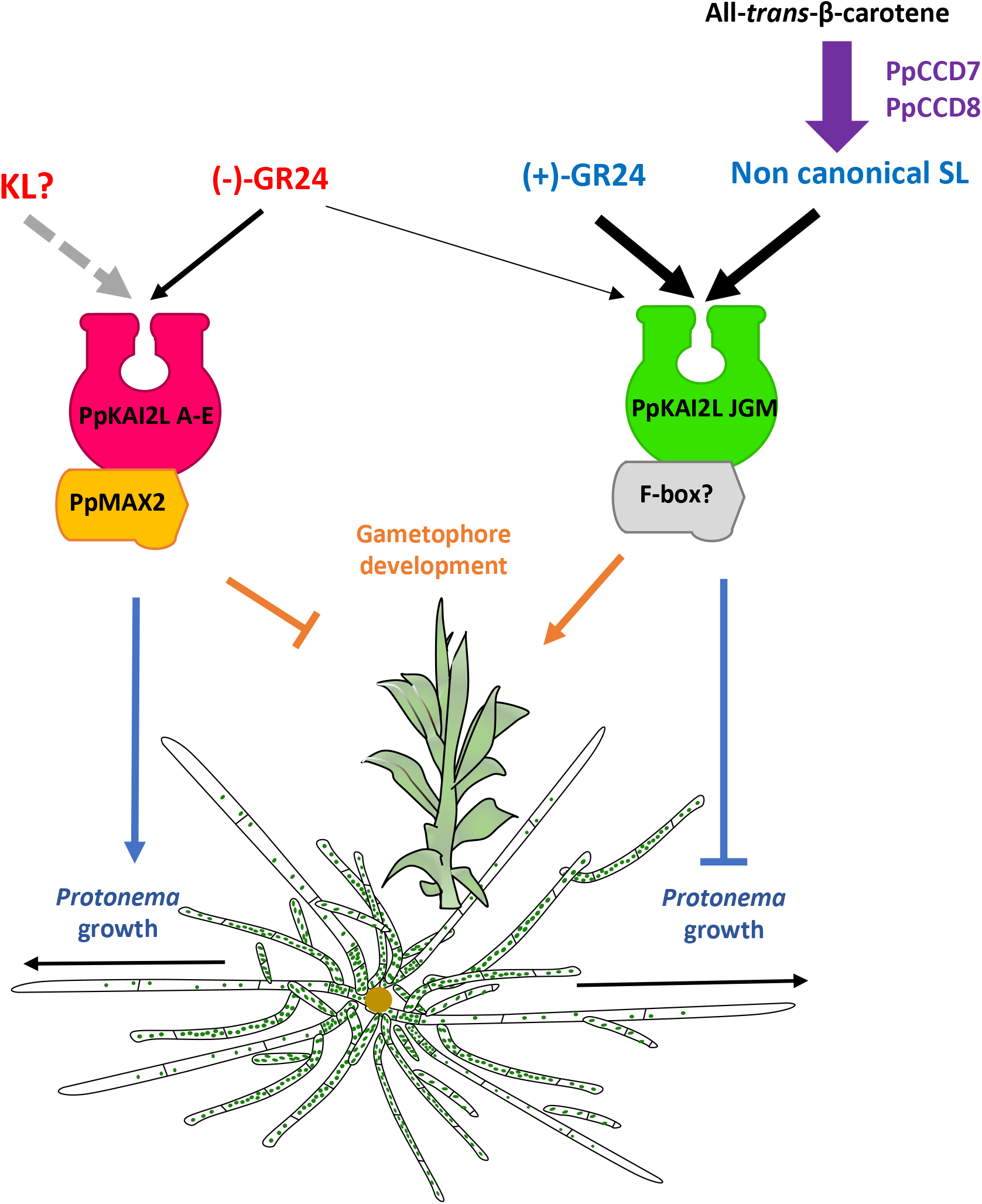
Current model for SL (PpMAX2-independent) and KL (PpMAX2-dependent) perception in *P. patens.* Clade (A-E) PpKAI2L proteins perceive KL compounds which promote *protonema* growth and inhibit gametophore development through a PpMAX2-dependent pathway. Clade (JGM) PpKAI2L proteins perceive PpCCD8-derived compounds (noted non-canonical SL) which inhibit protonema growth while promoting gametophore development, in a PpMAX2-independent manner. (+)-GR24 mimics PpCCD8- derived compound effects and is perceived by clade (JGM) PpKAI2L proteins. (-)-GR24 is perceived by both clade (A-E) and clade (JGM) PpKAI2L proteins, making it a poor mimic of moss KL. Grey doted arrow indicates non-demonstrated effect. Thickness of black arrows indicate the strength of the compound’s effect.

### Biochemical characterization of the PpKAI2L proteins suggests that PpKAI2L from the (A-E) subclade may act as KL receptors

In biochemistry experiments, among the analogs used, we observed the best binding of the (-)- GR24 enantiomer to AtKAI2 and all tested PpKAI2L proteins from the (A-E) clade. This confirms a recent report (Bürger et al. 2019) of preferential binding of (-)-5DS by PpKAI2L- C, -D and -E. In addition, we found that the (+)-GR24 enantiomer interacts poorly with these PpKAI2L proteins whereas it does not with AtKAI2, indicating less stringency for moss eu- KAI2 clade proteins. Still, this result suggests that clade (A-E) PpKAI2L proteins share a perception mechanism with AtKAI2, and furthermore may recognize KL-like compounds. The moss KL compound(s) may however differ from the angiosperm KL, since the expression of PpKAI2L-C or -D does not complement the *kai2-2* hypocotyl phenotype in Arabidopsis (see below). It should be noted that AtKAI2 is degraded following KAR perception, in a MAX2- independent manner (Waters et al. 2015), and this could be tested on clade (A-E) PpKAI2L proteins. As with the two other tested clades (FK) and (HIL), none of the interaction assays highlighted any preferential binding of GR24 isomers, despite the use of pure enantiomers. Indeed, only PpKAI2L-K showed similar binding affinity with both (-)-GR24 and (+)-GR24, but no stereoselectivity. Unfortunately, none of the PpKAI2L proteins from clade (JGM) could be purified for interaction assays, which was also reported by (Bürger et al. 2019). In the future, overexpression in *P. patens* or in other heterologous systems (yeast, insect cells) may be a solution for producing these proteins and carrying out biochemistry studies.

### PpKAI2L-H is the most efficient hydrolase among PpKAI2L proteins

The PpKAI2L-H protein shows strong cleavage activity towards all four GR24 stereoisomers as well as towards the synthetic probe (±)-GC242, compared to any other PpKAI2L protein, but also compared to the Arabidopsis AtKAI2 and AtD14 proteins. Mutating the L^28^ residue into a F is sufficient to reduce the efficiency of this activity (strong reduction of the *k*cat and of the *V*_max_), but has no effect on the *K*_1/2_ towards (±)-GC242. The efficient cleavage activity of PpKAI2L-H is therefore not likely due to a stronger affinity of this protein for the substrate. It has been hypothesized that the L^28^ residue (as the F^181^ residue), that is unique to PpKAI2L-H, does not particularly enlarge the pocket size of PpKAI2L-H (Bürger et al. 2019). Then, our results may highlight the ability of a residue to control exit of the cleavage reaction product. Indeed, the conserved F^28^ residue in D14/KAI2 proteins may act as gate keeper and this could explain the single turnover kinetics observed with some SL analogs (de Saint Germain et al. 2016). We hypothesize that PpKAI2L-H high velocity may be linked to the lack of this gate keeper residue, allowing a high substrate turnover. The strong enzyme activity of PpKAI2L-H could have a specific role in plants, perhaps as a catabolic enzyme, to regulate the levels of bioactive signaling molecules (Seto et al. 2019).

When only the *PpKAI2L-H* gene is mutated (Δ*h* mutant), no striking phenotype is observed (Supplemental Figure 12B), and in particular, the phenotypic response to (+)-GR24, that mimics CCD8-derived SLs, is similar to that of WT plants (Figure 11D) but contrasts with the other mutants' response (see below). In red light however, the gametophores of Δ*h-i2* and△h- i3*-l1* mutants are less elongated than WT gametophores, which is also observed in *Ppccd8.* However, in *Ppccd8* this could be related to the higher number of filaments, leading to the initiation of more (but smaller) gametophores, whereas the number of filaments in the dark is not higher in clade (HIL) mutants. In addition, clade (HIL) mutants tend to have less filaments than WT (Figure 11D and 11E and Supplemental Figure 14A). To note, our results do not support a likely role for PpKAI2L-H in the KL pathway, despite KAR_1_ binding by this protein was reported (Bürger et al. 2019). Altogether, although quite tenuous, the mutant phenotypes could suggest that clade (HIL) genes undertake a specific role in *P. patens* development. The association of this role with PpKAI2L-H enzyme activity remains to be discovered.

### The Arabidopsis *Atd14-1 kai2-2* mutant complementation assay is more than anecdotal

Using the endogenous AtD14 promoter, we confirmed that expression of the PpKAI2L-C protein does not complement the Arabidopsis D14 function in rosette branching, as previously reported using the 35S promoter (Bürger et al. 2019). We can extend this observation to PpKAI2L-D, PpKAI2L-F and PpKAI2L-G, which were never previously tested for rosette branching complementation. Interestingly, one line expressing PpKAI2L-J did show partial complementation of branching that could be a hint of a SL perception function for this protein, or at least of possible interaction with SL signaling machinery. However, given the phenotype of *P. patens* mutants in clade (JGM), PpKAI2L-G expressing lines would have also been expected to complement the branching phenotype, unless the PpKAI2L-J is more active than its counterparts from the same clade, or somehow able to better interact with the Arabidopsis SL signaling factors. Using the endogenous AtKAI2 promoter, we also confirmed as observed by (Bürger et al. 2019), that PpKAI2L-C cannot complement the *kai2-2* mutation. Furthermore, we extend this observation to PpKAI2L-D and PpKAI2L-J. However, expressing moss PpKAI2L-G reduces the size of *Atd14-1 kai2-2* hypocotyls, suggesting that PpKAI2L-G may be able to perceive and transduce the endogenous KL signal, even though it does not respond to (-)-GR24. Strikingly, when expressed under the control of AtKAI2 promoter in the Arabidopsis At*d14-1 kai2-2* mutant, the AtD14 protein but also the moss PpKAI2L-C or -J proteins exacerbate the defect induced by the *kai2-2* mutation by leading to even more elongated hypocotyls. This suggests a putative interaction of these proteins with the Arabidopsis KAI2/KL pathway that should be further investigated. Still, it is clear that none of the PpKAI2L proteins fully complements the AtD14 or KAI2 function, probably because of defective interactions with AtMAX2 and/or other components of SL/KL pathways.

### Genetic analysis suggests that genes from the (A-E) clade are involved in the PpMAX2 dependent pathway

Mutant phenotypes clearly distinguish clade (A-E) from the three other clades. Indeed, the quintuple PpKAI2L-A to -E mutant shows a phenotype in white light quite similar to that of *Ppmax2-1.* It also has elongated gametophores under red light, and a low number of *caulonema* filaments in the dark, suggesting that clade (A-E) PpKAI2L and PpMAX2 proteins could be members of the same pathway (Figure 13). As PpKAI2L proteins from the (A-E) clade preferentially bind the (-)-GR24 enantiomer, we expected the mutants in this clade to be blind to (-)-GR24 application. This is what we observed when transcriptional response markers were examined, with transcript levels remaining unchanged following (-)-GR24 application in both the *Ppmax2-1* and the quintuple (clade A-E) *Ppkai2L* mutant, but increasing in the *Ppccd8* mutant (but not in WT, Figure 12). However, as mentioned above, (-)-GR24 does not appear to be a perfect mimic of the unknown moss KL, and other transcriptional response markers need to be found, that would better reflect the moss KL response. As for the phenotypic response, application of 0.1 μM (-)-GR24 had no effect on clade (A-E) mutant *caulonema* number in the dark, or on WT, but significantly decreased both *Ppccd8* and *Ppmax2-1 caulonema* number (Figure 11A). Strikingly, a 1 μM concentration of (-)-GR24, that has the opposite effect on WT (increases *caulonema* number) and *Ppccd8* (decreases *caulonema* number), led to no response in the *Ppmax2-1* mutant, while the quintuple clade (A-E) mutant showed a significant decrease in *caulonema* number (Supplemental Figure 14). Thus the quintuple (clade A-E) mutant is still able to perceive (-)-GR24, as well as the *Ppmax2-1* mutant. As said above, we hypothesize that other PpKAI2L proteins (presumably the (JGM) clade, see below) may perceive the (-)-GR24, triggering the PpCCD8-derived compounds pathway (Figure 13). This does not rule out the hypothesis that clade (A-E) PpKAI2L and PpMAX2 proteins are in a same pathway. In addition, (-)-GR24 was reported to have a dual effect, promoting both the KAI2 and the D14 pathways in Arabidopsis roots (Villaecija-Aguilar et al. 2019).

Finally, both phenotypic response in the dark and transcriptional response to the (+)-GR24 enantiomer are unaffected in the clade (A-E) *Ppkai2l* mutants, indicating that clade (A-E) PpKAI2L proteins are not receptors for PpCCD8-derived compounds.

### PpKAI2L- J, -G, -M mediate PpCCD8-derived (SL-related) responses

In white light conditions (Figure 9), mutants affecting clades other than (A-E) showed phenotypes either similar to WT (mutants in clade (HIL) genes), or intermediate between WT and *Ppccd8* (mutants in clade (FK) and (JGM) clades). In the dark in control conditions (Figure 11), the *caulonema* number of mutants in clades (FK) and (JGM) is also intermediate between that of WT and *Ppccd8*, while it is similar to WT (or slightly smaller) in clade (HIL) mutants. Based on the hypothesis that synthesis and response mutants show similar phenotypes, genes from clades (FK) and (JGM) are thus the best candidates for PpCCD8-derived compound receptors. When the phenotypic response of these mutants to (+)-GR24 application was examined, plants with KO mutations in *PpKAI2L-J* or *PpKAI2L-G/M* (Figure 11, *j1*,* and *j1*- g2*-m2*,* and Supplemental Figure 14, *j7*-g1–m1, j8*-g1 -m5*,* and *j6-g5*-mł)* no longer responded to this compound. In contrast, both *Δh-f1*-k1*-j4 andΔh-f3*-k3*-j6* mutants, show a significant response to (+)-GR24 application (Figure 11 and Supplemental Figure 14), indicating that KO mutations in both *PpKAI2L-F* and *PpKAI2L-K*, or deletion of *PpKAI2L-H* do not abolish the response to the CCD8-derived mimic compounds, not even additively. The absence of a response in higher order mutants where either *PpKAI2L-J* or *PpKAI2L-G* are KO confirms the prominent role of both genes in the response to (+)-GR24. However, if, as expected, transcript levels of the *Pp3c6 15020* response marker gene did not change in either *j1\** and*j3-g3*-m1* mutants following (+)-GR24 application (Figure 12), transcript levels of the *PpKUFILA* gene increased in both mutants, suggesting a response to the SL analog. Thus, while the KO mutation of either *PpKAI2L-J* or *PpKAI2L-G* is sufficient to abolish the phenotypic response in the dark, it does not completely abolish the transcriptional response to (+)-GR24. Presumably, mutation of both *PpKAI2L-J* and *-G* or even all three genes *(-J-G-M)* is necessary to fully impair this response. Another possibility is that the transcriptional markers, first identified using (±)-GR24 (Lopez-Obando et al. 2016a; Lopez-Obando et al. 2018) are not fully specific for assaying the response to enantiomers. This result could also suggest that the transcriptional response, which is assessed far earlier than the phenotypic response (6 hours versus 15 days), is perhaps more sensitive to a very slight activation of the PpCCD8-derived SL pathway by PpKAI2L proteins.

### PpKAI2L proteins are likely receptors in two separate pathways, one which is dependent and one which is independent from the PpMAX2 F-box protein

Our previous results on the PpMAX2 F-box protein indicated that, in contrast to its homolog in flowering plants, it is not involved in the response to CCD8-derived compounds (Lopez-Obando et al. 2018). Like MAX2 in flowering plants however, PpMAX2 plays a role in early gametophore development and photomorphogenesis. We suggested that PpMAX2 could play a role in the moss KL signalling pathway, but we lacked evidence for other actors in this pathway in *P. patens*. The present study suggests that PpKAI2L-A to -E are α/β hydrolases involved in the same pathway as PpMAX2, since mutating these genes resulted in similar light-related phenotypes as those of the *Ppmax2* mutant. Specific mimics for the moss KL are however still missing for further evidence that PpKAI2L-A-E are receptors of the moss KL (Figure 13). Still, these results are consistent with the view that the KL pathway is ancestral relative to the SL pathway, and that the ancestral role of MAX2 in the land plants lineage is the transduction of the KL signal (Bythell-Douglas et al. 2017; Walker et al. 2019). Such a KL pathway has been very recently reported in *M. polymorpha*, further supporting this view (Mizuno et al. 2020).

PpKAI2L-J and PpKAI2L-G proteins are likely receptors of PpCCD8-derived compounds, which we suspect to be non-canonical SLs. Strikingly, these receptors are not particularly more similar to D14 than other PpKAI2L proteins. As hypothesized earlier (Lopez-Obando et al. 2016a; Bythell-Douglas et al. 2017), the expansion of the PpKAI2L family in *P. patens* (and not in other bryophytes such as *Marchantia polymorpha*, that contains two *MpKAI2* genes), as in parasitic angiosperms (Conn et al. 2015; Toh et al. 2015), may have allowed the emergence of SL sensitivity (de Saint Germain et al. 2021). Neofunctionalization of additional KAI2 copies in *P. patens* ancestry towards SL perception is therefore a possible explanation for our findings in this moss and would reveal a convergent evolution process, relative to the emergence of D14 in seed plants. We can also imagine that these neofunctionalized PpKAI2L lost the ability to interact with MAX2 in *P. patens*, and established a different protein network that potentially integrates new factors such as an alternative F-box (Figure 13). The remaining question is therefore to determine how SL signal transduction is achieved downstream of perception by clade (JGM) PpKAI2L proteins.

Consequently, the search for proteins that interact with *P. patens* KL and PpCCD8-derived compound receptors should be a priority in the near future. In particular, since SMXL proteins are key members of both the SL and KL pathways in flowering plants (Soudappan et al, 2015, 2017; Wang et al, 2015; Khosla et al, 2020), specific involvement of PpSMXL homologs (four genes) is currently under investigation.

Five more PpKAI2L proteins are present and expressed in *P. patens* (Lopez-Obando et al, 2016), and mutant analyses indicate that those are neither KL nor CCD8-derived compound receptors. Three of them however, from the (HIL) clade, among which the efficient hydrolase PpKAI2L-H, are likely involved in *P. patens* development, through pathways that remain to be discovered.

## METHODS

### Plant Materials and Growth Conditions

The *Physcomitrium* (*Physcomitrella*) *patens* Gransden wild-type (WT) strain was used and grown as previously described (Hoffmann et al. 2014; Lopez-Obando et al. 2018) in long day (16h) conditions. Unless otherwise stated in legends, experiments were always carried out on PpNO_3_ medium (minimal medium described by Ashton et al., 1979), in the following control conditions: 25°C during the day and 23°C at night, 50% humidity, quantum irradiance of ~80 μmol/m^2^/s. Tissues from young *protonema* fragments were multiplied prior to every experiment in the same conditions but using medium with higher nitrogen content (PpNH4 medium, PpNO_3_ medium supplemented with 2.7 mM NH_4_ tartrate). For red light experiments, plants were grown on PpNO_3_ medium in Magenta pots at 25°C, in continuous red-light (~45 μmol μmol/m^2^/s).

### Germination assays on root parasitic plant seeds

*P. patens* WT plants were grown on PpNH4 plates with cellophane disks for two weeks then the plants (and cellophane) were transferred onto PpNO_3_ medium with low phosphate (Phosphate buffer was replaced with 1g/L of MES buffer and the pH adjusted to 5.8) for another two weeks. *P. patens* exudates were collected by transferring the plants (still on cellophane disks) onto plates containing 10 mL distilled water and incubating them in the growth chamber with gentle agitation. After 48h, exudates were pipeted and filtered (0.2 μm). Exudates were diluted twice prior to testing their germination stimulating activity on preconditioned seeds of parasitic plants, as described previously (Pouvreau et al. 2013). Distilled water was used as a control. For germination assays on plates (Figure 1B-C), WT and the *Ppccd8* mutant were cultivated as above, and *P. ramosa* seeds were placed onto the plates after 10 days of phosphate starvation. Seeds were counted on three plates, with 7-10 microscope fields per plate.

### CRISPR-Cas9 mediated mutagenesis and homologous recombination

*P. patens* mutants were obtained as described in (Lopez-Obando et al. 2016b), using CRISPR- Cas9 technology. *PpKAI2L* coding sequences were used to search for CRISPR RNA (crRNA) contiguous to a PAM motif recognized by *Streptococcus pyogenes* Cas9 (NGG), using the webtool CRISPOR V4 against the *P. patens* genome Phytozome V9 (http://crispor.tefor.net/). Guide crRNAs were chosen in each *PpKAI2L* gene, preferably in the first exon, to ideally obtain the earliest nonsense mutation possible. When no guide could be designed in the first exon, it was alternatively chosen to recognize a region in close proximity to the codon for one of the last two residues of the catalytic triad. (Figure 1B). The same crRNA was used to target *PpKAI2L-G* and -*M*. Small constructs containing each crRNA fused to either the proU6 or the proU3 snRNA promoters (Collonnier et al. 2017) between the attB1/attB2 Gateway recombination sequences, were synthesized by Twist Biosciences. These inserts were then cloned into pDONR207 vectors (Invitrogen). Polyethylene glycol–mediated protoplast transformation was performed with multiple pDONR207-sgRNA as described previously (Lopez-Obando et al. 2016b). Mutations in the *PpKAI2L* genes were confirmed by PCR amplification of *PpKAI2L* loci around the recognition sequence of each guide RNA, and sequencing of the PCR products.

The PpKAI2L-H (Δ*h*) deletion mutant was obtained through homologous recombination. The full coding sequence of PpKAI2L-H from the ATG to stop was replaced with a resistance cassette. A 550 bp *PpKAI2L-H* 5' CDS flanking sequence was cloned into the pBNRF vector (Thelander et al. 2007) cut with BstBI/XhoI. Then a 500 bp *PpKAI2L-H* 3' CDS flanking sequence was cloned into the *BNRF-PpKAI2L-H* 5' construct digested with BcuI, so that the kanamycin resistance cassette of the vector was flanked by the PpKAI2L-H 5' and 3' sequences. *P. patens* WT protoplasts were transformed with the resulting construct as described previously (Lopez-Obando et al. 2016b) and transformants selected on 50mg/L Geneticin/G418. Transient expression of the CRE recombinase (Trouiller et al. 2006) in a confirmed transformant removed the resistance cassette to obtain the *Ppkai2L-Δh* mutant, as described in Figure 8 and Supplemental Figure 12.

### Phenotypic assays on *P. patens*

Analysis of *caulonema* growth in the dark was performed in 24-well plates, starting from very small pieces of protonema. These were grown for ~ two weeks in control conditions before being placed vertically (± treatment) in the dark for ~10 days. A single picture of each well was taken using an axiozoom (Zeiss) with a dedicated program. *Caulonema* were counted and measured using ImageJ (http://imagej.nih.gov/ij/) (see also (Guillory and Bonhomme 2021)).

### Chemicals

Racemic and pure enantiomers of GR24 and the (±)-GC242 probe were produced by F-D Boyer (ICSN, France). Racemic CL was kindly provided by A. Scaffidi (University of Western Australia, Perth, Australia). Chemicals were diluted in DMSO or acetone as indicated in figure legends.

### RT-qPCR analysis

Freshly ground WT (Gransden) tissues were inoculated in Petri dishes of PpNO_3_ medium, overlaid with a cellophane sheet. *Protonema* tissues were harvested after six days, 10 days or 15 days of growth in long days conditions (see above). To obtain gametophores and spores, plants regenerated from spores were cultivated for two weeks then transferred to Magenta pots containing PpNO_3_ medium (nine plants per pot) and cultivated in the same conditions as above. Gametophores were harvested after 35 days. Both *protonema* and gametophore samples were flash frozen in liquid nitrogen and kept at −80°C until RNA extraction. At 35 days the remaining pots were transferred to short day conditions (15°C, 100% hygrometry, quantum irradiance of 15 μmol m^-2^ s^-1^) for approximately two months until capsule maturity. Capsules were sterilized (90% chlore and 10% pure ethanol) then rinsed with sterile water. Each of the four biological replicates consisted of 10-20 capsules from which spores were freed by mechanical disruption and separated from capsule debris by filtering through a 25μm nylon mesh. Spores were kept in sterile water, flash frozen in liquid nitrogen and stored at −80°C until RNA extraction. For all samples except spores, tissues were ground in liquid nitrogen using a mortar and pestle and RNA were extracted and subsequently treated with DNAse I using the Plant RNeasy Mini extraction kit (Qiagen) following the manufacturer’s instructions. Spores were recovered in 1mL TRIzol reagent (Invitrogen) and crushed manually using fine pestles. RNA was separated from cell debris and protein using chloroform and then precipitated with isopropanol and washed with 70% ethanol. The spore RNA pellets were dissolved in RLT buffer from the Qiagen Plant RNeasy Mini kit and treated with DNAse I on columns following the manufacturer’s instructions. 500 ng of each RNA sample was used for retro-transcription using the RevertAid H Minus Reverse Transcriptase from Thermo Fisher. Quality of obtained cDNA extracts was verified by semi-quantitative RT-PCR using the reference gene *PpAPT.* Quantitative RT-PCR reactions were carried out in 5μL using the SsoAdvanced Universal SYBR Green Supermix from BioRad and the following program on QuantStudio™ 5 (Thermo Fisher Scientific): initial denaturation at 95°C for 3 minutes, then 45 cycles of 95°C for 10 seconds and 60°C for 30 seconds. Using the CTi (for the genes of interest) and CTref (mean for the two reference genes) values obtained, relative expression (RE) was expressed as RE = 2-CT^i^/2-CTref.

### Constructs and generation of transgenic lines

The expression vectors for transgenic *Arabidopsis* were constructed using the MultiSite Gateway Three-Fragment Vector Construction kit (Invitrogen). All the PpKAI2L constructs were tagged with a 6xHA epitope tag at their C-terminus. Lines were resistant to hygromycin. The AtD14 native promoter (0.8 kb) and AtKAI2 native promoter (0.7 kb) were amplified by PCR from Col-0 genomic DNA and cloned into *pDONR-P4P1R*, using Gateway recombination (Invitrogen) (see Supplementary Table 2 for primers). AtD14 CDS and AtKAI2 CDS were PCR amplified from Col-0 cDNA, PpKAI2L CDS were PCR amplified from *P. patens* cDNA and recombined into *pDONR221* (Invitrogen). *6xHA* with a linker (gift from U. Pedmale) was cloned into *pDONR-P2RP3* (Invitrogen). The suitable combination of promoters, CDS and *6xHA* were cloned into the *pH7m34GW* final destination vectors using the three fragment recombination system (Karimi et al. 2007), and named pD14:CDS-6xHA or pKAI2:CDS- 6xHA. The *Arabidopsis Atd14-1 kai2-2* double mutant in the Landsberg background (gift from M. Waters) was transformed following the conventional dipping method (Clough and Bent 1998), with Agrobacterium strain GV3101. For all constructs, more than 12 independent T1 lines were isolated and between 2-4 representative single-insertion lines were selected in T2. Two lines per construct were shown in these analyses. The phenotypic analysis shown in Figure 7 and Supplementary Figure 10 was performed on segregating T3 homozygous lines.

### Arabidopsis hypocotyl elongation assays

Arabidopsis seeds were surface sterilized by consecutive treatments of 5 min in 70% (v/v) ethanol and 0.05% (w/v) sodium dodecyl sulfate (SDS) and 5 min in 95% (v/v) ethanol. Then seeds were sown on 0.25 X Murashige and Skoog (MS) media (Duchefa Biochemie) containing 1% agar, supplemented with 1 μM (+)-GR24, (-)-GR24 or with 0.01 % DMSO (control). Seeds were stratified at 4 °C (2 days in dark) then exposed to white light for 3 h, transferred to darkness for 21 h, and exposed to low light for 4 days at 21°C. Plates were photographed and hypocotyl lengths were quantified using ImageJ (http://imagej.nih.gov/ij/).

### Arabidopsis branching quantification

Experiments were carried out in summer in a greenhouse under long photoperiods (15-16 h per day); daily temperatures fluctuated between 18°C and 25°C. Peak PAR levels were between 700 and 1000 μmol m^-2^ s^-1^. Plants were watered twice a week with tap water. The number of rosette branches longer than 5 mm was counted when the plants were 40-days-old.

### Protein expression and purification

AtD14, RMS3 and AtKAI2, with cleavable GST tags were expressed and purified as described in (de Saint Germain et al. 2016). For PpKAI2L protein expression, the full length coding sequences from *P. patens* were amplified by PCR using cDNA template and specific primers (see Supplementary Table 1) containing a protease cleavage site for tag removal, and subsequently cloned into the pGEXT-4T-3 expression vector. For PpKAI2L-L, the N-terminal 47 amino acids were removed. The expression and purification of PpKAI2L proteins followed the same method as for AtD14 and AtKAI2.

### Site-directed mutagenesis

Site-directed mutagenesis of *PpKAI2L-H* was performed using the QuickChange II XL Site Directed Mutagenesis kit (Stratagene), performed on pGEX-4T-3-PpKAI2L-H (see Supplementary Table 1 for primers). Mutagenesis was verified by DNA sequencing.

### Enzymatic degradation of GR24 isomers by purified proteins

The ligand (10 μM) was incubated with and without purified RMS3/AtKAI2/PpKAI2L proteins (5 μM) for 150 min at 25 °C in PBS (0.1 mL, pH = 6.8) in the presence of (±)-1-indanol (100 μM) as internal standard. The solutions were acidified to pH = 1 by addition of trifluoroacetic acid (2 μL) to quench the reaction and centrifuged (12 min, 12,000 tr/min). The samples were then subjected to RP-UPLC-MS analyses. The instrument used for all the analyses was an Ultra Performance Liquid Chromatography system equipped with a PDA and Triple Quadrupole mass spectrometer Detector (Acquity UPLC-TQD, Waters, USA). RP-UPLC (HSS C18 column, 1.8 μm, 2.1 mm × 50 mm) was carried out with 0.1% formic acid in CH_3_CN and 0.1% formic acid in water (aq. FA, 0.1%, v/v, pH 2.8) as eluents [10% CH_3_CN, followed by linear gradient from 10 to 100% of CH_3_CN (4 min)] at a flow rate of 0.6 mL/min. The detection was performed by PDA and using the TQD mass spectrometer operated in Electrospray ionization positive mode at 3.2 kV capillary voltage. The cone voltage and collision energy were optimized to maximize the signal and was 20 V for cone voltage and 12 eV for collision energy. The collision gas was argon at a pressure maintained near of 4.5.10^-3^ mBar.

### Enzymatic kinetic assays

Enzyme assays with pro-fluorescent probes and *p-*nitrophenyl acetate were performed as described in (de Saint Germain et al. 2016), using a TriStar LB 941 Multimode Microplate Reader from *Berthold Technologies*.

### Protein melting temperature assays

#### Differential Scanning Fluorimetry (DSF)

experiments were performed on a CFX96 Touch™ Real-Time PCR Detection System (Bio-Rad Laboratories, Inc., Hercules, California, USA) as described in (de Saint Germain et al. 2016).

#### nanoDSF

Proteins were diluted in Phosphate buffer saline (PBS) (100 mM Phosphate, pH 6.8, 150 mM NaCl) to ~10μM concentration. Ligand was tested at the concentration of 200 μM. The intrinsic fluorescence signal was measured as a function of increasing temperature in a Prometheus NT.48 fluorimeter (Nanotemper™), with 55% excitation light intensity and 1 °C/minute temperature ramp. Analyses were performed on capillaries filled with 10 μL of respective samples. The intrinsic fluorescence signal was expressed as the 350 nm/330 nm emission ratio, which increases as the proteins unfold, and was plotted as a function of temperature. The plots show one of the three independent data collections that were performed for each protein.

### Intrinsic tryptophan fluorescence assays

Intrinsic tryptophan fluorescence assays and determination of the dissociation constant *K_D_* were performed as described in (de Saint Germain et al. 2016), using a Spark^®^ Multimode Microplate Reader from Tecan.

### Direct electrospray ionization - mass spectrometry of PpKAI2L proteins (ESI)-MS under denaturing conditions

Mass spectrometry measurements were performed with an electrospray Q-TOF mass spectrometer (Waters) equipped with the Nanomate device (Advion, Inc.). The HD_A_384 chip (5 μm I.D. nozzle chip, flow rate range 100-500 nL/min) was calibrated before use. For ESI-MS measurements, the Q-TOF instrument was operated in RF quadrupole mode with the TOF data being collected between m/z 400-2990. Collision energy was set to 10 eV and argon was used as the collision gas. Mass spectra acquisition of PpKAI2L proteins (50 μM) in 50 mM ammonium acetate in presence or without GR24 enantiomers (500 μM) was performed after denaturation in 50% acetonitrile and 1% formic acid. Mass Lynx version 4.1 (Waters) and Peakview version 2.2 (Sciex) software were used for acquisition and data processing, respectively. Deconvolution of multiply charged ions was performed by applying the MaxEnt algorithm (Sciex). The average protein masses were annotated in the spectra and the estimated mass accuracy was ± 2 Da. External calibration was performed with NaI clusters (2 μg/μL, isopropanol/H_2_O 50/50, Waters) in the acquisition m/z mass range.

### Homology modeling

Figure 6D showing superimposition model was prepared using PyMOL (DeLano Scientific) with the crystal structure of AtD14 (PDB ID 4IH4) and PpKAI2L-H (PDB ID 6AZD).

### Statistical analyses

Kruskal-Wallis, Mann-Whitney and *post-hoc* Dunn multiple comparisons tests (details in Figures legends) were carried out either in R version 3.6.3 or in GraphPad Prism version 8.4.2. When applicable, one-way ANOVA analyses were preferred (see Figures 11 and Supplemental Figure 14 and their legends). Unless otherwise defined, the statistical significance scores used were as follow: # 0.05≤p<0.1, * 0.01≤p<0.05, ** 0.001≤p<0.01, *** p<0.001. Same letters scores indicate that p≥0.05 (non-significant differences).

## ACKNOWLEDGEMENTS

The authors thank Adrian Scaffidi (University of Western Australia, Perth, Australia) for the gift of carlactone, and Mark Waters (University of Western Australia, Perth, Australia) for Arabidopsis *kai2-2* and *d14-1 kai2-2* mutants. We are grateful to Jean-Paul Pillot (IJPB) for precious help with Arabidopsis branching assays, and to Fabien Nogué (IJPB) for stimulating discussions. MLO thanks Eva Sundberg for her support on his SL/KL’s activities at SLU.

## AUTHOR CONTRIBUTIONS

SB, AdSG, ML-O and CR designed the project. ML-O, AG, F-DB, DC, BH, PLB, J-BP, AdSG and SB conducted experiments. AG, ML-O, F-DB, DC, PLB, J-BP, PD, CR, AdSG and SB analyzed the data. AG and SB wrote the manuscript, with essential contributions from ML-O, F-DB, PD, CR and AdSG.

