## Supplemental Figures for "The *Physcomitrium (Physcomitrella) patens* PpKAI2L receptors for strigolactones and related compounds highlight MAX2 dependent and independent pathways"

### Supplemental Figure 1

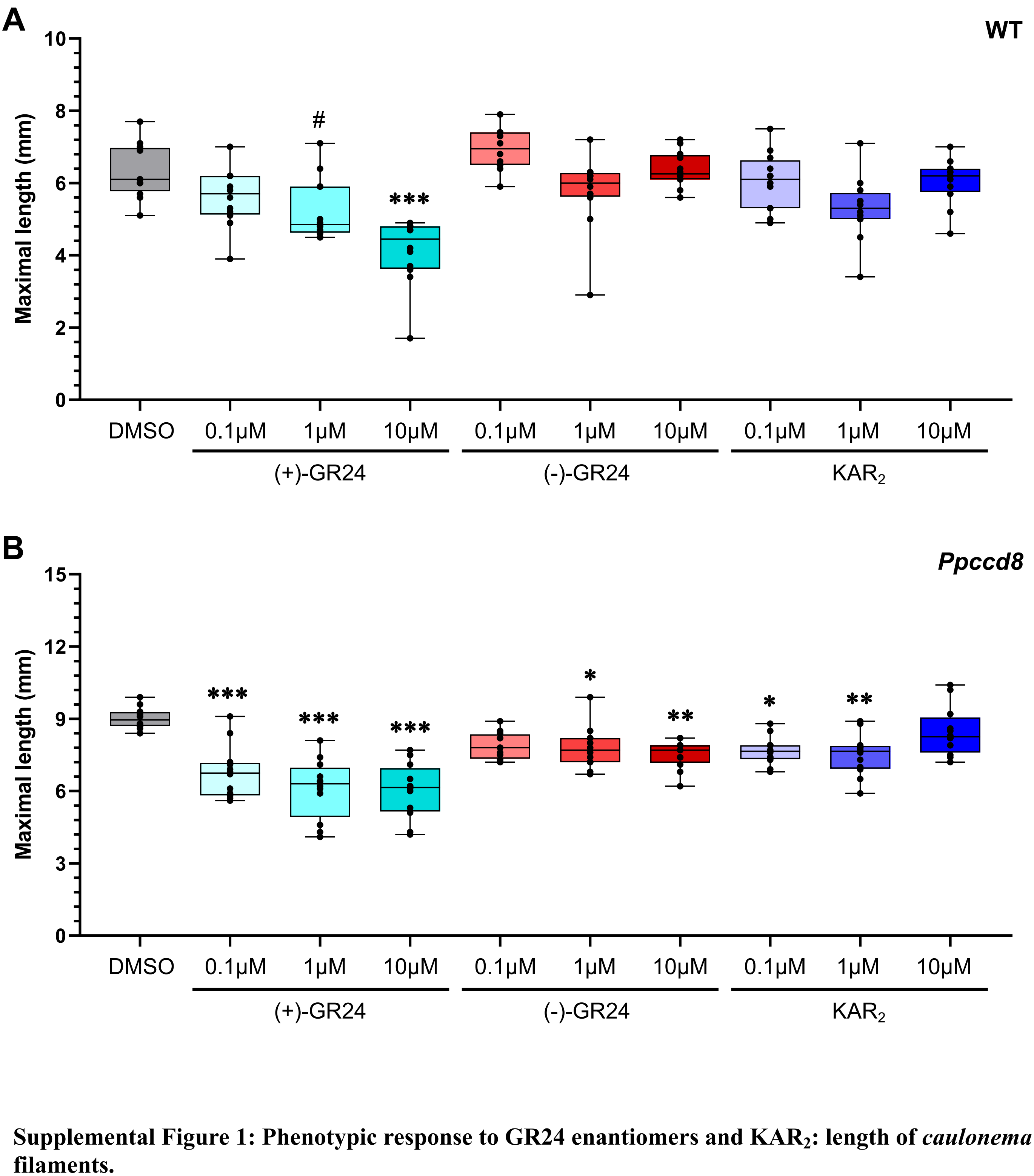


**Supplemental Figure 1: Phenotypic response to GR24 enantiomers and $KAR_2$: length of *caulonema* filaments.**

Maximal length of *caulonema* filaments was measured in WT (**A**) and the *Ppccd8* SL synthesis mutant (**B**) grown for 10 days vertically in the dark, following application of increasing concentrations of (+)-GR24 (turquoise), (–)-GR24 (red) or $KAR_2$ (blue). Control = DMSO. Significant differences between control and treated plants within a genotype are given, based on a Kruskal-Wallis test (Dunn *post-hoc*): ***, $p < 0.001$; **, $p < 0.01$; *, $p < 0.05$; # $p < 0.1$. For each genotype and treatment, n = 24 plants grown in three different 24 well-plates.

Supplemental Figure 2

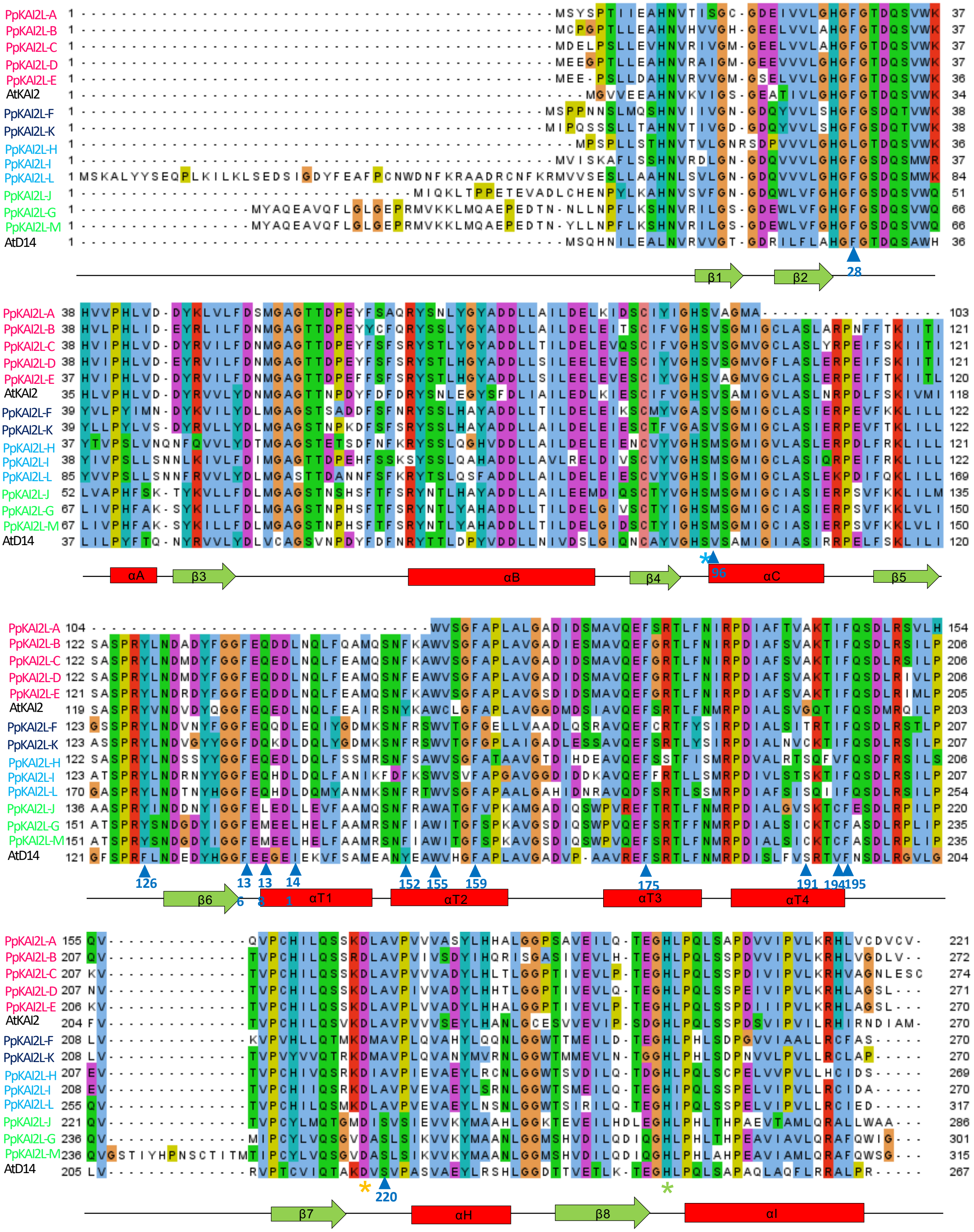

**Supplemental Figure 2: Sequence alignment of *Physcomitrium patens* (Pp) protein with AtD14 and AtKAI2 proteins from *Arabidopsis thaliana* (At).**
Protein sequences were aligned using CLUSTALX. The three amino acid residues of the catalytic triad are labelled with stars. Amino acid residues interacting with GR24 analogs in the binding pocket according to Zhao et al. (2015) are indicated with a blue arrowhead. Amino acid numbers are indicated following AtD14 numbering. The secondary structure assignment is based on the crystal structure of AtD14, and labels of the major strands and helices are based on Yao et al. (2016).

### Supplemental Figure 3

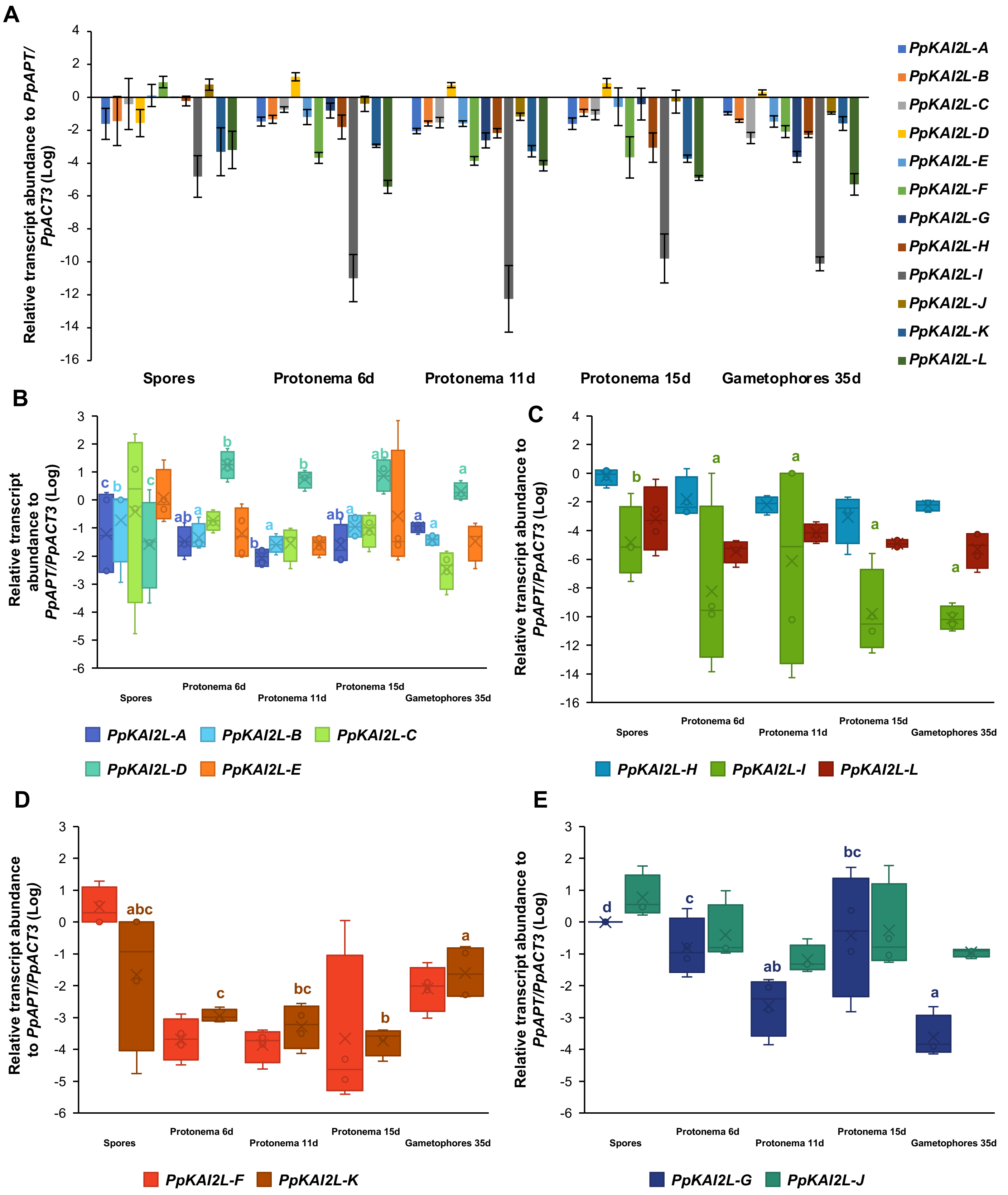

**Supplemental Figure 3: Expression of *PpKAI2L* genes throughout *P. patens* vegetative development.**

Relative expression is given as Log($2^{-Cti}/2^{-mCtref}$), where mCtref is the mean of Ct values for two reference genes (*PpAPT,* Pp3c8_16590 and *PpACT3,* Pp3c10_17080), for each biological replicate. Four biological replicates and two technical repeats were included in the analysis for each gene and tissue. For each technical repeat, normalization was carried out using the mean of the expression of the two reference genes. (**A**) Comparison of mean values (error bars represent standard errors) amongst all *PpKAI2L* genes. (**B-E**) Subclade-specific comparisons. When relevant, results of statistical analyses between tissues for a given gene (Kruskal-Wallis, $p < 0.05$) are indicated with bold letters. Samples recorded as *PpKAI2L-G* represent a mix of *PpKAI2L-G* and *PpKAI2L-M* transcripts, as their transcripts are almost identical.

### Supplemental Figure 4

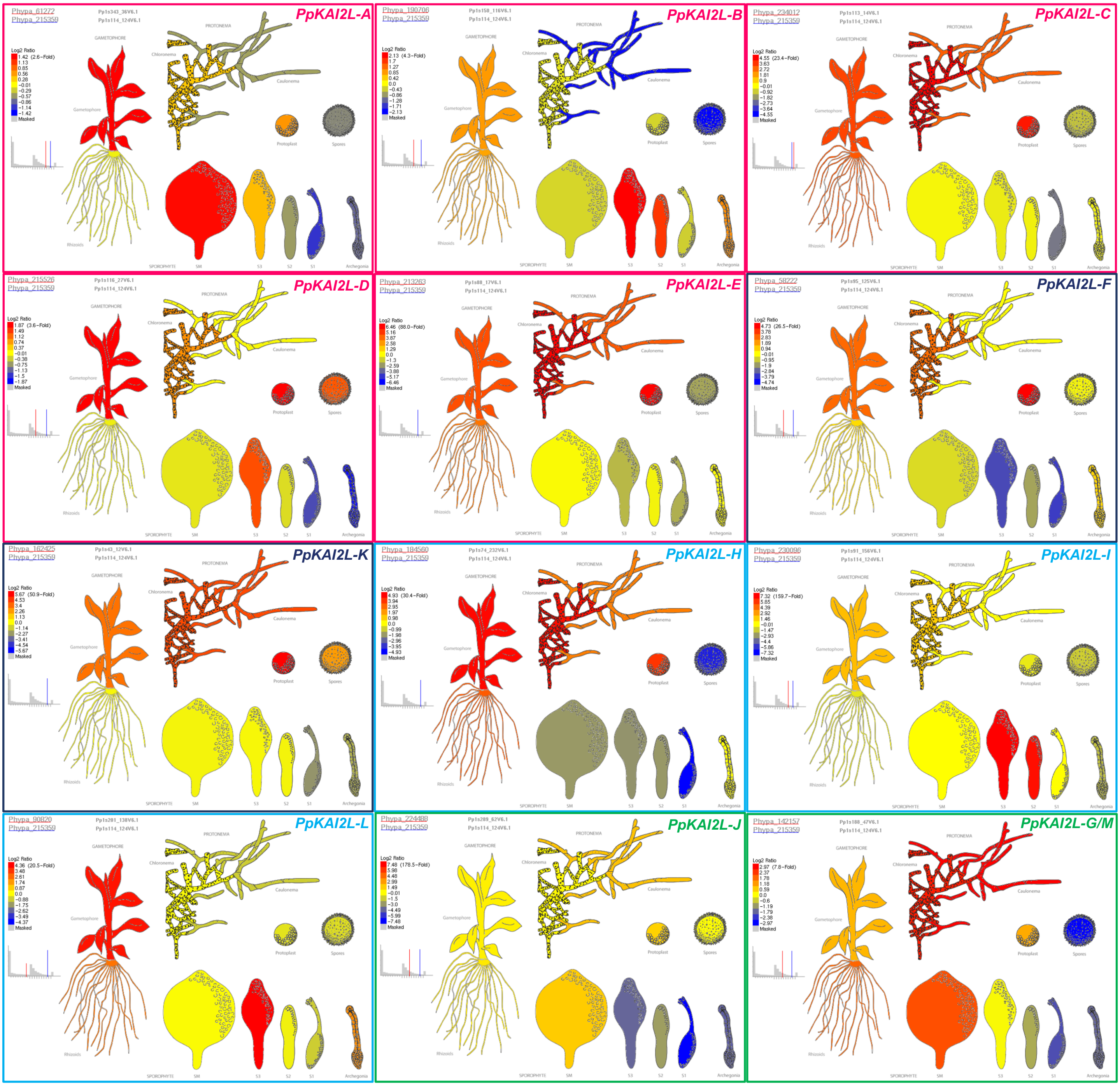


**Supplemental Figure 4: eFP-Browser expression data for *PpKAI2L* genes.**
Expression levels are shown in a color scale, relative to the expression of the *PpAPT* reference gene. Diagrams were taken from *www.bar.utoronto.ca* in May 2018. Owing to the extreme similarity of the *PpKAI2L-G* and *PpKAI2L-M* transcripts, they could not be singled out in the data set of Ortiz Ramirez et al. (2016).

Supplemental Figure 5

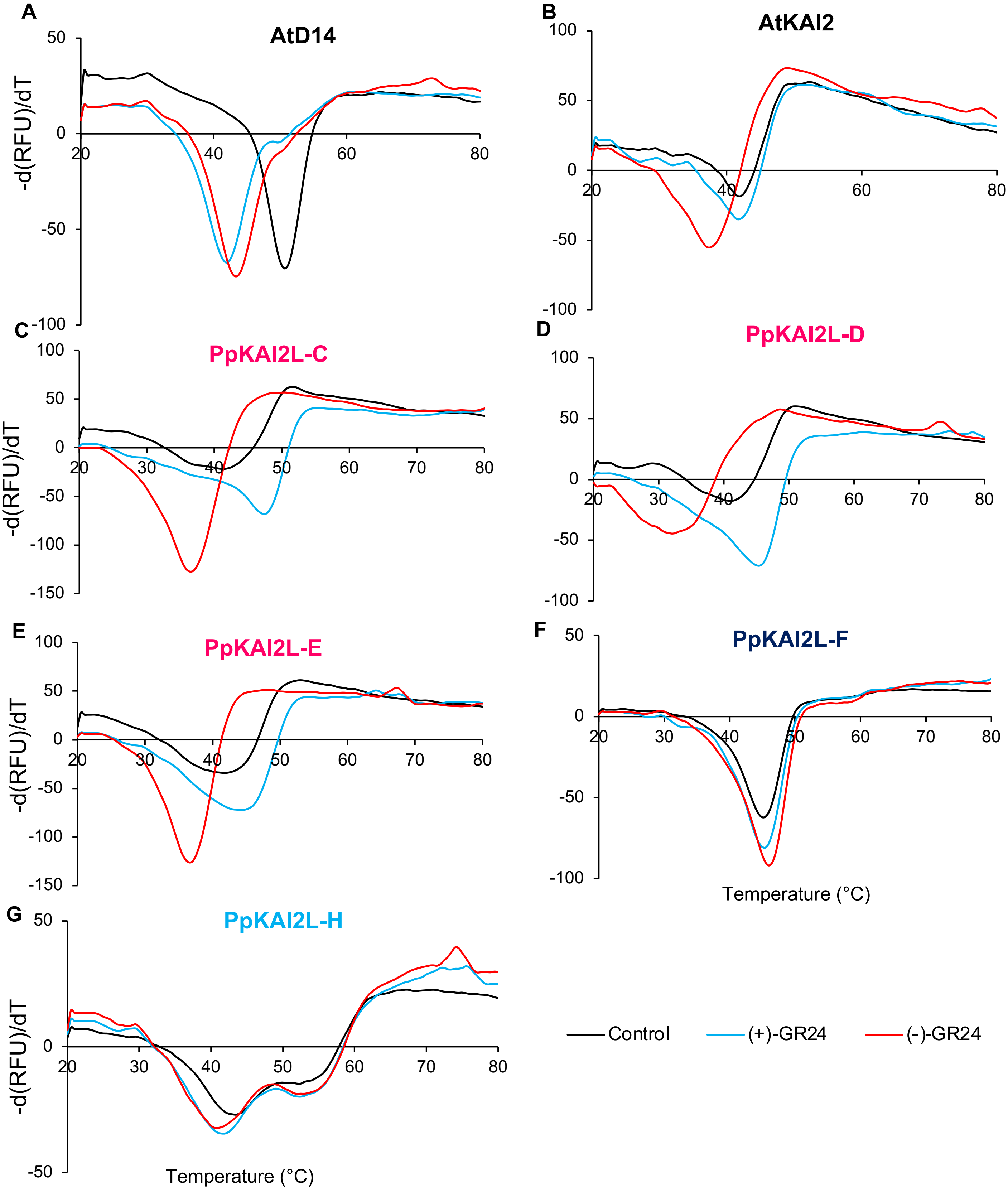

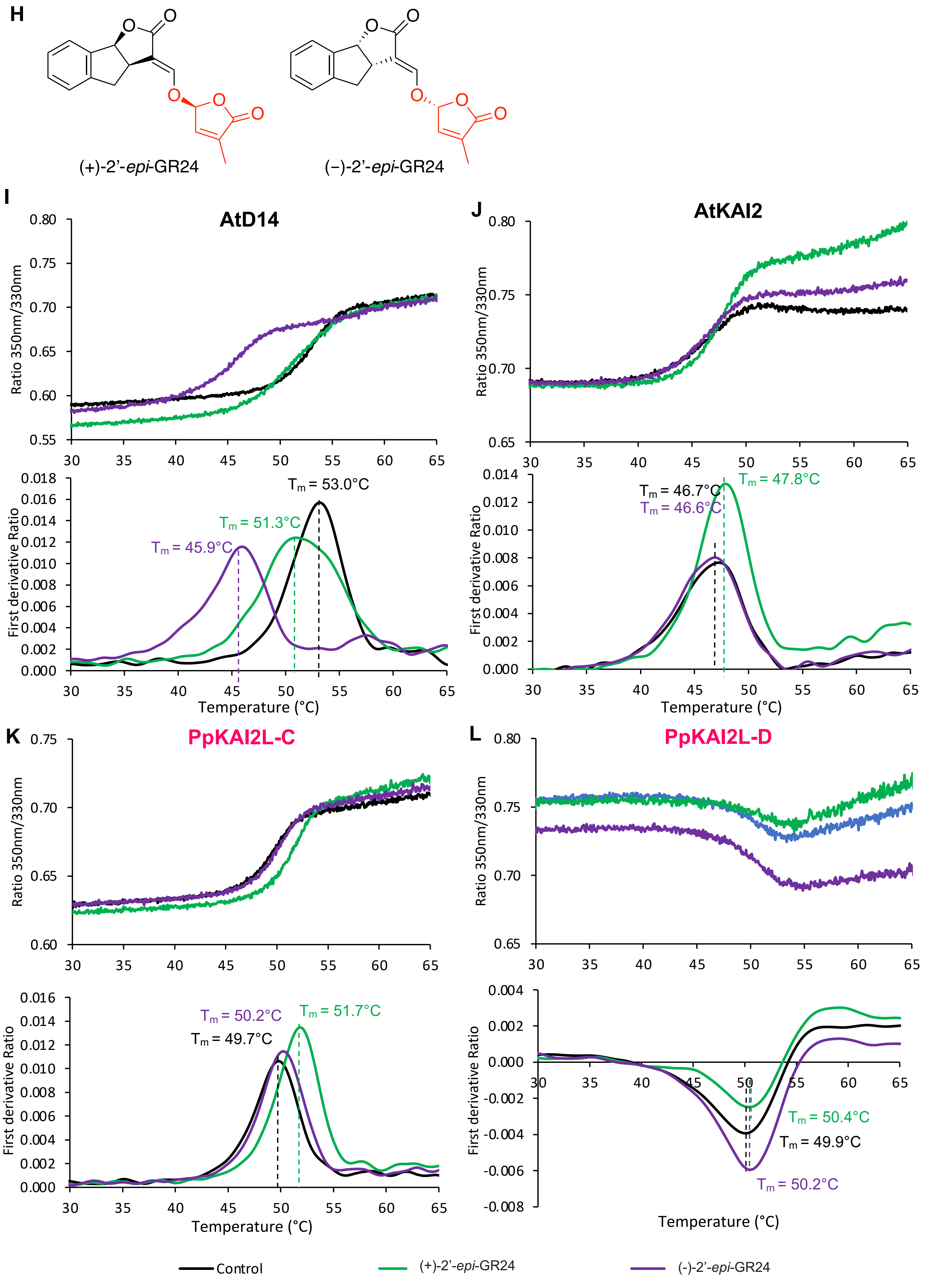
H
(+)-2'-*epi*-GR24
(−)-2'-*epi*-GR24
I
AtD14
Ratio 350nm/330nm
First derivative Ratio
$T_m$ = 53.0°C
$T_m$ = 51.3°C
$T_m$ = 45.9°C
Temperature (°C)
J
AtKAI2
Ratio 350nm/330nm
First derivative Ratio
$T_m$ = 47.8°C
$T_m$ = 46.7°C
$T_m$ = 46.6°C
Temperature (°C)
K
PpKAI2L-C
Ratio 350nm/330nm
First derivative Ratio
$T_m$ = 50.2°C
$T_m$ = 51.7°C
$T_m$ = 49.7°C
Temperature (°C)
L
PpKAI2L-D
Ratio 350nm/330nm
First derivative Ratio
$T_m$ = 50.4°C
$T_m$ = 49.9°C
$T_m$ = 50.2°C
Temperature (°C)
Control
(+)-2'-*epi*-GR24
(-)-2'-*epi*-GR24

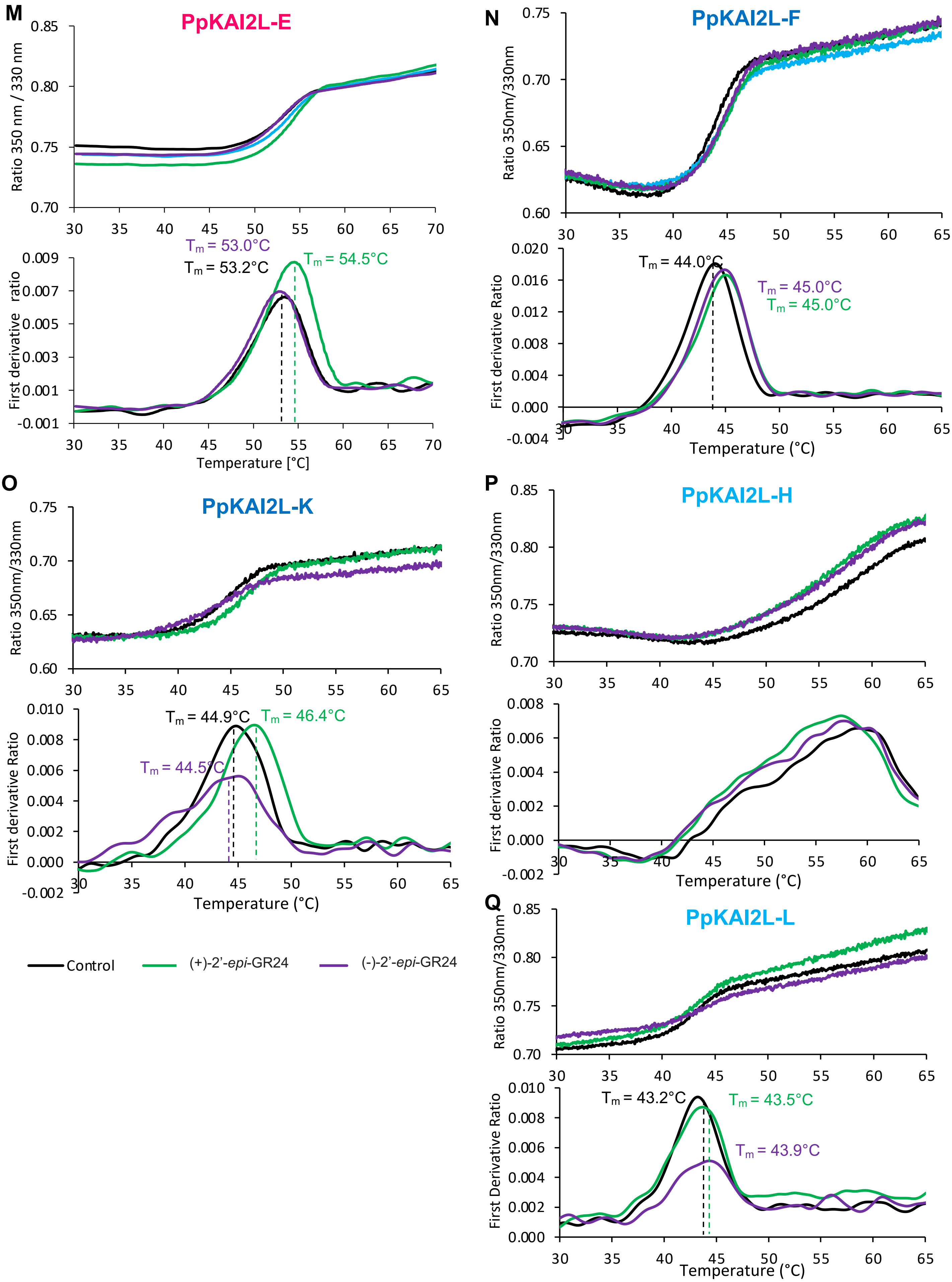
M
PpKAI2L-E
Ratio 350 nm / 330 nm
First derivative ratio
Tm = 53.0°C
Tm = 53.2°C
Tm = 54.5°C
Temperature [°C]
N
PpKAI2L-F
Ratio 350nm/330nm
First derivative Ratio
Tm = 44.0°C
Tm = 45.0°C
Tm = 45.0°C
Temperature (°C)
O
PpKAI2L-K
Ratio 350nm/330nm
First derivative Ratio
Tm = 44.9°C
Tm = 46.4°C
Tm = 44.5°C
Temperature (°C)
P
PpKAI2L-H
Ratio 350nm/330nm
First derivative Ratio
Temperature (°C)
Q
PpKAI2L-L
Ratio 350nm/330nm
First Derivative Ratio
Tm = 43.2°C
Tm = 43.5°C
Tm = 43.9°C
Temperature (°C)
Control
(+)-2'-epi-GR24
(-)-2'-epi-GR24

**Supplemental Figure 5: Response of the PpKAI2L proteins to the four GR24 isomers as determined by Differential Scanning Fluorimetry (DSF) (A-G) and nanoDSF (H-Q).**

Melting temperature curves of AtD14 (**A**), AtKAI2 (**B**), PpKAI2L-C (**C**), PpKAI2L-D (**D**), PpKAI2L-E (**E**), PpKAI2L-F (**F**), and PpKAI2L-H (**G**) at 10 µM, with (+)-GR24 (blue) and (−)-GR24 (red) at 200 µM, and without ligand (black) are shown as assessed by DSF. Each line represents the average protein melt curve for three technical replicates and the experiment was carried out twice. (**H**) Chemical structures of (+)-2'-*epi*-GR24 and (-)-2'-*epi*-GR24 isomers. (**I-Q**) Thermostability of AtD14, AtKAI2 and PpKAI2 proteins at 10 µM in the absence of a ligand (black line) or in the presence of (+)-2'-*epi*-GR24 (green line) or (-)-2'-*epi*-GR24 (purple line) at 100 µM, analyzed by nanoDSF. For each protein, top panel shows the changes in fluorescence (ratio $F_{350nm}/F_{330nm)}$ with temperature. The bottom panel shows the first derivative for the $F_{350nm}/F_{330nm}$ curve, against the temperature gradient from which the apparent melting temperatures ($T_m$) for each sample was determined. The experiment was carried out twice.

### Supplementary Figure 6

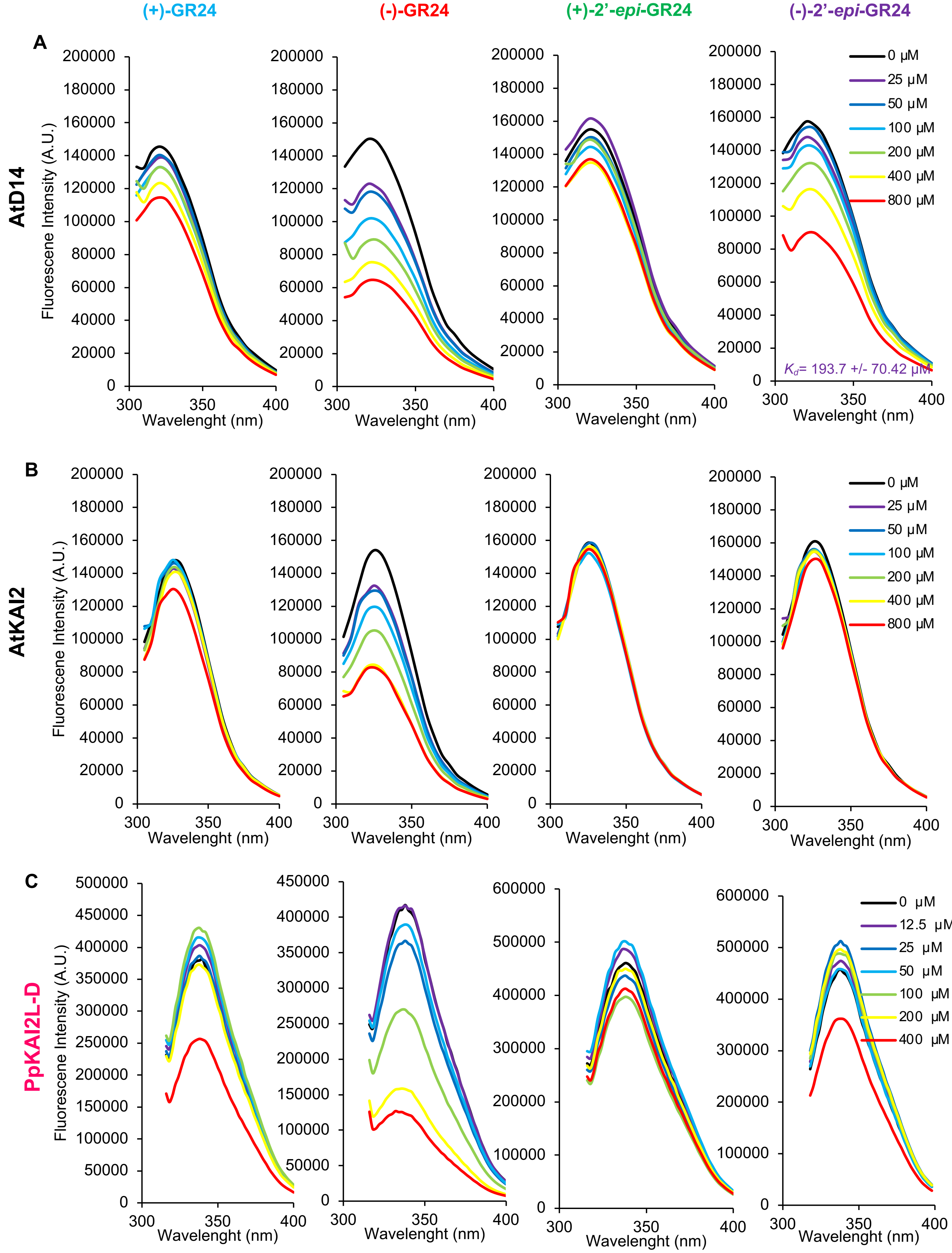

Supplemental Figure 6

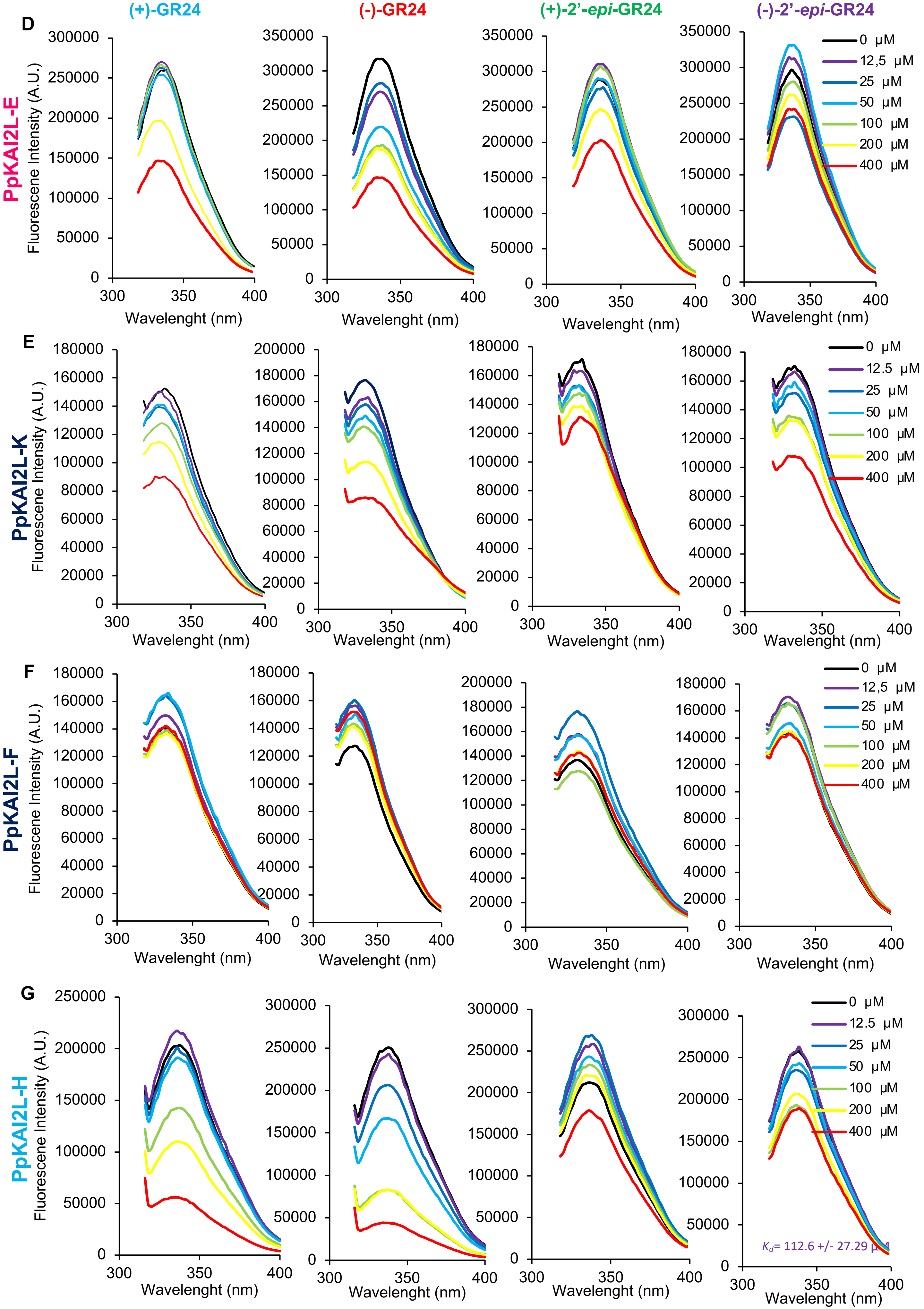

**Supplemental Figure 6: Intrinsic tryptophan fluorescence of AtD14 (A), AtKAI2 (B), PpKAI2L-D (C), PpKAI2L-E (D), PpKAI2L-K (E), PpKAI2L-F (F), and PpKAI2L-H (G) proteins in the presence of GR24 isomers.**

Changes in intrinsic fluorescence emission spectra of PpKAI2L proteins, in the presence of various concentrations of (+)-GR24 **(left)**, (−)-GR24 **(middle left)**, (+)-2'-*epi*-GR24 **(middle right)**, (−)-2'-*epi*-GR24 **(right)**. Proteins (10 µM) were incubated with increasing amounts of ligand (0–800 µM, top line to bottom line, respectively). The observed relative changes in intrinsic fluorescence were plotted as a function of SL analog concentration and transformed to degree of saturation and used to determine the apparent $K_d$ values relevant to Figure 5. Plots represent means of two replicates and the experiments were repeated at least three times.

### Supplemental Figure 7

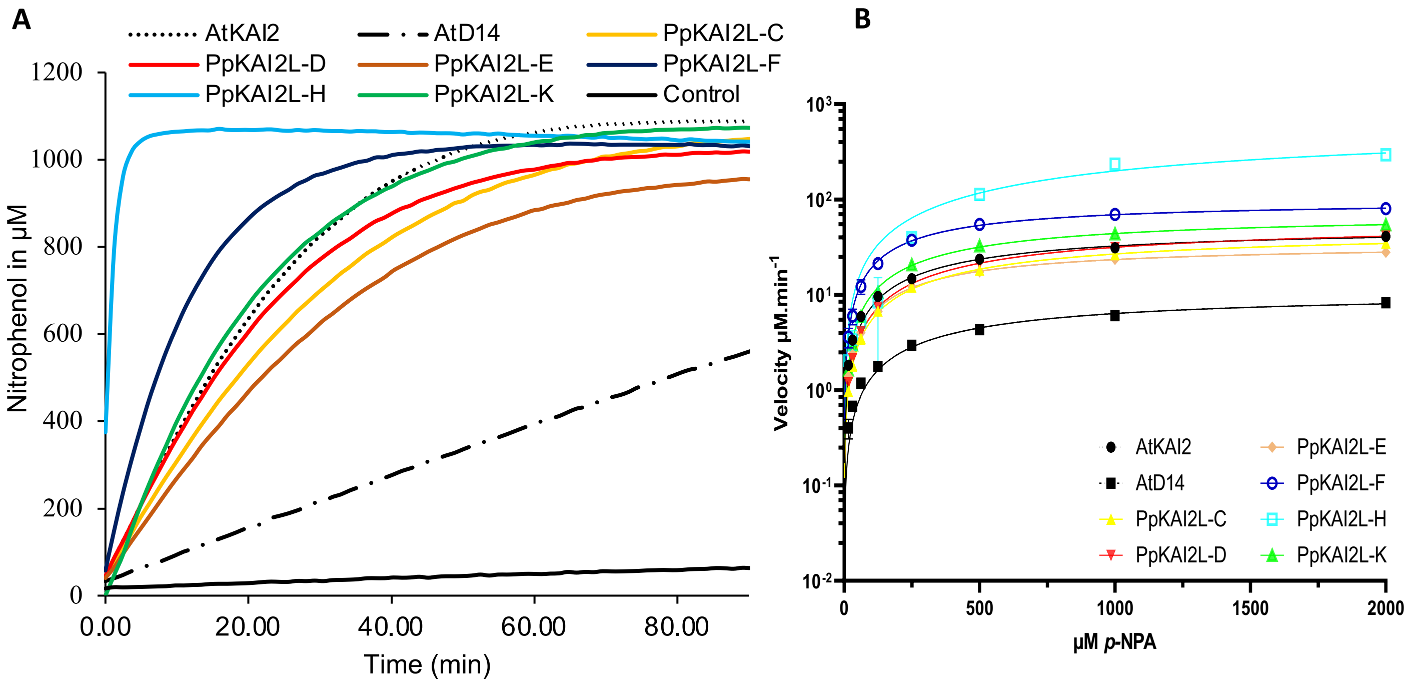


C

| | AtKAI2 | AtD14 | PpKAI2L-C | PpKAI2L-D | PpKAI2L-E | PpKAI2L-F | PpKAI2L-H | PpKAI2L-K |
|---|---|---|---|---|---|---|---|---|
| $V_{max}$ (µM $min^{-1}$) | 52.39 ±1.299 | 10.91 ±0.3434 | 48.48 ±1.088 | 59.54 ±1.152 | 34.75 ±0.3452 | 97.81 ±2.254 | 673.5 ±115.5 | 71.76 ±1.459 |
| $K_M$ *(µM)* | 610.1 ±36.69 | 714.3 ±51.91 | 807.4 ±40.24 | 888.7 ±37.02 | 483.5 ±12.5 | 413.9 ±25.98 | 2381 ±644 | 632.9 ±30.87 |
| $k_{cat}$ ($min^{-1}$) | 158.8 ±3.935 | 33.06 ±1.04 | 146.9 ±3.297 | 180.4 ±3.491 | 105.3 ±1.046 | 296.4 ±6.83 | 2041 ±350 | 217.4 ±4.421 |

**Supplemental Figure 7: PpKAI2L hydrolysis activity towards *p*-NPA.**

(**A**) Progress curves during 4-nitrophenyl acetate (*p*-NPA) (1 mM) hydrolysis by AtKAI2, AtD14 and PpKAI2L proteins (4 µM). The release of 4-nitrophenol was monitored ($A_{405}$) at 25 °C.

(**B**) Michaelis-Menten plot of AtKAI2, AtD14 and PpKAI2L steady state kinetics reaction velocity with *p*-NPA. Initial velocity was determined with *p*-NPA concentration from 15 µM to 2000 µM and protein at 4 µM. Points are the mean of 3 replicates and error bars represent SE. Experiments were repeated at least twice.

(**C**) Table: Kinetic constants of PpKAI2 proteins with the substrate *p*-NPA. $K_M$ and $k_{cat}$ are steady-state kinetic constants and values represent the mean ± SE of three replicates.

Supplemental Figure 8

A

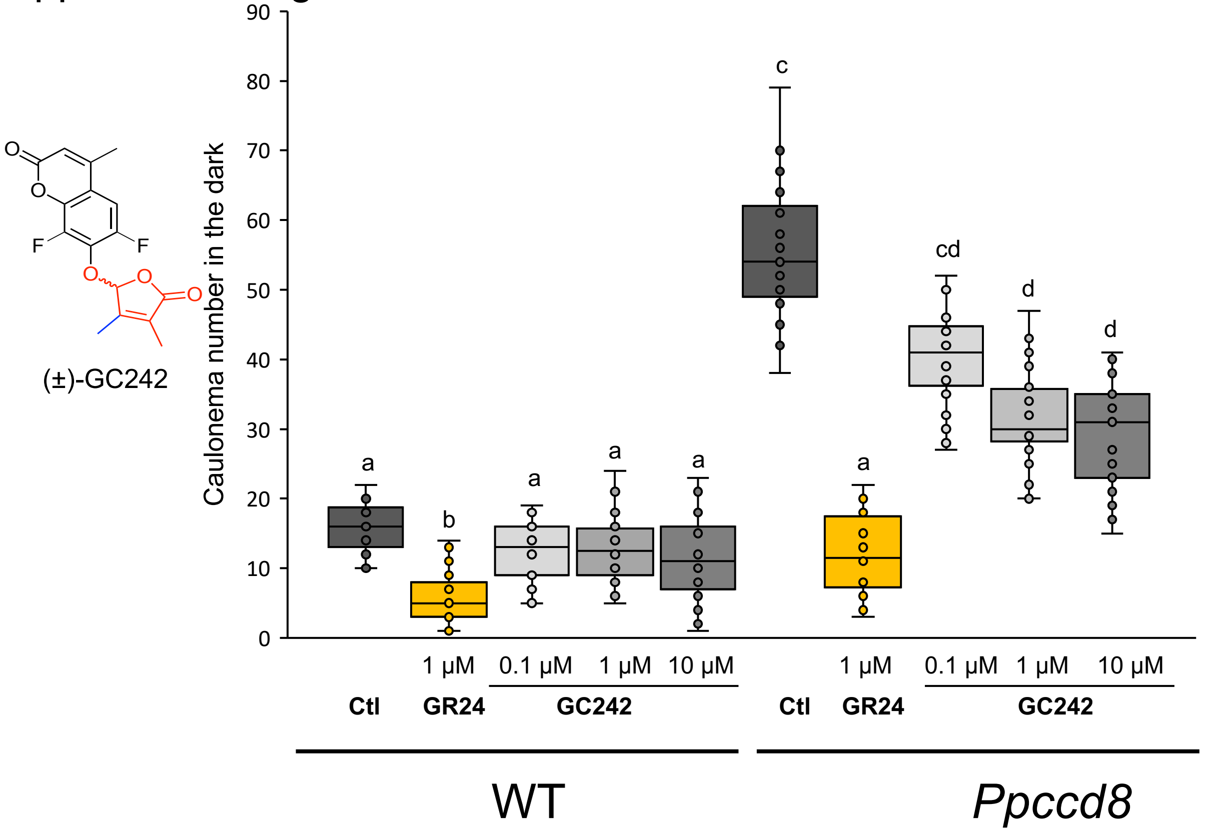


B

| | AtD14 | PpKAI2L-H | PpKAI2L-$H^{L28F}$ |
|---|---|---|---|
| $V_{max}$ (µM.min$^{-1}$) | 0.09566 ± 0.002728 | 0.06794 ± 0.004744 | 0.01465 ±0.001005 |
| $K_{1/2}$ (µM) | 0.7733 ± 0.09334 | 4.794 ± 1.056 | 4.675 ± 1.019 |
| $k_{cat}$ (min$^{-1}$) | 0.2899 ± 0.008268 | 0.2059 ± 0.01437 | 0.04438 ± 0.003047 |

**Supplemental Figure 8: Characterization of (±)-GC242 profluorescent probe activity on moss.**
(**A**) *Caulonema* number measurements in the dark in the WT and *Ppccd8* SL synthesis mutant, following application of 1 µM (±)-GR24 (yellow) and increasing concentrations of (±)-GC242. Ctl, control (same amount of DMSO). Mean of 3 biological repeats; n = 9-10 in each repeat. Statistical groups (comparison of all genotypes and treatments) are indicated by letters and were determined with a Kruskal-Wallis test followed by a Dunn *post hoc* test ($p < 0.05$).
(**B**) Kinetic constants of AtD14, PpKAI2L-H, PpKAI2L-$H^{L28F}$ towards (±)-GC242**.** $K_{1/2}$ and $k_{cat}$ are pre-steady-state kinetic constants. $K_{1/2}$ and $k_{cat}$ values represent the mean ± SE of three replicates.

Supplemental Figure 9

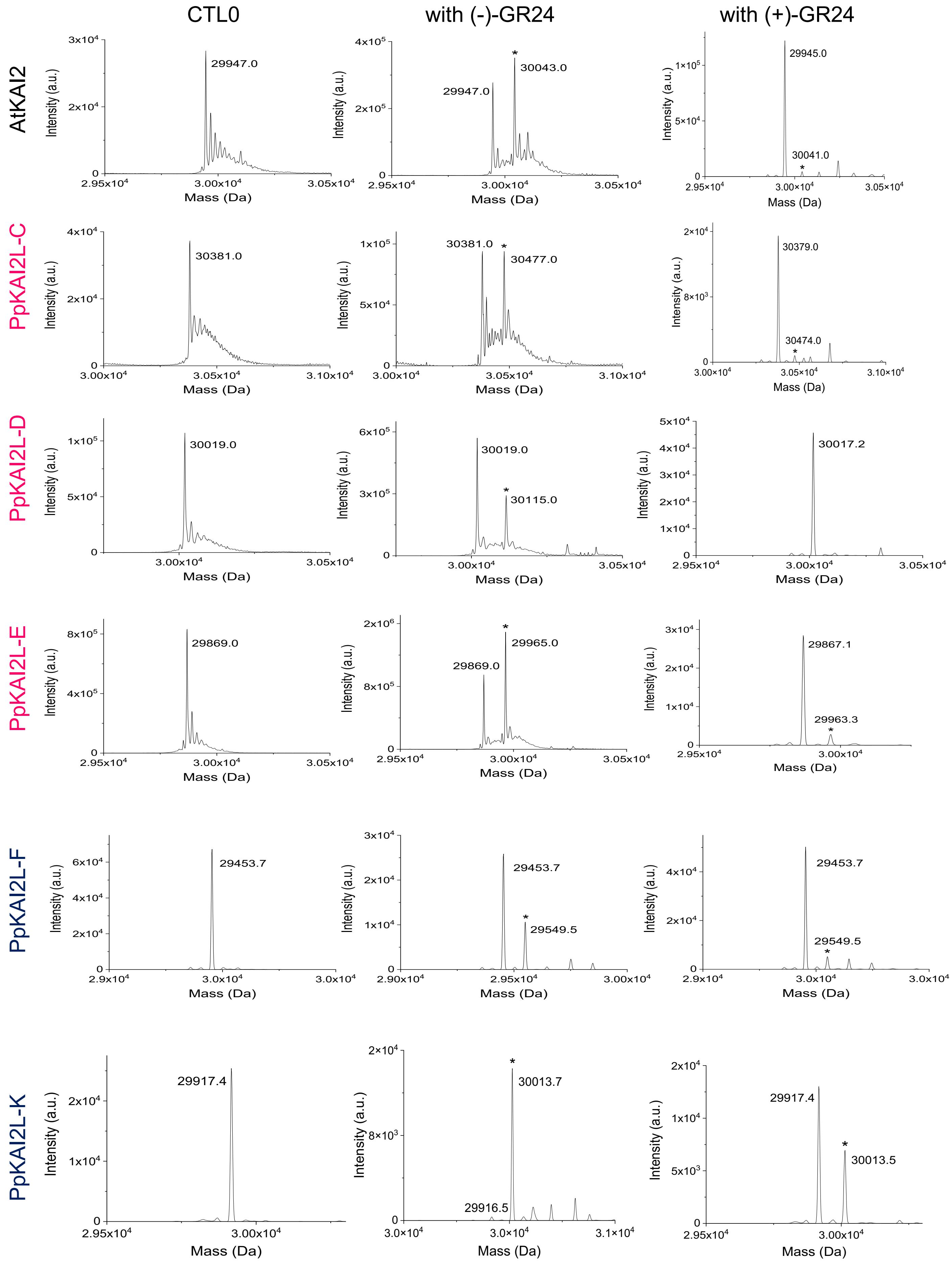

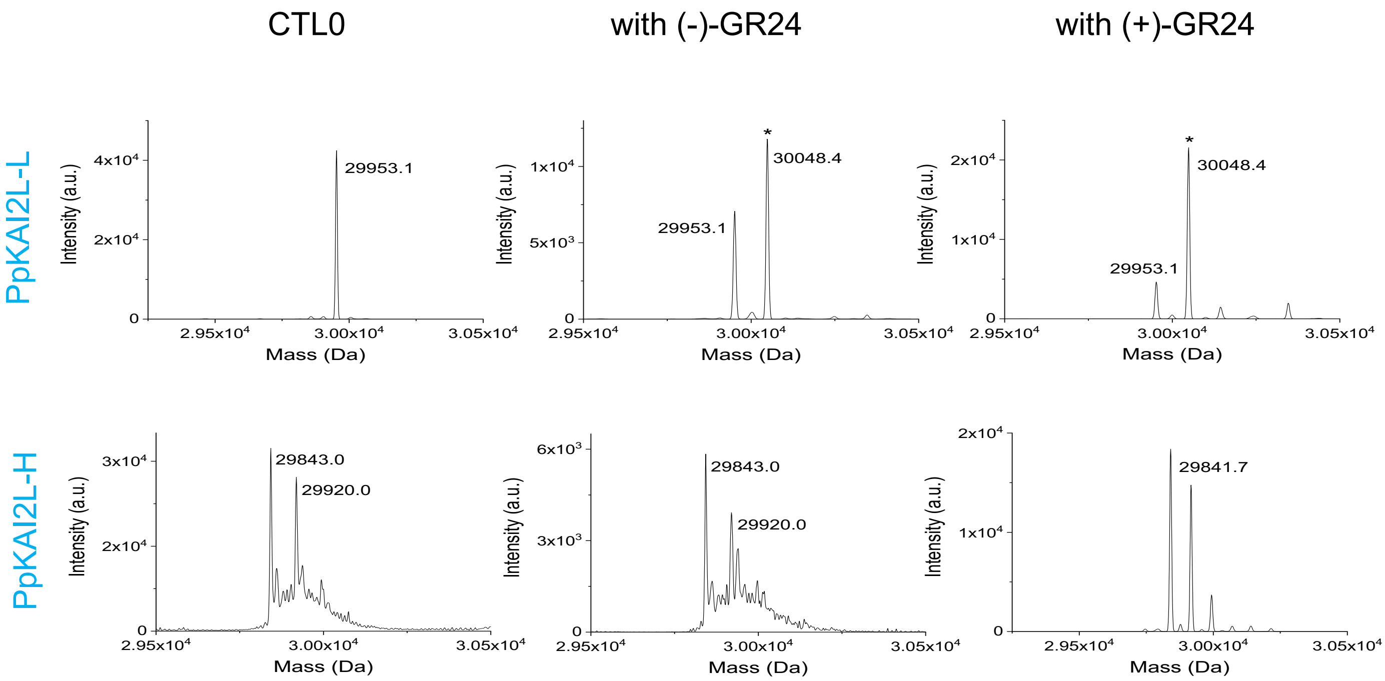


**Supplemental figure 9. Mass spectrometry characterization of covalent PpKAI2L-ligand complexes.** Deconvoluted electrospray mass spectra of PpKAI2L protein before (left column) and after adding (−)-GR24 (middle column) or (+)-GR24 (right column). Peaks with an asterisk correspond to PpKAI2L covalently bound to a ligand. Mass increments are measured for different PpKAI2L-ligand complexes: 96.3 Da for (+)-GR24 and (−)-GR24).

Supplemental Figure 10 A-B

A

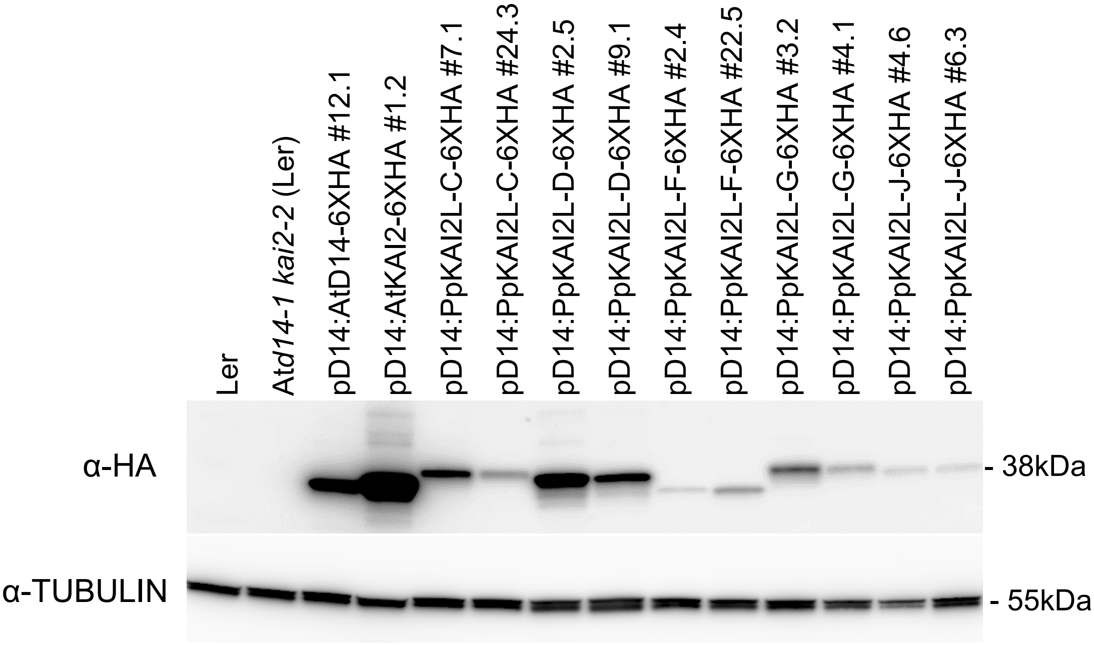


B

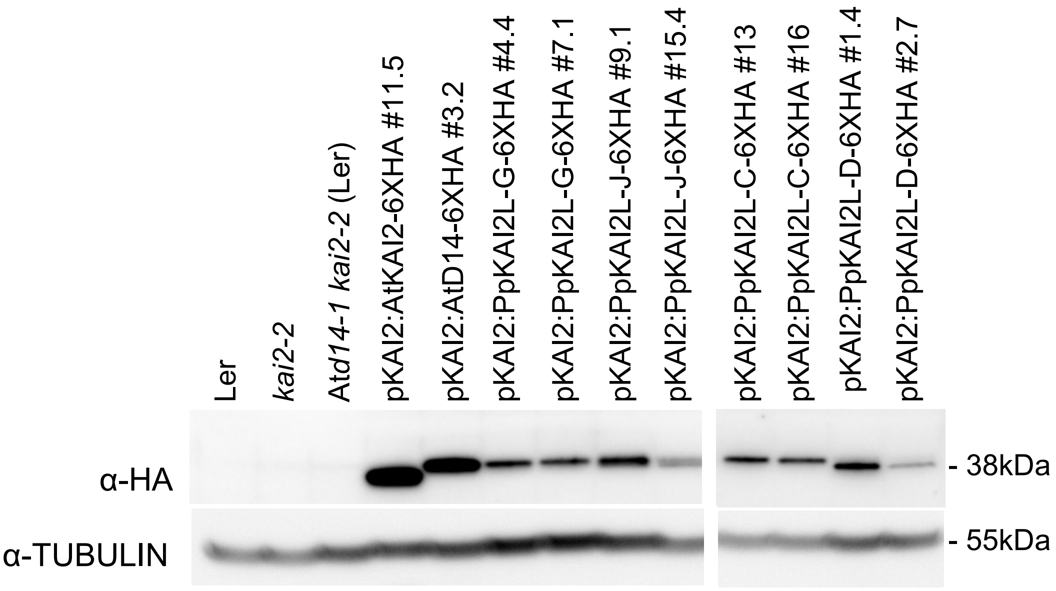

Supplemental Figure 10C:

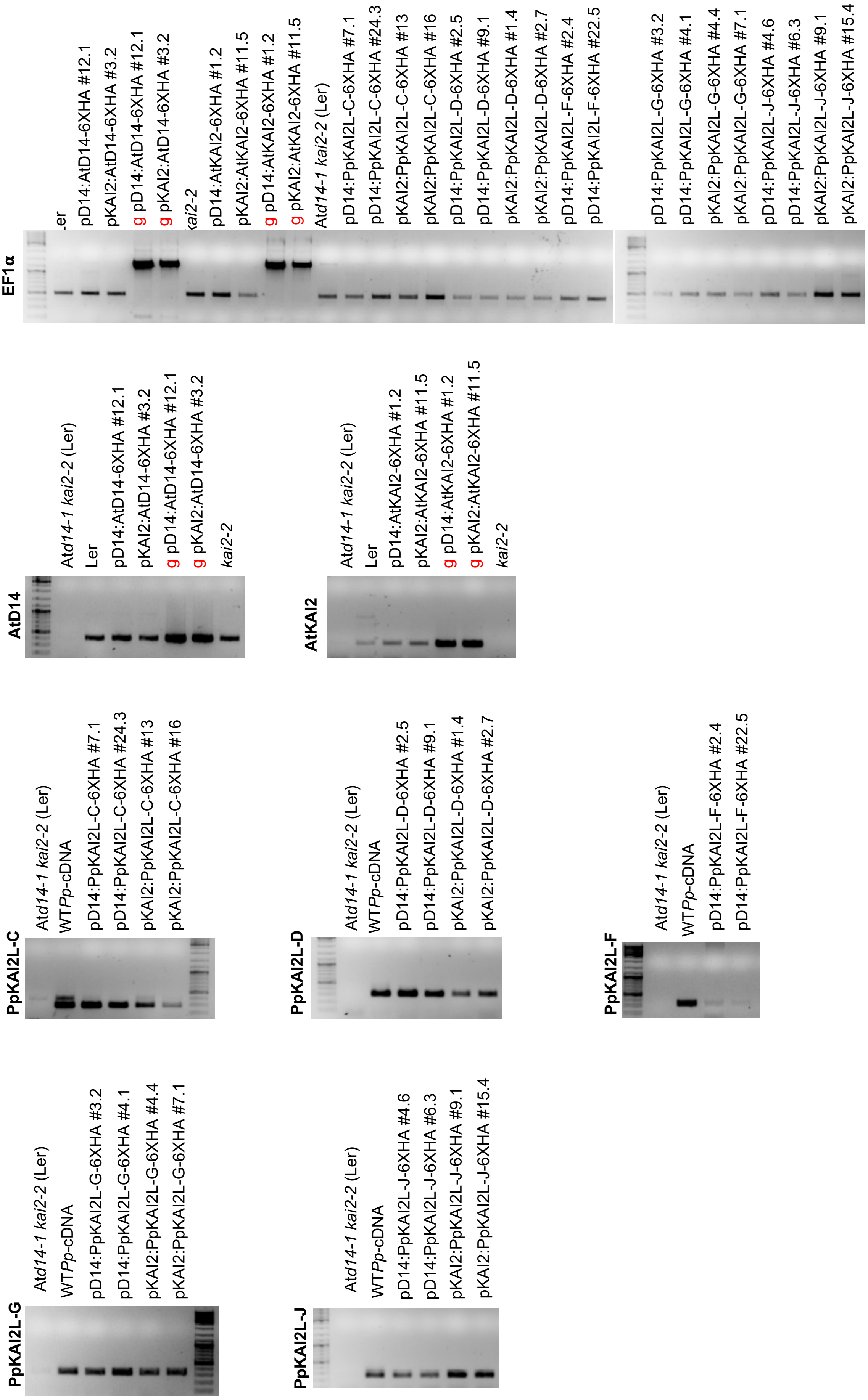

Supplemental Figure 10 D:

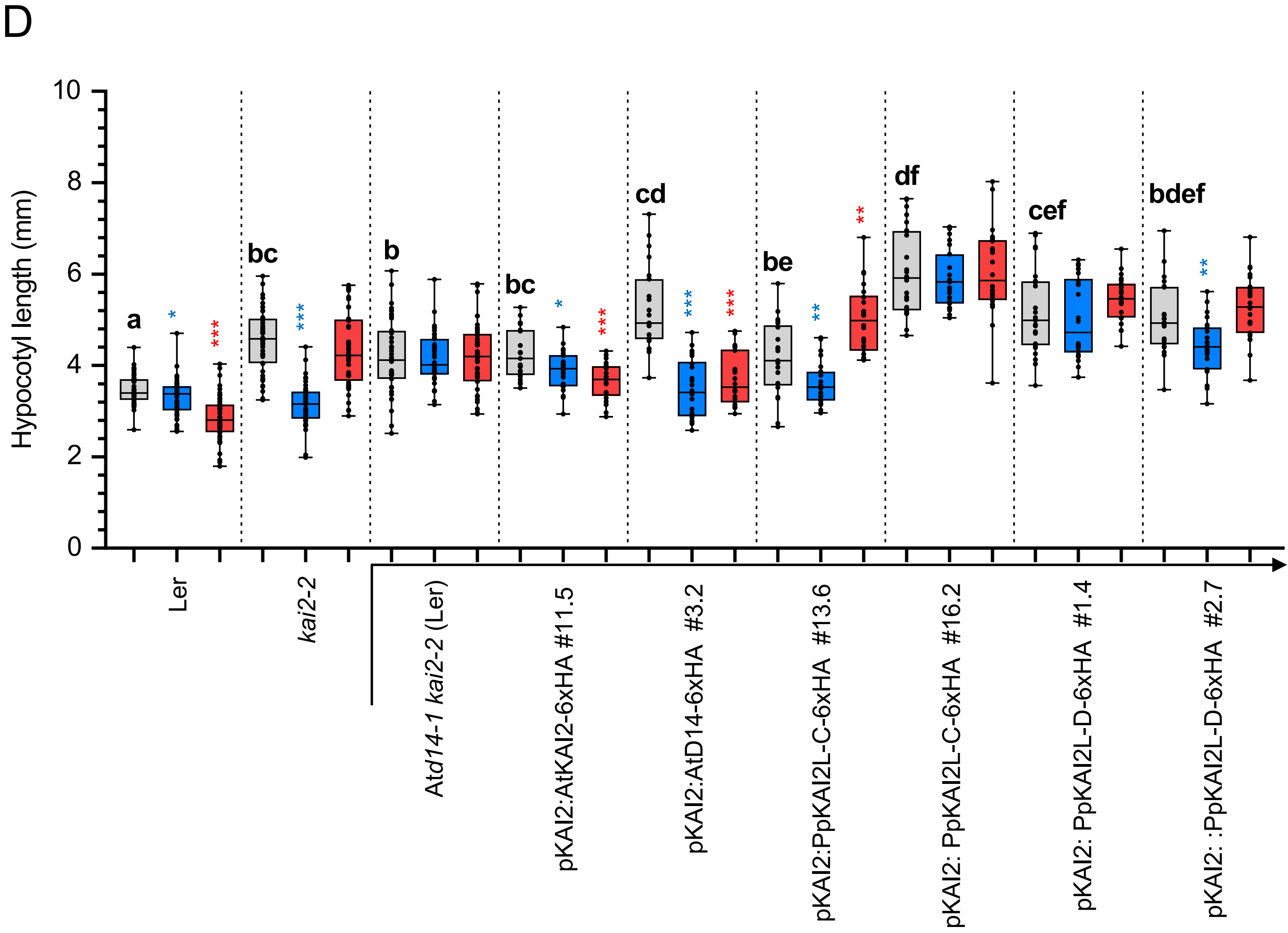


**Supplemental Figure 10: Complementation assays of the Arabidopsis At*d14-1 kai2-2* double mutant.**

**(A-B)** AtD14-6xHA, AtKAI2-6xHA and PpKAI2L-6xHA protein levels analyzed by immunoblot using an α-HA antibody in Ler (wild type), *kai2-2, Atd14-1 kai2-2 (Ler)* and transformed *Atd14-1 kai2-2* plants, expressed under the control of AtD14 promoter **(A)** or AtKAI2 promoter **(B).** Protein extracts from 10-d-old seedling were separated by 10% SDS-PAGE and identified as a 35 kD band. An immunoblot detecting tubulin was included for loading reference. **(C)** Results of PCR amplification of CDS from cDNAs of Arabidopsis transformed plants (homozygous T3 generation), as indicated. The presence of transcripts was verified in all samples using EF1**α** as a control. **(D)** Hypocotyl length of Ler (WT), *kai2-2*, At*d14-1 kai2-2* mutant (in Ler), and At*d14-1 kai2-2* mutant transformed using AtKAI2 promoter to control *AtD14*, *AtKAI2* (controls, same as shown figure 7B) or *PpKAI2L-C* and -*D* genes as noted below the graph. Hypocotyl length under low light, on ½ MS medium with DMSO (control, grey bars) 1 µM (+)-GR24 (blue bars) or 1 µM (–)-GR24 (red bars). Different letters indicate significantly different results between genotypes in control conditions based on a Kruskal–Wallis test ($p < 0.05$, Dunn *post hoc* test with p values corrected following the Benjamini-Hochberg method). Symbols in blue and red give the statistical significance of response to (+)-GR24 and (–)-GR24 respectively (Mann-Whitney tests, *** $p \leq 0.001$; ** $0.001<p<0.01$; * $0.01 \leq p < 0.05$).

### Supplemental Figure 11

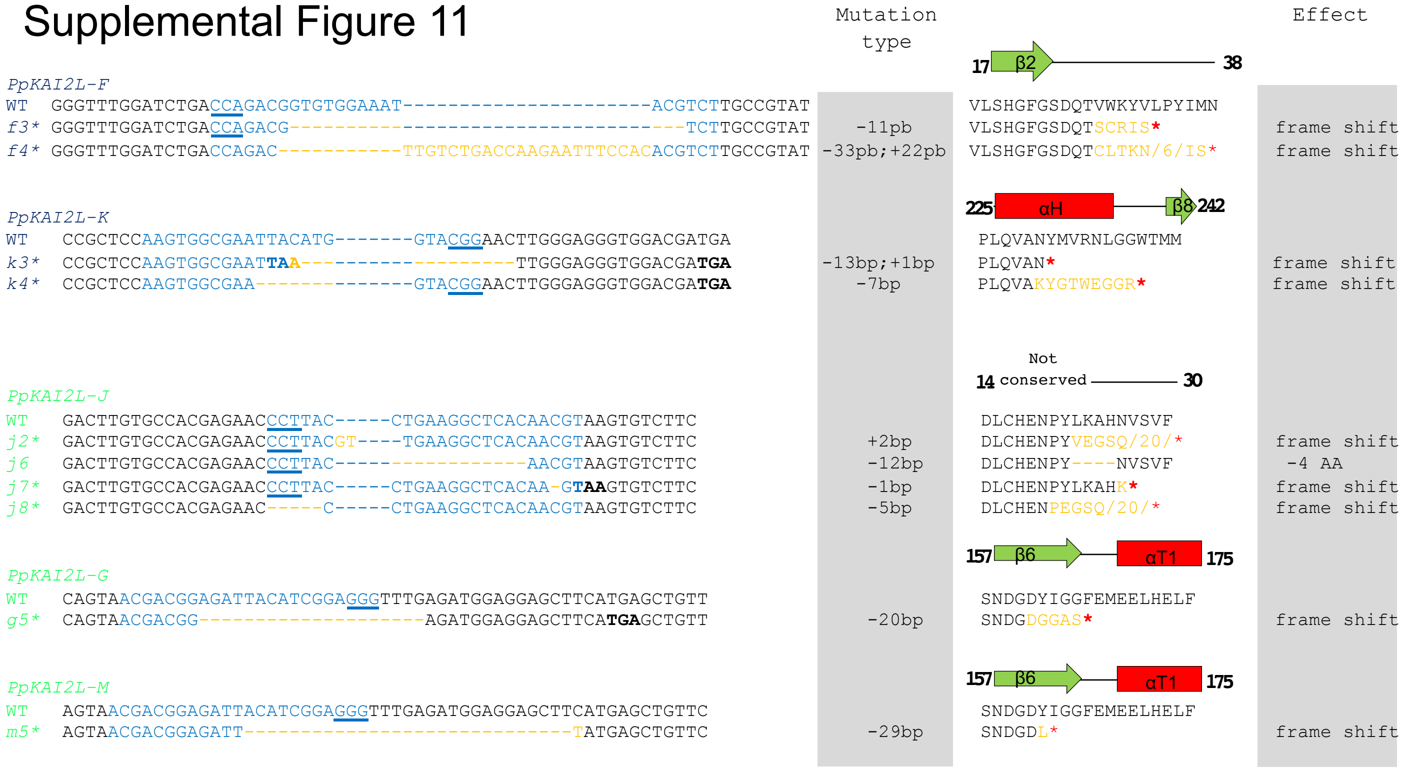


**Supplemental Figure 11: Extra mutations obtained in *PpKAI2L* genes.**
WT nucleotide and protein sequences are shown, above altered sequences found in various CRISPR-Cas9 lines (in italics, numbered). The number of first shown amino acid (aa) and the predicted secondary structure are indicated above the WT protein sequence. The crRNA sequence is shown in blue, with the PAM site underlined. Deletions are shown as dashes, insertions are noted with orange letters. The mutation type is shown on the right. Premature STOP codons are noted in bold, and with a red star on the aa sequence. On protein sequences, the number of masked aa is noted between slashes. See Supplemental Table 2 for the list of mutants carrying one or several of the shown mutations. Predicted knock out mutations due to premature STOP codons or a large deletion induced in the mutant sequences are noted with an *.

### Supplemental Figure 12

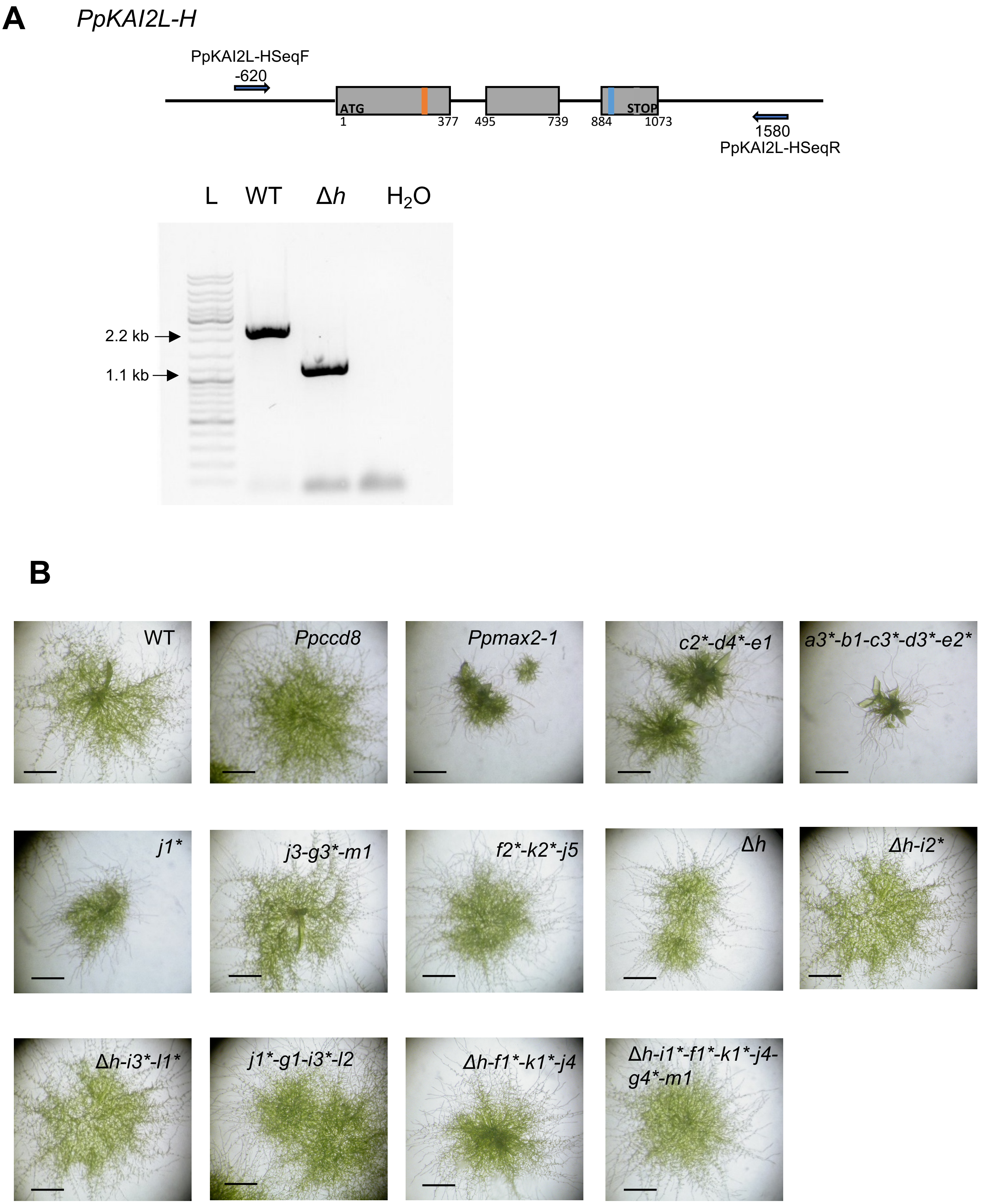


**Supplemental Figure 12: *Ppkai2L-Δh* mutant and phenotype of *Ppkai2L* mutants in light.**
**(A)** Effective deletion of the *PpKAI2L-H* gene after Homologous Recombination was verified. Primers used for PCR are indicated with blue arrows. PCR was carried out on WT and Δ*h* genomic DNA. Sequence of the deletion site is shown on Figure 8. **(B)** 10-day-old *Ppkai2L* mutants' phenotype in light conditions. Scale bars = 1 mm.

Supplemental Figure 13

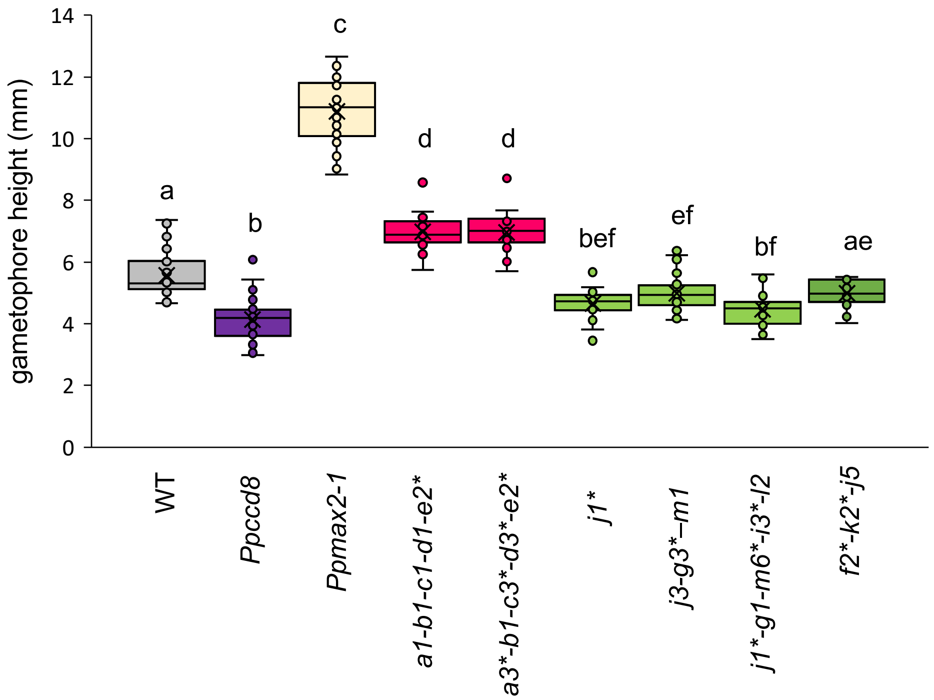


**Supplemental Figure 13: Gametophores of *Ppkai2L* mutants in red light.**
Gametophore height of *Ppkai2L* mutants, compared to that of WT, *Ppccd8* and *Ppmax2-1* mutants, following a 2-month growth under red light. Mutations are detailed in Figure 8 and Supplemental Table 2. Asterisks indicate null mutations. Box plots of n = 32-36 gametophores, grown in 3 Magenta pots, harboring between 15 and 25 leaves. Statistical groups (all genotypes comparison) are indicated by letters and were determined with a Kruskal-Wallis test followed by a Dunn *post hoc* test ($p < 0.05$).

#### Supplemental Figure 14

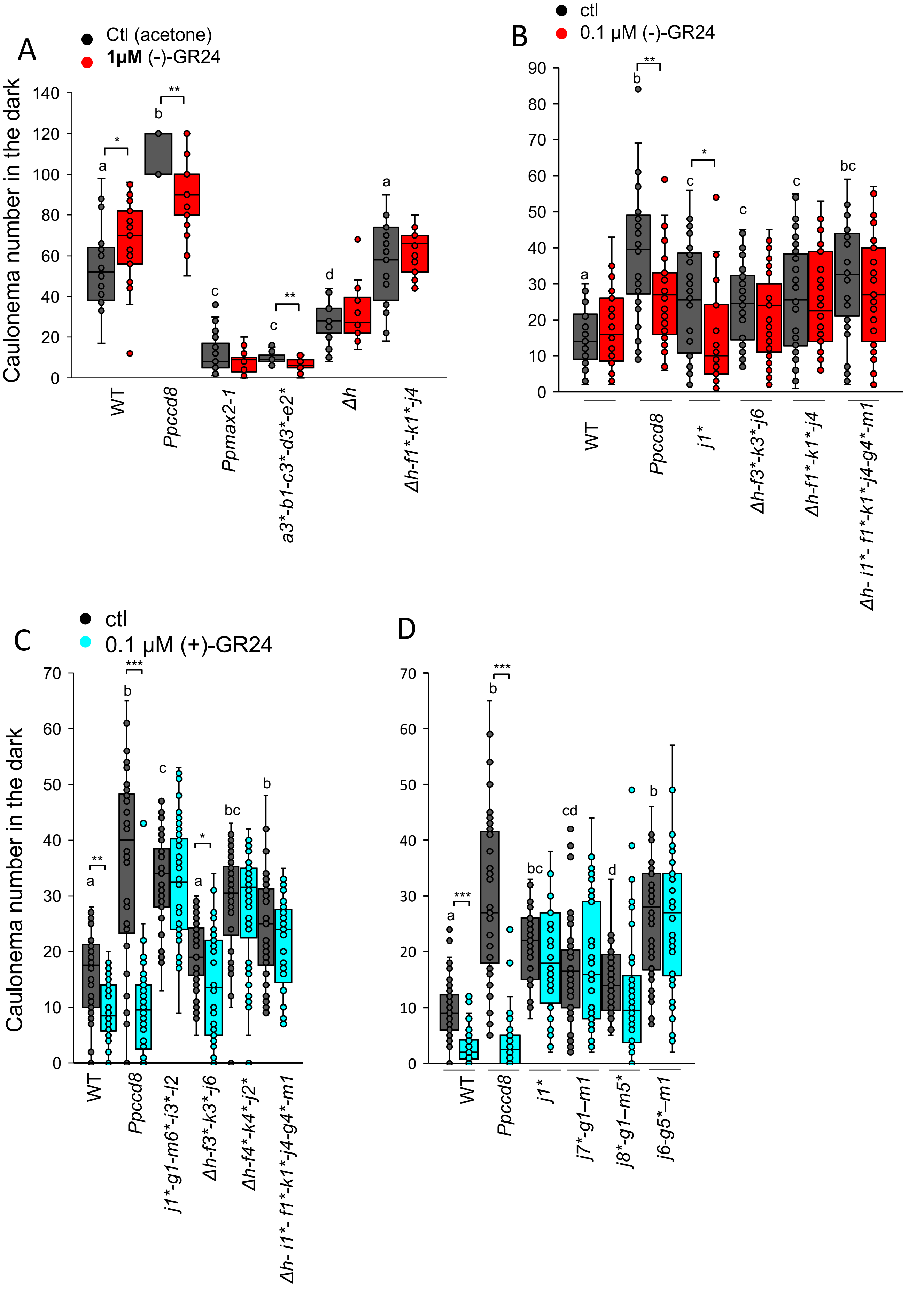

**Supplemental Figure 14: Phenotypic response of *Ppkai2L* mutants to (−)-GR24 and (+)-GR24 application: *caulonema* number in the dark.**

*Caulonema* numbers from *Ppkai2L* mutants following application of **(A)** 1 µM (−)-GR24 (in red), **(B)** 0.1 µM (+)-GR24 (in turquoise) **(C, D)** 0.1 µM (−)-GR24 (in red). DMSO was applied as control treatment (ctl, dark grey) except in (A), where acetone was applied. WT and both *Ppccd8* and *Ppmax2-1* mutants were used as control genotypes. Mutant genotypes carry mutations as indicated in Figure 8 and Supplemental Table 2, with asterisks for null mutations. For each genotype, *caulonema* were counted after two weeks in the dark, from 24 individuals, grown in three different 24-well plates. Statistical groups (comparing genotypes in control conditions) are indicated by letters and were determined by a one-way ANOVA with Welch test (95% Cl). Significant differences between control and treated plants within a genotype based on one-way ANOVA with Welch test (*** $p < 0.001$; ** $p < 0.01$; * $p < 0.5$).
