## Supplemental Tables for "The *Physcomitrium (Physcomitrella) patens* PpKAI2L receptors for strigolactones and related compounds highlight MAX2 dependent and independent pathways"

**Supplemental Table 1.** Oligonucleotides used in this study.

| **Primer name/purpose** | **Sequence (5'—3')** | note |
|---|---|---|
| Cloning for protein expression | | |
| PpKAI2L-C_attb1_HRV3C | ggggacaagtttgtacaaaaaagcaggctccctggaagtgctgtttcagggcccgATGGATGAGCTACCATCACTC | |
| PpKAI2L-C_attb2 | ggggaccactttgtacaagaaagctgggtctcaTCAACAAGACTCCAGATTCC | |
| PpKAI2L-D_attb1_HRV3C | ggggacaagtttgtacaaaaaagcaggctccctggaagtgctgtttcagggcccgATGGAGGAAGGACCAACACT | |
| PpKAI2L-D_attb2 | ggggaccactttgtacaagaaagctgggtctcaTTACAGACTTCCTGCGAGGT | |
| PpKAI2L-E_attb1_HRV3C | ggggacaagtttgtacaaaaaagcaggctccctggaagtgctgtttcagggcccgATGGAGGAGCCATCCTTGTT | |
| PpKAI2L-E_attb2 | ggggaccactttgtacaagaaagctgggtctcaTTATAGACTTCCAGCGAGGT | Gene specifics sequences are capitalized<br>Start and Stop codons are highlighted in bold<br>Gateway recombination sites are underlined<br>protease sites are in italic |
| PpKAI2L-F_attb1_HRV3C | ggggacaagtttgtacaaaaaagcaggctccctggaagtgctgtttcagggcccgATGCAGTCCCACAATGTGAT | |
| PpKAI2L-F_attb2 | ggggaccactttgtacaagaaagctgggtctcaTCATGATGCAAAACATCGCAA | |
| PpKAI2L-H_attb1_HRV3C | ggggacaagtttgtacaaaaaagcaggctccctggaagtgctgtttcagggcccgATGCCGAGCCCGTTGCTCTC | |
| PpKAI2L-H_attb2 | ggggaccactttgtacaagaaagctgggtctcaTCATGAGTCGATGCAGTGGAG | |
| PpKAI2L-K_attb1_HRV3C | ggggacaagtttgtacaaaaaagcaggctccctggaagtgctgtttcagggcccgATGATTCCGCAATCGAGCTC | |
| PpKAI2L-K_attb2 | ggggaccactttgtacaagaaagctgggtctcaTCACGGCGCAAGGCAGCGGAGA | |
| PpKAI2L-L-Δ47_attb1_HRV3C | ggggacaagtttgtacaaaaaagcaggctccctggaagtgctgtttcagggcccgATGGTGGTGTCCGAGTCCTT<br>G | |
| PpKAI2L-L_attb2 | ggggaccactttgtacaagaaagctgggtctcaTTAATCTTCAATGCAGCGAA | |

| Cloning for complementation assay | | |
|---|---|---|
| AtD14_promo_attB4 | AtD14 promoter | ggggacaactttgtatagaaaagttgccCCTCTTGTTGGATTCTTGGC |
| AtD14_promo_attB1R | AtD14 promoter | ggggactgcttttttgtacaaacttgcTTTTTTATGTGTTTGGGTTTGAGG |
| AtD14_attB1 | AtD14 coding sequence | ggggacaagtttgtacaaaaaagcaggcttcATGAGTCAACACAACATCTT |
| AtD14_attB2_ΔS | AtD14 coding sequence | ggggaccactttgtacaagaaagctgggtcCCGAGGAAGAGCTCGCCGGA |
| AtKAI2_promo_attB4 | AtKAI2 promoter | ggggacaactttgtatagaaaagttgccTTCACGACCAGTATGGTTTACTCA |
| AtKAI2_promo_attB1R | AtKAI2 promoter | ggggactgcttttttgtacaaacttgcCTCTCTAAAGAAGATTCTTCTCTGGTT |
| AtKAI2_attB1 | AtKAI2 coding sequence | ggggacaagtttgtacaaaaaagcaggcttcATGGGTGTGGTAGAAGAAGC |
| AtKAI2_attB2_ΔS | AtKAI2 coding sequence | ggggaccactttgtacaagaaagctgggtcCATAGCAATGTCATTACGAAT |

| PCR-Based mutagenesis | |
|---|---|
| PpKAI2H-L28F | GTGGTGCTGGGGCATGGCTTTGGAACCGACCAATCAG |

| Q-PCR primer | | | |
|---|---|---|---|
| PpKAI2L-A_qF468 | ATGGCAGTGCAGGAGTTTAG | PpKAI2L-A_qR571 | GTAACACGCTGCGCAAATC |
| PpKAI2L-B_qF631 | CTTGCCACATCTTGCAAAGC | PpKAI2L-B_qR735 | ATGCAGCACCTCAACAATGC |
| PpKAI2L-C_qF525 | GGAGTTTGGTAGGACGCTATTC | PpKAI2L-C_qR632 | GCACAGTCACCTTTGGTAGAA |
| PpKAI2L-D_qF623 | CACCGTCCCTTGCCATATT | PpKAI2L-D_qR736 | TCTGCAACACCTCGACAATC |
| PpKAI2L-E_qF667 | CCTTGGTAGTCGCGGATTAT | PpKAI2L-E_qR761 | GAACTGAGCTGAGGCAAATG |
| PpKAI2L-F_qF602 | TGAAAGTGCCAGTGCATCTC | PpKAI2L-F_qR744 | TCACTCAAATGCGGCAAG |
| PpKAI2L-K_qF589 | TCTTCCAGAGCGATCTACGTTC | PpKAI2L-K_qR738 | GTTCAGAACCTCCATCATCGTC |
| PpKAI2L-H_qF671 | AATTGAAGTGGCGGAGTACC | PpKAI2L-H_qR779 | ACCACCAATTCTGGACAACTC |
| PpKAI2L-I_qF509 | TGACGACAAAGCAGTGCAAG | PpKAI2L-I_qR643 | TATGGCAAGGCACTGTAACCTC |
| PpKAI2L-L_qF453 | AGGTTGCCTTGCATCTCTTG | PpKAI2L-L_qR569 | AGGTCATGCTGCTCAAATCC |
| PpKAI2L-J_qF393 | CTCATTCTCATGGCAGCATCTC | PpKAI2L-J_qR538 | CCATCGCCTTAGGTACAAAACC |
| PpKAI2L-G/M_qF638 | GCGACCAGATATTGCCCTTA | PpKAI2L-G/M_qR743 | ACTCCACTTTGCACTAGATAGC |
| PpKUF1LA_qF | GGAGGTGCTCATTGGAACTAAA | PpKUF1LA_qR | GGTGCATCCGAAGCAATATCTA |
| Pp3c6_15020_qF | CAGAACGGCTTTGTGGATTTG | Pp3c6_15020_qR | GTCCGAGTTGGTAGAGGTAGTA |
| PpAPT_qF | ACTTGCCGTGGCGAGCTAC | PpAPT_qR | CATCCTTGGAGGCCGACATC |
| PpACT3_qF | AGCGAGTACGATGAATCTGG | PpACT3_qR | ACACAGCAAGAGCTCAATCC |

| SemiQ-RT PCR | | | |
|---|---|---|---|
| AtEF1-F | TAATGCCAGTCTTGAGGCTG | AtEF1-R | TTGTTCAGTCTTTGTGGCCG |
| AtD14.F4 | AGAAGCTCTAAATGTCCGGG | Atd141RP2 | CTCACCTTCTTCGAATCCTCC |
| kai2.2.lp | CGTCCTCTACGACAACATGG | Kai2-2_RP-3.R | TTCTTGTTCGAATCCACCTTG |
| PpKai2.CF1 | TGCAGTCCAATTTCAAGGCC | PpKai2.CR1 | GCAGCACCTCAACTATCGTG |
| PpKai2L.DR1 | ACCCCGAGTACTTCAGCTTC | PpKai2L.DR1 | AATCCTGAGACCCATGCCTC |
| PpKai2.L.FF2 | GCACCTCCGCAGATGATTTCA | PpKai2.L.FR2 | GCCACCAACAACTCTCCAAA |
| PpKai2.L.GF1 | ATCCTCGATGAGTTGGGCAT | PpKai2.L.GR1 | AGGGCAATATCTGGTCGCAT |
| PpKai2.JF1 | TATGCTGACGACTTGCTTGC | PpKai2.JR1 | GTACAAAACCAGTGGCCCAG |

**Supplemental Table 1 (following).** Oligonucleotides used in this study.

| Primer name/purpose | Sequence (5'—3') | note |
|---|---|---|
| sgRNA | | |
| sgRNA_A_66F | GTTGTCCTCGGCCATGGTTTTGG | |
| sgRNA_B_2F | GTGTCCGGGACCGACTCTGTTGG | |
| sgRNA_C_175R | GCTAAAATACTCCGGGTCGGTGG | |
| sgRNA_D_85F | TCGGCACGGATCAATCTGTGTGG | |
| sgRNA_E_281F | TGGACATTCGGTGGCCGGTATGG | |
| sgRNA_F_98R | AGACGTATTTCCACACCGTCTGG | |
| sgRNA_K_1077F | AAGTGGCGAATTACATGGTACGG | |
| sgRNA_I_172R | GTGCTCAGGGTCCGTAGTGCCGG | |
| sgRNA_L_736R | GCCCAGAACACTCAGATTGTGGG | |
| sgRNA_J_57R | ACGTTGTGAGCCTTCAGGTAAGG | |
| sgRNA_GM_560F | ACGACGGAGATTACATCGGAGGG | Similar for G and M |
| mutant sequencing primers | | |
| PpKAI2L-A SeqF | GGAGATAGAACCATGTCTTACTCTCCTACTATTATTGAGG | |
| PpKAI2L-A SeqR | TCCACCTCCGGATCAAACGCATACATCGCAAACAA | |
| PpKAI2L-B SeqF | GTAACATTGCGCACCTTGTG | |
| PpKAI2L-B SeqR | ATGCAGCACCTCAACAATGC | |
| PpKAI2L-C SeqF | GGAAACACGTCATTCCTCATC | |
| PpKAI2L-C SeqR | TCTAGGAGAAGCGGAGATAGTG | |
| PpKAI2L-D SeqF | GGAGATAGAACCATGGAGGAAGGACCAACACTCT | |
| PpKAI2L-D SeqR | TCGCCACACTAAAGGCTATGT | |
| PpKAI2L-E SeqF | CTCGAGATACTCTACCCTTCACG | |
| PpKAI2L-E SeqR | GAACTGAGCTGAGGCAAATG | |
| PpKAI2L-F SeqF | GGAGATAGAACCATGCAGTCCCACAATGTGATAAT | |
| PpKAI2L-F SeqR | TCACTCAAATGCGGCAAG | |
| PpKAI2L-K SeqF | ATCTCGCGGTTTTCTGTGAT | |
| PpKAI2L-K SeqR | GCAGTGCGAACTGAGTGATG | |
| PpKAI2L-H SeqF | CTTTAAATTCACGCTTCAATGG | |
| PpKAI2L-H SeqR | GAGATAATACACGCGTTTTCATGGAC | |
| PpKAI2L-I SeqF | GGAGATAGAACCATGGTGATTTCCAAGGCGTTTCT | |
| PpKAI2L-I SeqR | TATGGCAAGGCACTGTAACCTC | |
| PpKAI2L-L SeqF | CTACCATGGAGGATTTGAGCAGC | |
| PpKAI2L-L SeqR | TGCAGCGAAGAAGTACTGGA | |
| PpKAI2L-J SeqF | GGAGATAGAACCATGATTCAGAAATTGACACCACCTGAGAC | |
| PpKAI2L-J SeqR | CCATCGCCTTAGGTACAAAACC | |
| PpKAI2L-G SeqF | AGCTTCACCTTCTCTCGTTACA | |
| PpKAI2L-G SeqR | CACTAGATAGCATGGAATCATTACC | |
| PpKAI2L-M SeqF | AGCTTCACCTTCTCTCGTTACA | |
| PpKAI2L-M SeqR | TAGATAGCATGGAATCGTCACT | |

### Supplemental Table 2 : Mutants used in the study

| **Mutant name** | **Clade** | **Mutation effects** |
|---|---|---|
| *Ppccd8* | na | PpCCD8: deletion Proust et al 2011 |
| *Ppmax2-1* | na | PpMAX2: Full CDS deletion, Lopez-Obando et al 2018 |
| *a2*-b4*-c2** | A-E | STOP in PpKAI2L-A (41), -B (15) and –C (64) |
| *c2*-d4*-e1* | A-E | STOP in PpKAI2L-C (64) and –D (39)<br>PpKAI2L-E: -($A^{99}G^{100}$) + ($D^{99}I^{100}R^{101}$) |
| *a1-b1-c1-d1-e2** | A-E | PpKAI2L-A: -($F^{30}G^{31}$); PpKAI2L-B: -($L^{7}LEA^{10}$); PpKAI2L-C: -($M^{55}GAGTTD^{61}$); PpKAI2L-D: -($Q^{33}S^{34}$) +($R^{33}$); STOP in PpKAI2L-E (106) |
| *a3*-b1-c3*-d3*-e2** | A-E | STOP in PpKAI2L-A (41), -C (64), -D (41) and -E (106); PpKAI2L-B: -($L^{7}LEA^{10}$) |
| *j1** | J,G,M | STOP in PpKAI2L-J (48) |
| *j1*-g2*–m2** | J,G,M | STOP in PpKAI2L-J (48), -G (174) and –M (182) |
| *j3-g3*–m1* | J,G,M | PpKAI2L-J: -($E^{18}NPY^{21}$); PpKAI2L-G -($G^{164}$--$W^{185}$); PpKAI2L-M: -($G^{160}DYI^{164}$) |
| *j7*-g1–m1* | J,G,M | STOP in PpKAI2L-J (27); PpKAI2L-G: -($G^{160}DYI^{164}$); PpKAI2L-M: -($G^{160}DYI^{164}$) |
| *j6-g5**–m1 | J,G,M | PpKAI2L-J: -($L^{22}KAH^{25}$); STOP in PpKAI2L-G (166); PpKAI2L-M: -($G^{160}DYI^{164}$) |
| *j8*-g1–m5** | J,G,M | STOP in PpKAI2L-J (44); PpKAI2L-G: -($G^{160}DYI^{164}$); STOP in PpKAI2L-M (163) |
| *f2*-k2*-j5* | F,K ; J,G,M | STOP in PpKAI2L-F (31); STOP in PpKAI2L-K (241); PpKAI2L-J: -($L^{22}KAHN^{26}$)+($Y^{22}$) |
| *Δh* | H,I,L | PpKAI2L-H: Full CDS deletion |
| *Δh-i2** | H,I,L | PpKAI2L-H: Full CDS deletion; STOP in PpKAI2L-I (62) |
| *Δh-i3*-l1** | H,I,L | PpKAI2L-H: Full CDS deletion; PpKAI2L-I: +($M^{59}VLSA^{63}$); STOP in PpKAI2L-L (74) |
| *j1*-g1-m6*-i3*-l2* | J,G,M ; H,I,L | STOP in PpKAI2L-J (48); PpKAI2L-G: -($G^{160}DYI^{164}$);STOP in PpKAI2L-M: 181 PpKAI2L-I: +($M^{59}VLSA^{63}$); PpKAI2L-L: -$N^{59}$ |
| *Δh-f1*-k1*-j4* | F,K ; J,G,M ; H,I,L | PpKAI2L-H: Full CDS deletion; STOP in PpKAI2L-F (31); STOP in PpKAI2L-K (233); PpKAI2L-J: -($L^{22}KAH^{25}$) |
| *Δh-f3*-k3*-j6* | F,K ; J,G,M ; H,I,L | PpKAI2L-H: Full CDS deletion; STOP in PpKAI2L-F (33); STOP in PpKAI2L-K (231); PpKAI2L-J: -($L^{22}KAH^{25}$) |
| *Δh-f4*-k4* -j2** | F,K ; J,G,M ; H,I,L | PpKAI2L-H: Full CDS deletion; STOP in PpKAI2L-F (41); STOP in PpKAI2L-K (239); STOP in PpKAI2L-J (47) |
| *Δh-i1*-f1*-k1*-j4-g4*-m1* | F,K ; J,G,M ; H,I,L | PpKAI2L-H: Full CDS deletion; STOP in PpKAI2L-I (75); STOP in PpKAI2L-F (31); STOP in PpKAI2L-K (233); PpKAI2L-J: -($L^{22}KAH^{25}$); STOP in PpKAI2L-G (165); PpKAI2L-M: -($G^{160}DYI^{164}$) |

**Supplemental Table 3.** List of gene sequences used in this study

| Sequence ID | Genebank GI number | Phytozome number | Splicing variants (underlined variant has been used in this study) |
|---|---|---|---|
| PsRMS3 | GI:1839264 | | |
| AtD14 | GI:18396732 | | |
| AtKAI2 | GI:15235567 | | |
| BsRsbQ | GI:757754288 | | |
| PpKAI2L-A | | Pp3c2_19340 | Pp3c2_19340V3.1 ; Pp3c2_19340V3.2 |
| PpKAI2L-B | | Pp3c14_6110 | Pp3c14_6110V3.1 (5'UTR sequence) ; Pp3c14_6110V3.2 ; Pp3c14_6110V3.3 |
| PpKAI2L-C | | Pp3c25_5350 | Pp3c25_5350V3.1 ; Pp3c25_5350V3.2 ; Pp3c25_5350V3.3 ; Pp3c25_5350V3.4; Pp3c25_5350V3.5 ; Pp3c25_5350V3.6 |
| PpKAI2L-D | | Pp3c6_10610 | Pp3c6_10610V3.1 ; Pp3c6_10610V3.2 ; Pp3c6_10610V3.3 ; Pp3c6_10610V3.4 ; Pp3c6_10610V3.5 ; Pp3c6_10610V3.6 ; Pp3c6_10610V3.7 |
| PpKAI2L-E | | Pp3c5_16420 | Pp3c5_16420V3.1 ; Pp3c5_16420V3.2 ; Pp3c5_16420V3.3 ; Pp3c5_16420V3.4 ; Pp3c5_16420V3.5 ; Pp3c5_16420V3.6 ; Pp3c5_16420V3.7 |
| PpKAI2L-F | | Pp3c10_1460 | Pp3c10_1460V3.1 ; Pp3c10_1460V3.2 |
| PpKAI2L-G | | Pp3c4_4910 | Pp3c4_4910V3.1 ; Pp3c4_4910V3.2 |
| PpKAI2L-H | | Pp3c3_11730 | Pp3c3_11730V3.1 ; Pp3c3_11730V3.2 ; Pp3c3_11730V3.3 |
| PpKAI2L-I | | Pp3c12_8770 | Pp3c12_8770V3.1 ; Pp3c12_8770V3.2 |
| PpKAI2L-J | | Pp3c4_32050 | Pp3c4_32050V3.1 ; Pp3c4_32050V3.2 ; Pp3c4_32050V3.3 |
| PpKAI2L-K | | Pp3c1_18010 | Pp3c1_18010V3.1 ; Pp3c1_18010V3.2 ; Pp3c1_18010V3.3 ; |
| PpKAI2L-L | | Pp3c4_19700 | Pp3c4_19700V3.1 ; Pp3c4_19700V3.2 ; Pp3c4_19700V3.3 ; Pp3c4_19700V3.4 ; Pp3c4_19700V3.5 (for protein expression) |
| PpKAI2L-M | | Pp3c26_13220 | Pp3c26_13220V3.1 ; Pp3c26_13220V3.2 ; Pp3c26_13220V3.3 |
| PpMAX2 | | Pp3c17_1180 | |
| PpCCD7 | | Pp3c6_21550 | |
| PpCCD8 | | Pp3c6_21520 | |
| PpKUF1LA | | Pp3c2_34130 | |

(At) Arabidopsis thaliana ; (Bs) Bacillus subtilis ; (Ps) Pisum sativum ; (Pp) Physcomitrium patens
